# Template-based assembly of proteomic short reads for *de novo* antibody sequencing and repertoire profiling

**DOI:** 10.1101/2022.03.07.483237

**Authors:** Douwe Schulte, Weiwei Peng, Joost Snijder

## Abstract

Antibodies can target a vast molecular diversity of antigens. This is achieved by generating a complementary diversity of antibody sequences though somatic recombination and hypermutation. A full understanding of the antibody repertoire in health and disease therefore requires dedicated *de novo* sequencing methods. Next generation cDNA sequencing methods have laid the foundation of our current understanding of the antibody repertoire, but these methods share one major limitation in that they target the antibody-producing B-cells, rather than the functional secreted product in bodily fluids. Mass spectrometry-based methods offer an opportunity to bridge this gap between antibody repertoire profiling and bulk serological assays, as they can access antibody sequence information straight from the secreted polypeptide products. In a step to meeting the challenge of MS-based antibody sequencing, we present a fast and simple software tool (Stitch) to map proteomic short reads to user-defined templates with dedicated features for both monoclonal antibody sequencing and profiling of polyclonal antibody repertoires. We demonstrate the use of Stitch by fully reconstructing 2 monoclonal antibody sequences with >98% accuracy (including I/L assignment); sequencing a Fab from patient serum isolated by reversed-phase LC fractionation against a high background of homologous antibody sequences; sequencing antibody light chains from urine of multiple-myeloma patients; and profiling the IgG repertoire in sera from patients hospitalized with COVID-19. We demonstrate that Stitch assembles a comprehensive overview of the antibody sequences that are represented in the dataset and provides an important first step towards analyzing polyclonal antibodies and repertoire profiling.

## Introduction

Antibodies bind a wide variety of antigens with high affinity and specificity, playing a major role in the adaptive immune response to infections, but can also target self-antigens to mediate autoimmune diseases [1–4]. Antibodies can mediate immunity by blocking essential steps of a pathogen’s replication cycle (*e.g*. receptor binding and cell entry), triggering the complement system, or activating a specific cell-mediated immune response known as antibody-dependent cellular cytotoxicity. Antibodies elicited in response to infection may persist in circulation for several months and regenerate quickly by subsequent exposures to the same antigen through a memory B-cell response [5, 6]. All this has made antibodies into popular serological markers of pathogen exposure and vaccine efficacy, therapeutic leads for the treatment of cancer and infectious disease, and invaluable research tools for specific labelling and detection of molecular targets.

The large diversity of antigens that antibodies can target comes from a complementary diversity of available antibody sequences and compositions [2, 7–11]. Antibodies of most classes consist of a combination of two unique, paired, homologous polypeptides: the heavy chain and the light chain, each consisting of a series of the characteristic Immunoglobulin (Ig) domains. Both chains can be subdivided into a variable region, involved in antigen binding, and a constant region, which plays a structural role in oligomerization, complement activation and receptor-binding on immune cells. Disulfide bonds covalently link the heavy and light chains, and two copies of this covalent heterodimer are in turn disulfide-linked on the heavy chains to form the characteristic Y-shaped antibody (consisting of two heavy chains and two light chains). Up to four separate gene segments encode each chain by somatic recombination of the Variable, Diversity (only in heavy chain), Joining, and Constant segments, known as V-(D)-J-C recombination. Every unique B-cell clone can draw from many unique alleles for each segment (up to hundreds for the V-segment) creating in the order of 10^5^ possible unique V-(D)-J permutations. The number of possible unique pairings between the heavy and light chains further adds to the available variety of the fully assembled antibody.

The V-(D)-J segments collectively make up the variable domain of the heavy and light chains, each containing three so-called complementarity-determining regions (CDRs), which are directly involved in antigen binding and ultimately responsible for binding affinity and specificity [2, 9, 11]. CDR1 and CDR2 lie fully encoded within the V-segment, while CDR3 lies encoded in the V-(D)-J junction and is therefore inherently more variable. After initial activation of naïve B-cells, each with their specific V-(D)-J recombination, individual clones undergo a process of somatic hypermutation, in which additional sequence variation is introduced in the variable domain of the antibody, especially in the CDRs, resulting in affinity maturation of the coded antibody through a process of natural selection for the strongest antigen binders. This combination of somatic recombination, hypermutation and heavy-light chain pairing thus creates a vast repertoire of mature antibody sequences.

This large sequence diversity within and between individuals requires dedicated *de novo* sequencing of antibodies to uncover the structural basis of antigen binding and to map out the antibody repertoire in both health and disease [12, 13]. Established methods for *de novo* antibody sequencing rely on cloning and sequencing of the coding mRNAs from single B-cells, recovering up to hundreds of paired heavy and light chain sequences in a single study. While these methods have laid the foundation for our current understanding of the antibody repertoire, they share one major limitation in that sequencing requires recovery of the antibody-producing B-cells. While immature and memory B-cells do present antibodies on their surface, the major functional contribution of antibodies to adaptive immunity comes from the vast amounts that are secreted in blood and other bodily fluids. In other words, most antibodies are physically disconnected from their producing B-cell and there is no straightforward quantitative relation between serological test results (*e.g*. binding and neutralization titers) and antibody repertoires derived from single B-cell sequencing results. Expansion of memory B-cells into a secreting B-cell population, production and secretion levels of antibodies, and the lifetimes of both B-cells and secreted antibody product in bodily fluid may vary by orders of magnitude between unique B-cell clones. To address this caveat, methods to sequence and profile the functional antibody repertoire on the level of the secreted product are necessary.

Mass spectrometry-based methods are particularly powerful for direct proteomic sequencing and profiling of secreted antibody products. This is illustrated by several recent proteogenomics studies in which targeted single B-cell sequencing data is used to generate a custom database for a conventional proteomics-type LC-MS/MS-based database search to quantitatively profile the abundance of sequenced clones in serum [14–21]. These methods still rely on complementary cDNA sequencing of the antibody producing B-cell, unlike true *de novo* protein sequencing methods. Direct protein sequencing methods have focused especially on *de novo* sequencing of monoclonal antibodies, based on bottom-up analysis of digested peptides [22–31]. With the aid of specialized software packages like DiPS, Supernovo, or ALPS, full and accurate sequences of the heavy and light chains can be reconstructed with the MS/MS spectra of the digested peptides [28, 29, 31]. The use of multiple complementary proteases and novel hybrid fragmentation techniques provides large benefits in sequence coverage and accuracy in these methods [26]. The obtained sequences are complete and accurate enough to reverse-engineer functional synthetic recombinant antibodies, for instance of monoclonal antibodies from lost hybridoma cell lines. Recently we also demonstrated complete sequencing of a monoclonal antibody isolated from patient serum by reversed-phase LC fractionation and integrated bottom-up and top-down analysis [32]. Plasma proteomics methods to profile polyclonal IgG mixtures and other heterogeneous variant proteins based on *de novo* methods (SpotLight and LAX) have also recently been described [33–36].

Characterization of polyclonal mixtures and a move toward full profiling of the circulating antibody repertoire remain major outstanding challenges for MS-based antibody sequencing. In a step to meeting these challenges, we present a fast and simple software tool (Stitch) to map proteomic short reads to user-defined templates with dedicated features for both monoclonal antibody sequencing and profiling of polyclonal antibody repertoires. We demonstrate the use of Stitch by fully reconstructing 2 monoclonal antibody sequences with >98% accuracy (including I/L assignment); sequencing a Fab from patient serum isolated by reversed-phase LC fractionation against a high background of homologous antibody sequences; sequencing antibody light chains from urine of multiple-myeloma patients; and profiling the IgG repertoire in sera from patients hospitalized with COVID-19.

## Results

The experimental *de novo* antibody sequence reads obtained from a typical LC-MS/MS experiment are 5-40 amino acids in length. Although these reads are relatively short for completely *de novo* assembly, the rates of somatic hypermutation are typically low enough (1-10%) that the translated germline sequences contained in the IMGT are of sufficient homology to accurately place all peptide reads in the correct framework of the heavy and light chains. Based on this notion, we developed Stitch to perform template-based assembly of antibody-derived *de novo* sequence reads using local Smith-Waterman alignment [37]. Although the program can also perform this task on any user-defined set of templates, using plain FASTA sequences as input, we developed dedicated procedures for both mono- and polyclonal antibody sequencing using *de novo* reads from PEAKS as input [38]. With input from PEAKS the program can use metadata of individual reads as filtering criteria and determine weighted consensus sequences from overlapping reads, based on global and local quality scores, as well as MS1 peak area. As output Stitch generates an interactive HTML report that contains a quantitative overview of matched reads, alignment scores and a combined peak area for every template. In addition, it generates the final consensus sequences for all matched templates together with a sequence logo, depth of coverage profiles, and a detailed overview of all assembled reads in the context of their templates (see Figure 1). Finally, the output report also contains a complete overview of all reads assigned to the CDRs.

**Figure 1.**
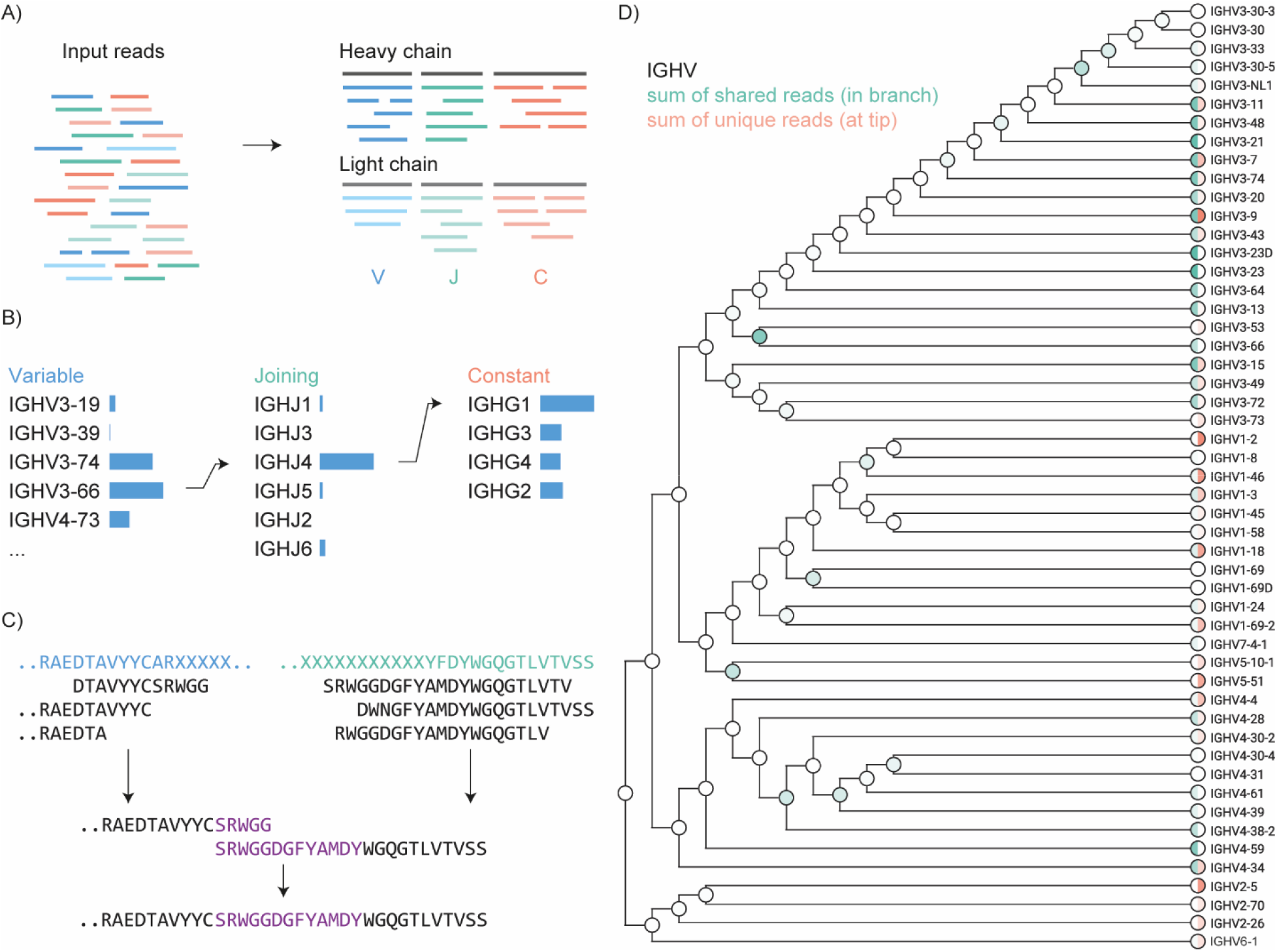
Schematic overview of Stitch. A) Input reads in PEAKS or FASTA format are matched to user-defined templates for V/J/C segments of the heavy and light chains by local Smith-Waterman alignment. B) For monoclonal antibodies the top-scoring segments can be recombined for a second template matching step on the full heavy and light chain sequences. C) Procedure to reconstruct CDRH3 looks for overlap between the overhanging reads extending the V- and J-segments. D) Shared and unique reads are placed at the corresponding position in a cladogram of homologous template sequences to provide a quantitative overview of the template matching with explicit consideration of ambiguity in the read placement. This example is whole IgG from a COVID-19 hospitalized patient, as further described in Figure 3.

In its most basic implementation, Stitch can simply match peptide reads to any homologous template in a user-defined database. Peptide reads are placed based on a user-defined cutoff score of the local alignment. When the database contains multiple templates, individual reads may match multiple entries with scores above this cutoff. This scenario is particularly relevant to antibody sequences as the multitude of available V/J/C alleles share a high degree of homology. The program can be set to place reads within all templates above the cutoff score or to place reads only on their single-highest scoring template. With this latter setting, reads with equal scores on multiple templates will be placed at all entries simultaneously. Reads with a single-highest scoring template are thereby defined as ‘unique’ for the program to track the total ‘unique’ alignment score and area of every template. Furthermore, Stitch explicitly considers the ambiguity of read placement across multiple homologous template sequences. A multiple sequence alignment is performed on each segment of the user-defined templates to generate a cladogram that represents the homology between the template sequences. Unique reads are placed at the tips of the branches, whereas shared reads are placed at the corresponding branching points of the tree (see Figure 1D). Stitch outputs the consensus sequence of every matched template based on all overlapping reads, accounting for frequency, global quality score and MS peak area with PEAKS data as input. The generated consensus sequence defaults to the template sequence in regions without coverage. Positions corresponding to I/L residues are defaulted to L in PEAKS data, as the two residues have identical masses and are therefore indistinguishable in most MS experiments. The consensus sequences in the output follow the matched template in these instances, changing the position to isoleucine when suggested by the template sequence.

Stitch allows templates to be defined in multiple separate groups, such that for antibody sequences we can sort peptide reads from heavy and light chains, and distinguish peptides from the V-, J- and C-segments of either chain. We have defined separate template databases for IGHV-IGHJ-IGHC, as well as IGLV-IGLJ-IGLC (with all kappa and lambda sequences combined in the same databases). The templates correspond to the germline sequences included in IMGT but filtered to create a reduced and non-redundant set of amino acid sequences (templates for human, mouse and rabbit antibodies are currently provided and the clean-up procedure to generate the non-redundant databases from additional species is included in the program). Templates for the D-segment are not taken from IMGT as they are too short and variable for any meaningful read-matching. In addition to the Ig segments, a separate decoy database for common contaminants of cell culture medium, plasma/serum, and proteomics sample preparation can be defined. The output report includes consensus sequences for all matched germline templates with annotation of the CDRs, as well as a quantitative overview of how each germline template is represented in the dataset by number of matched reads, alignment score, and combined peak area for the total set of matched reads, or the set of unique reads. Moreover, the program generates an aligned overview of all reads overlapping the CDRs (grouped by CDRH1, CDRH2, CDRH3, CDRL1, CDRL2, and CDRL3). The resulting report is a comprehensive overview of the antibody sequences that are represented in the dataset and provides an important first step towards analyzing polyclonal antibodies and repertoire profiling.

We have also built a dedicated procedure to reconstruct full monoclonal antibody sequences (see Figure 1B-C). Using the template-matching procedure described above, Stitch then selects the top-*N* scoring templates for each segment (with *N* = 1 for monoclonal antibodies) and recombines their consensus sequences into new V-J-C templates. As part of this recombination, CDRH3 is reconstructed by extending the V- and J-segments with the consensus sequences of overhanging reads to fill in the missing D-segment. The program then searches for identical sequences within the V- and J-overhanging regions to find the correct junction between the segments. A single gap is placed at the V-J junction if no overlap between the overhanging sequences can be found. The new recombined templates with reconstructed CDRH3 are then used for a second round of template matching to determine the final consensus sequences of the full heavy and light chains of the antibody. Stitch offers an option to use all non-selected germline templates as decoys in this second step to accommodate sequencing of monoclonal antibodies against a high background of homologous sequences, such as those collected from serum by LC fractionation or to cope with the presence of the multiple light or heavy chains which are often observed in hybridoma-derived antibodies [39].

To demonstrate the use of Stitch, we assembled *de novo* peptide reads to reconstruct the full heavy and light chain sequences of three different monoclonal antibodies. First, we used the human-mouse chimeric therapeutic antibody Herceptin (also known as Trastuzumab). Herceptin is composed of mouse CDR sequences placed within a human IgG1 framework and targets the Her2 receptor in treatment of a variety of cancers [26, 40, 41]. Second, we reconstructed the sequence of the anti-FLAG-M2 antibody, which is a mouse antibody targeting the DYKDDDDK epitope used to label and purify recombinant proteins [26, 42]. Alignment of the assembled output sequences reveals an overall accuracy of 98% and 99% (including I/L assignments) for Herceptin and anti-FLAG-M2, respectively (see Figure 2). A close-up view of the CDRH3 reconstructions demonstrates how the missing D-segment in the heavy chain is obtained through the two-step procedure described above (*i.e*. by extending the V- and J-segments with the consensus sequence of overlapping reads, searching for the V-J junction in the extended templates and performing a second round of template matching on the recombined V-J-C template, see Supplementary Figure S1). The third monoclonal antibody represents a more challenging case, as it is a Fab isolated directly from patient serum by reversed-phase LC-fractionation. Its sequence was originally determined by integrated use of both bottom-up and top-down LC-MS/MS data [32]. Using only the bottom-up data as input for Stitch we achieve a high accuracy of the reconstructed sequence, although we still see 2 errors (including 1 I/L misassignment) without the aid of complementary top-down LC-MS/MS data.

**Figure 2.**
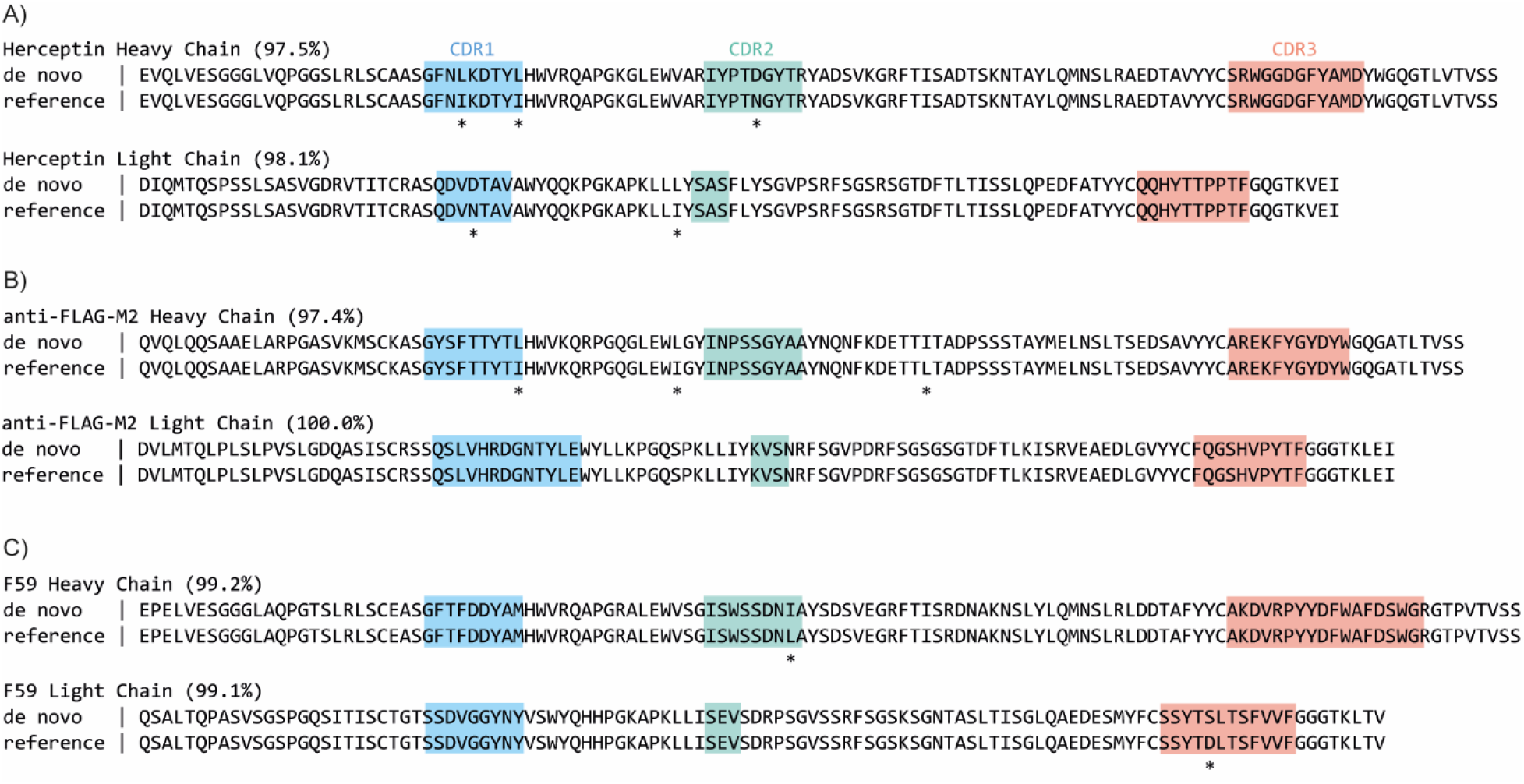
Stitch analysis of monoclonal antibodies. A) recombinant purified Herceptin. B) recombinant purified anti-FLAG-M2. C) Fab fragment F59 purified directly from patient serum using reversed-phase LC fractionation. CDRs are annotated, sequence conflicts highlighted by an asterisk (*) and the sequence identity listed in parentheses.

In addition to monoclonal antibodies, Stitch can also be used to assemble proteomic short reads of free light chains, such as those observed in multiple myeloma patients. Chamot-Rooke and colleagues recently reported proteomic sequencing of multiple myeloma light chains from patient urine, using an integrated bottom-up and top-down sequencing approach [43]. Here we used the bottom-up *de novo* sequencing reads from that study as a test case to reconstruct the light chains (see Supplementary Figure S2). The reconstructed sequences are in good agreement with those reported in the original study, with an average accuracy of 96% (ranging from 89% to 99%, including I/L assignments). Notably, in all instances of sequence conflicts, the coverage of the bottom-up data is limited or the corresponding alternative sequence is also present in reads within the dataset (but does not stand out in terms of quality score and MS peak area to dominate the consensus sequence). This further stresses the importance of the (depth of) coverage in the input data and highlights the added benefit of complementary top-down LC-MS/MS data for antibody sequencing.

To demonstrate the use of Stitch for profiling polyclonal antibody mixtures we generated a new dataset of *de novo* peptide reads from human serum. We obtained the total fraction of IgG, isolated by protein G affinity purification, from two individuals hospitalized with COVID-19. The purified IgG fractions were digested in parallel with four different proteases (trypsin, chymotrypsin, elastase and thermolysin), and analyzed by LC-MS/MS with a dual fragmentation scheme using both stepped HCD and EThcD fragmentation, with all obtained *de novo* sequence reads pooled into a single Stitch run. The analysis provides a quantitative overview of the IgG classes, use of kappa vs lambda light chains, and corresponding use of V-alleles across the total IgG repertoire of these patients (see Figure 3). For each of the two patient samples, we mapped 1276 and 1292 reads to IGHC, 697 and 837 reads to IGLC, 513 and 624 reads to IGHV, and 697 and 837 reads to IGLV. The profiles of both patients are remarkably similar, dominated by IgG1 with kappa light chains and drawing primarily from IGHV1/3 and IGLV1/3 alleles. Of the matched reads to the IGHV segment, 74 and 89 map to CDRH1, 65 and 92 to CDRH2, and 19 and 23 to CDRH2. Whereas the reads mapping to CDRH1/2 collectively span the full region, the CDRH3 reads are mostly limited to the first conserved AR/K residues following the preceding cysteine, or the conserved parts of the J-segment. Of the matched reads to the IGLV segment, 98 and 116 map to CDRL1, 136 and 182 to CDRL2, and 93 and 97 to CDRL3. The CDRL3 reads span a larger region compared to CDRH3, likely because the read assembly does not suffer from the missing D-segment. The Stitch analysis thus provides a quantitative overview of V-gene usage in polyclonal IgG mixtures, obtained straight from human serum samples, covering CDR1 and CDR2, but with notable limitations of CDR3.

**Figure 3.**
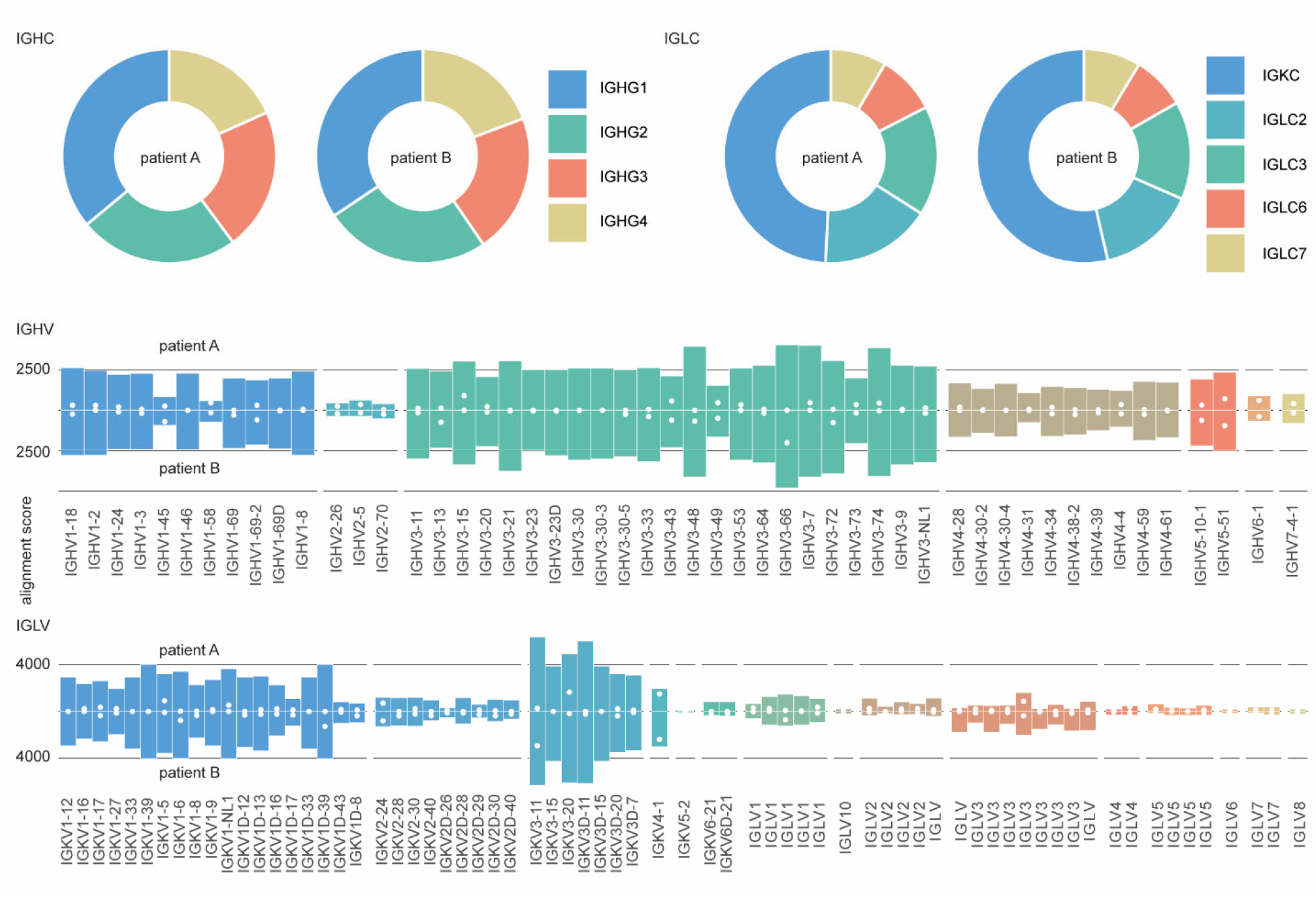
Repertoire profiling of protein G purified whole IgG from human serum. Profiles of IGHC, IGLC, IGHV and IGLV segments from two hospitalized COVID-19 patients as determined by Stitch. Shown are the total alignment scores of each matched template. The closed white circles in the IGHV/IGLV segments indicate score of the uniquely matched reads.

## Discussion

Stitch provides a quick and accessible way to assemble proteomic short reads against user-defined templates. It enables full reconstruction of monoclonal antibodies and free light chains, as well as profiling of polyclonal antibody mixtures. Given high-quality input reads, Stitch generates accurate consensus sequences, with remaining errors being fundamental to MS-based sequencing. These are errors related to deamidation (N to D) and assignment of isomeric I/L residues. Currently Stitch assigns I/L residues based on the matched template sequence, but this can potentially be further improved by considering experimental information, such as the cleavage specificity of chymotrypsin (cleaves only at L, not I) and use of diagnostic *w*-ions [44–47].

Whereas Stitch already explicitly considers both global and local quality scores of sequence reads, it does not yet provide integrated access to the underlying raw MS/MS data itself, which we aim to implement in the future. It is currently also limited to plain FASTA and PEAKS data as input reads, but we aim to adapt it to data formats from additional *de novo* sequencing software in the future. Current limitations regarding polyclonal antibody profiling will have to be solved with improved experimental approaches: obtaining longer sequence reads will reduce ambiguity in the correlation of sequence variants against the database of homologous templates, and top-down MS/MS of intact Fabs/antibodies or additional cross-linking MS workflows will have to elucidate the heavy-light chain pairings in the antibody mixture.

By enabling antibody sequencing and profiling from the purified secreted product, the development of Stitch contributes to an emerging new serology, in which bulk measures of antigen binding and neutralization can be directly related to the composition and sequence of a polyclonal antibody mixture. Direct MS-based sequencing and profiling of secreted antibodies thereby bridges the gap between bulk serological assays and B-cell sequencing approaches. These developments promise to provide a better understanding of antibody-mediated immunity in natural infection, vaccination and autoimmune disorders.

## Methods

### Monoclonal antibodies and COVID-19 serum IgG – sample preparation

Herceptin and anti-FLAG-M2 were obtained as described in reference [26], F59 was purified from patient serum as described in reference [32]. Convalescent serum from COVID-19 patients were obtained under the Radboud UMC Biobank protocol; IgG was purified with Protein G affinity resin (Millipore). Samples were denatured in 2% sodium deoxycholate (SDC), 200 mM Tris–HCl, 10 mM Tris(2-carboxyethyl)phosphine (TCEP), pH 8.0 at 95 °C for 10 min, followed by 30 min incubation at 37 °C for reduction. The samples were then alkylated by adding iodoacetic acid to a final concentration of 40 mM and incubated in the dark at room temperature for 45 min. For herceptin and anti-FLAG-M2 a 3 μg sample was then digested by one of the following proteases: trypsin (Promega), chymotrypsin (Roche), lysN (Thermo Fisher Scientific), lysC (FUJIFILM Wako Pure Chemical Corporation), gluC (Roche), aspN (Roche), aLP (Sigma-Aldrich), thermolysin (Promega), and elastase (Sigma-Aldrich) in a 1:50 ratio (w/w) in a total volume of 100 μL of 50 mM ammonium bicarbonate at 37 °C for 4 h. After digestion, SDC was removed by adding 2 μL of formic acid (FA) and centrifugation at 14 000g for 20 min. Following centrifugation, the supernatant containing the peptides was collected for desalting on a 30 μm Oasis HLB 96-well plate (Waters). The F59 monoclonal isolated from patient serum was digested in parallel by four proteases: trypsin, chymotrypsin, thermolysin and pepsin. Digestion with trypsin, chymotrypsin and thermolysin was done with 0.1 μg protease following the SDC protocol described above. For pepsin digestion, a urea buffer was added to a total volume of 80 μL, 2M Urea, 10 mM TCEP. Sample was denatured for 10 min at 95 °C followed by reduction for 20 min at 37 °C. Next, iodoacetic acid was added to a final concentration of 40 mM and incubated in the dark for 45 min at room temperature for alkylation of free cysteines. For pepsin digestion 1 M HCl was added to a final concentration of 0.04 M. Digestion was carried out overnight with 0.1 μg of protease, after which the entire digest was collected for desalting with the Oasis HLB 96-well plate. The Oasis HLB sorbent was activated with 100% acetonitrile and subsequently equilibrated with 10% formic acid in water. Next, peptides were bound to the sorbent, washed twice with 10% formic acid in water, and eluted with 100 μL of 50% acetonitrile/5% formic acid in water (v/v). The eluted peptides were vacuum-dried and reconstituted in 100 μL of 2% FA.

### LC-MS/MS

The digested peptides were separated by online reversed phase chromatography on an Agilent 1290 UHPLC coupled to a Thermo Scientific Orbitrap Fusion mass spectrometer. Peptides were separated using a Poroshell 120 EC-C18 2.7-Micron analytical column (ZORBAX Chromatographic Packing, Agilent) and a C18 PepMap 100 trap column (5 mm x 300 μm, 5 μm, Thermo Fisher Scientific). Samples were eluted over a 90 min gradient from 0 to 35% acetonitrile at a flow rate of 0.3 μL/min. Peptides were analyzed with a resolution setting of 60 000 in MS1. MS1 scans were obtained with a standard automatic gain control (AGC) target, a maximum injection time of 50 ms, and a scan range of 350–2000. The precursors were selected with a 3 m/z window and fragmented by stepped high-energy collision dissociation (HCD) as well as electron-transfer high-energy collision dissociation (EThcD). The stepped HCD fragmentation included steps of 25, 35, and 50% normalized collision energies (NCE). EThcD fragmentation was performed with calibrated charge-dependent electron-transfer dissociation (ETD) parameters and 27% NCE supplemental activation. For both fragmentation types, MS2 scans were acquired at a 30 000 resolution, a 4e5 AGC target, a 250 ms maximum injection time, and a scan range of 120–3500.

### Peptide sequencing from MS/MS spectra

MS/MS spectra were used to determine *de novo* peptide sequences using PEAKS Studio X (version 10.5). We used a tolerance of 20 ppm and 0.02 Da for MS1 and MS2, respectively. Carboxymethylation was set as fixed modification of cysteine, and variable modification of peptide N-termini and lysine. Oxidation of methionine and tryptophan, pyroglutamic acid modification of N-terminal glutamic acid and glutamine were set as additional variable modifications.

### Stitch analysis parameters

All Stitch analysis results are provided as supplementary data. The batch file parameters used for every analysis are outputted in the results file. Briefly, we typically use PEAKS ALC cutoff of ≥85, local alignment cutoff score of ≥8 and adjust these to the quality and complexity of the input data.

## Data and Code availability

The source code of Stitch is available on the Snijderlab GitHub page (https://github.com/snijderlab/stitch). All Stitch HTML results related to this study are provided as supplementary data. The raw data and PEAKS analyses unique to this study have been deposited in the ProteomeXchange Consortium via the PRIDE partner repository with the data set identifier PXD031941. The raw data of the monoclonal antibodies herceptin and anti-FLAG-M2 is available under identifier PXD023419. The raw data and PEAKS analyses of the multiple myeloma light chain dataset of Chamot-Rooke and colleagues is available under identifier PXD025884. The raw data of the serum-derived F59 monoclonal antibody is available at https://doi.org/10.6084/m9.figshare.13194005.

## Acknowledgments

We would like to thank Marien de Jonge and Dimitri Diavatopoulos (Radboud UMC) for sharing the COVID-19 patient sera. We would like to thank Keri T. Schmidt, Jade A.C. van der Hout, and Mariëlle E. Floor for extensive testing of Stitch. Further thanks to Bastiaan de Graaf and the rest of our colleagues in the Biomolecular Mass Spectrometry and Proteomics group at Utrecht University for support and helpful discussions. This research was funded by the Dutch Research Council NWO Gravitation 2013 BOO, Institute for Chemical Immunology (ICI; 024.002.009), and the Utrecht Molecular Immunology Hub.

## Notes

https://github.com/snijderlab/stitch

### Competing Interest Statement

The authors have declared no competing interest.

### Summary of Updates

Fixed an error in the analysis of Supplementary Figure S2 (original version omitted input from trypsin, lysC, and pepsin; all proteases are included now). The main text, Supplementary Data and Supplementary Information have been updated to reflect the corrected analysis. Some typos in the main text were also corrected and additional students have been listed in the Acknowledgements section to thank them for their tests of Stitch.

