## Supplementary material for "Template-based assembly of proteomic short reads for *de novo* antibody sequencing and repertoire profiling": F1_3107.html

Details F1\_3107

OverviewUndefined

### Read F1:3107

#### Sequence

DKKLVPRDCG

#### Sequence Length

10

#### Meta Information from PEAKS

##### Scan Identifier

F1:3107

##### Original Sequence (length=18)

D

K

K

L

V

P

R

D

C

+58.01

G

##### Posttranslational Modifications

Carboxymethyl

##### Source File

20191211\_F1\_Ag5\_peng0013\_SA\_Flag\_Asp\_N.raw

##### Fraction

1

##### Scan Feature

-

##### De Novo Score

96

##### Confidence score

96

##### Mass Charge Ratio

396.8729

##### Mass

1187.5969

##### Charge

3

##### Retention Time

16.86

##### Predicted Retention Time

-

##### Area

0

##### Parts Per Million

0

##### Fragmentation Mode

ETHCD
