## Supplementary material for "Template-based assembly of proteomic short reads for *de novo* antibody sequencing and repertoire profiling": F1_3137.html

Details F1\_3137

OverviewUndefined

### Read F1:3137

#### Sequence

DALHNHHTEKSLSHSPG

#### Sequence Length

17

#### Meta Information from PEAKS

##### Scan Identifier

F1:3137

##### Original Sequence (length=17)

D

A

L

H

N

H

H

T

E

K

S

L

S

H

S

P

G

##### Posttranslational Modifications

##### Source File

20191211\_F1\_Ag5\_peng0013\_SA\_Flag\_Asp\_N.raw

##### Fraction

1

##### Scan Feature

F1:9032

##### De Novo Score

96

##### Confidence score

96

##### Mass Charge Ratio

622.9662

##### Mass

1865.877

##### Charge

3

##### Retention Time

16.74

##### Predicted Retention Time

-

##### Area

28991000

##### Parts Per Million

0

##### Fragmentation Mode

ETHCD

##### Also found in scans

F1:3112 F1:3130 F1:3400 F1:3180
