## Supplementary material for "Template-based assembly of proteomic short reads for *de novo* antibody sequencing and repertoire profiling": F1_3432.html

Details F1\_3432

OverviewUndefined

### Read F1:3432

#### Sequence

DEYERHDSYV

#### Sequence Length

10

#### Meta Information from PEAKS

##### Scan Identifier

F1:3432

##### Original Sequence (length=10)

D

E

Y

E

R

H

D

S

Y

V

##### Posttranslational Modifications

##### Source File

20191211\_F1\_Ag5\_peng0013\_SA\_Flag\_Asp\_N.raw

##### Fraction

1

##### Scan Feature

F1:3194

##### De Novo Score

95

##### Confidence score

95

##### Mass Charge Ratio

438.1899

##### Mass

1311.5366

##### Charge

3

##### Retention Time

18.79

##### Predicted Retention Time

-

##### Area

14398000

##### Parts Per Million

8.7

##### Fragmentation Mode

ETHCD

##### Also found in scans

F1:3433
