## Supplementary material for "Template-based assembly of proteomic short reads for *de novo* antibody sequencing and repertoire profiling": F1_3451.html

Details F1\_3451

OverviewUndefined

### Read F1:3451

#### Sequence

DVVPPGLHNHHTEKSLSHSPG

#### Sequence Length

21

#### Meta Information from PEAKS

##### Scan Identifier

F1:3451

##### Original Sequence (length=29)

D

+58.01

V

V

P

P

G

L

H

N

H

H

T

E

K

S

L

S

H

S

P

G

##### Posttranslational Modifications

Carboxymethyl (KW X@N-term)

##### Source File

20191211\_F1\_Ag5\_peng0013\_SA\_Flag\_Asp\_N.raw

##### Fraction

1

##### Scan Feature

F1:7226

##### De Novo Score

90

##### Confidence score

90

##### Mass Charge Ratio

576.5377

##### Mass

2302.1089

##### Charge

4

##### Retention Time

18.76

##### Predicted Retention Time

-

##### Area

578590

##### Parts Per Million

5.6

##### Fragmentation Mode

ETHCD
