## Supplementary material for "Template-based assembly of proteomic short reads for *de novo* antibody sequencing and repertoire profiling": F1_3537.html

Details F1\_3537

OverviewUndefined

### Read F1:3537

#### Sequence

DVEVHTAQTQPR

#### Sequence Length

12

#### Meta Information from PEAKS

##### Scan Identifier

F1:3537

##### Original Sequence (length=12)

D

V

E

V

H

T

A

Q

T

Q

P

R

##### Posttranslational Modifications

##### Source File

20191211\_F1\_Ag5\_peng0013\_SA\_Flag\_Asp\_N.raw

##### Fraction

1

##### Scan Feature

F1:3962

##### De Novo Score

97

##### Confidence score

97

##### Mass Charge Ratio

460.9017

##### Mass

1379.6792

##### Charge

3

##### Retention Time

19.24

##### Predicted Retention Time

-

##### Area

406000

##### Parts Per Million

2.9

##### Fragmentation Mode

HCD

##### Also found in scans

F1:3534
