## Supplementary material for "Template-based assembly of proteomic short reads for *de novo* antibody sequencing and repertoire profiling": F1_3556.html

Details F1\_3556

OverviewUndefined

### Read F1:3556

#### Sequence

DEYERHGGSYTCEATHKTS

#### Sequence Length

19

#### Meta Information from PEAKS

##### Scan Identifier

F1:3556

##### Original Sequence (length=27)

D

E

Y

E

R

H

G

G

S

Y

T

C

+58.01

E

A

T

H

K

T

S

##### Posttranslational Modifications

Carboxymethyl

##### Source File

20191211\_F1\_Ag5\_peng0013\_SA\_Flag\_Asp\_N.raw

##### Fraction

1

##### Scan Feature

F1:6603

##### De Novo Score

92

##### Confidence score

92

##### Mass Charge Ratio

557.9833

##### Mass

2227.9075

##### Charge

4

##### Retention Time

19.35

##### Predicted Retention Time

-

##### Area

408390

##### Fragmentation Mode

HCD
