## Supplementary material for "Template-based assembly of proteomic short reads for *de novo* antibody sequencing and repertoire profiling": F1_3558.html

Details F1\_3558

OverviewUndefined

### Read F1:3558

#### Sequence

DEYERHWYV

#### Sequence Length

9

#### Meta Information from PEAKS

##### Scan Identifier

F1:3558

##### Original Sequence (length=17)

D

E

Y

E

R

H

W

+15.99

Y

V

##### Posttranslational Modifications

Oxidation (HW)

##### Source File

20191211\_F1\_Ag5\_peng0013\_SA\_Flag\_Asp\_N.raw

##### Fraction

1

##### Scan Feature

F1:10630

##### De Novo Score

97

##### Confidence score

97

##### Mass Charge Ratio

656.7808

##### Mass

1311.552

##### Charge

2

##### Retention Time

18.79

##### Predicted Retention Time

-

##### Area

9697900

##### Fragmentation Mode

ETHCD
