## Supplementary material for "Template-based assembly of proteomic short reads for *de novo* antibody sequencing and repertoire profiling": F1_3562.html

Details F1\_3562

OverviewUndefined

### Read F1:3562

#### Sequence

DQASLSCRSSQ

#### Sequence Length

11

#### Meta Information from PEAKS

##### Scan Identifier

F1:3562

##### Original Sequence (length=19)

D

Q

A

S

L

S

C

+58.01

R

S

S

Q

##### Posttranslational Modifications

Carboxymethyl

##### Source File

20191211\_F1\_Ag5\_peng0013\_SA\_Flag\_Asp\_N.raw

##### Fraction

1

##### Scan Feature

F1:8870

##### De Novo Score

95

##### Confidence score

95

##### Mass Charge Ratio

620.2672

##### Mass

1238.5198

##### Charge

2

##### Retention Time

19.38

##### Predicted Retention Time

-

##### Area

178010

##### Parts Per Million

0.1

##### Fragmentation Mode

HCD
