## Supplementary material for "Template-based assembly of proteomic short reads for *de novo* antibody sequencing and repertoire profiling": F1_3579.html

Details F1\_3579

OverviewUndefined

### Read F1:3579

#### Sequence

DEYERHNSY

#### Sequence Length

9

#### Meta Information from PEAKS

##### Scan Identifier

F1:3579

##### Original Sequence (length=9)

D

E

Y

E

R

H

N

S

Y

##### Posttranslational Modifications

##### Source File

20191211\_F1\_Ag5\_peng0013\_SA\_Flag\_Asp\_N.raw

##### Fraction

1

##### Scan Feature

F1:1917

##### De Novo Score

96

##### Confidence score

96

##### Mass Charge Ratio

404.8352

##### Mass

1211.4844

##### Charge

3

##### Retention Time

19.46

##### Predicted Retention Time

-

##### Area

42106

##### Fragmentation Mode

ETHCD
