## Supplementary material for "Template-based assembly of proteomic short reads for *de novo* antibody sequencing and repertoire profiling": F1_3580.html

Details F1\_3580

OverviewUndefined

### Read F1:3580

#### Sequence

DKKLPVRDCGCKP

#### Sequence Length

13

#### Meta Information from PEAKS

##### Scan Identifier

F1:3580

##### Original Sequence (length=29)

D

K

K

L

P

V

R

D

C

+58.01

G

C

+58.01

K

P

##### Posttranslational Modifications

Carboxymethyl

##### Source File

20191211\_F1\_Ag5\_peng0013\_SA\_Flag\_Asp\_N.raw

##### Fraction

1

##### Scan Feature

F1:5673

##### De Novo Score

91

##### Confidence score

91

##### Mass Charge Ratio

525.5935

##### Mass

1573.7593

##### Charge

3

##### Retention Time

19.46

##### Predicted Retention Time

-

##### Area

334210

##### Fragmentation Mode

ETHCD
