## Supplementary material for "Template-based assembly of proteomic short reads for *de novo* antibody sequencing and repertoire profiling": F1_3596.html

Details F1\_3596

OverviewUndefined

### Read F1:3596

#### Sequence

DDVEVHTAQTQPR

#### Sequence Length

13

#### Meta Information from PEAKS

##### Scan Identifier

F1:3596

##### Original Sequence (length=13)

D

D

V

E

V

H

T

A

Q

T

Q

P

R

##### Posttranslational Modifications

##### Source File

20191211\_F1\_Ag5\_peng0013\_SA\_Flag\_Asp\_N.raw

##### Fraction

1

##### Scan Feature

F1:4956

##### De Novo Score

98

##### Confidence score

98

##### Mass Charge Ratio

499.2432

##### Mass

1494.7063

##### Charge

3

##### Retention Time

19.56

##### Predicted Retention Time

-

##### Area

15663000

##### Parts Per Million

1

##### Fragmentation Mode

HCD

##### Also found in scans

F1:3626
