## Supplementary material for "Template-based assembly of proteomic short reads for *de novo* antibody sequencing and repertoire profiling": F1_3606.html

Details F1\_3606

OverviewUndefined

### Read F1:3606

#### Sequence

DVAHPASSTKV

#### Sequence Length

11

#### Meta Information from PEAKS

##### Scan Identifier

F1:3606

##### Original Sequence (length=11)

D

V

A

H

P

A

S

S

T

K

V

##### Posttranslational Modifications

##### Source File

20191211\_F1\_Ag5\_peng0013\_SA\_Flag\_Asp\_N.raw

##### Fraction

1

##### Scan Feature

F1:6551

##### De Novo Score

98

##### Confidence score

98

##### Mass Charge Ratio

556.2907

##### Mass

1110.5669

##### Charge

2

##### Retention Time

19.61

##### Predicted Retention Time

-

##### Area

265050

##### Parts Per Million

0

##### Fragmentation Mode

ETHCD
