## Supplementary material for "Template-based assembly of proteomic short reads for *de novo* antibody sequencing and repertoire profiling": F1_3608.html

Details F1\_3608

OverviewUndefined

### Read F1:3608

#### Sequence

DEYERHNSYT

#### Sequence Length

10

#### Meta Information from PEAKS

##### Scan Identifier

F1:3608

##### Original Sequence (length=10)

D

E

Y

E

R

H

N

S

Y

T

##### Posttranslational Modifications

##### Source File

20191211\_F1\_Ag5\_peng0013\_SA\_Flag\_Asp\_N.raw

##### Fraction

1

##### Scan Feature

F1:10643

##### De Novo Score

99

##### Confidence score

99

##### Mass Charge Ratio

657.2731

##### Mass

1312.532

##### Charge

2

##### Retention Time

19.61

##### Predicted Retention Time

-

##### Area

5560300

##### Fragmentation Mode

ETHCD
