## Supplementary material for "Template-based assembly of proteomic short reads for *de novo* antibody sequencing and repertoire profiling": F1_3616.html

Details F1\_3616

OverviewUndefined

### Read F1:3616

#### Sequence

DKGGNGGGLGSGGTVRSS

#### Sequence Length

18

#### Meta Information from PEAKS

##### Scan Identifier

F1:3616

##### Original Sequence (length=18)

D

K

G

G

N

G

G

G

L

G

S

G

G

T

V

R

S

S

##### Posttranslational Modifications

##### Source File

20191211\_F1\_Ag5\_peng0013\_SA\_Flag\_Asp\_N.raw

##### Fraction

1

##### Scan Feature

F1:5528

##### De Novo Score

94

##### Confidence score

94

##### Mass Charge Ratio

521.5928

##### Mass

1561.7444

##### Charge

3

##### Retention Time

19.72

##### Predicted Retention Time

-

##### Area

53587

##### Parts Per Million

7.8

##### Fragmentation Mode

ETHCD
