## Supplementary material for "Template-based assembly of proteomic short reads for *de novo* antibody sequencing and repertoire profiling": F1_3636.html

Details F1\_3636

OverviewUndefined

### Read F1:3636

#### Sequence

DALQQAKE

#### Sequence Length

8

#### Meta Information from PEAKS

##### Scan Identifier

F1:3636

##### Original Sequence (length=8)

D

A

L

Q

Q

A

K

E

##### Posttranslational Modifications

##### Source File

20191211\_F1\_Ag5\_peng0013\_SA\_Flag\_Asp\_N.raw

##### Fraction

1

##### Scan Feature

F1:3683

##### De Novo Score

93

##### Confidence score

93

##### Mass Charge Ratio

451.7327

##### Mass

901.4505

##### Charge

2

##### Retention Time

19.77

##### Predicted Retention Time

-

##### Area

62696

##### Parts Per Million

0.3

##### Fragmentation Mode

HCD
