## Supplementary material for "Template-based assembly of proteomic short reads for *de novo* antibody sequencing and repertoire profiling": F1_3638.html

Details F1\_3638

OverviewUndefined

### Read F1:3638

#### Sequence

DQASLSCRS

#### Sequence Length

9

#### Meta Information from PEAKS

##### Scan Identifier

F1:3638

##### Original Sequence (length=17)

D

Q

A

S

L

S

C

+58.01

R

S

##### Posttranslational Modifications

Carboxymethyl

##### Source File

20191211\_F1\_Ag5\_peng0013\_SA\_Flag\_Asp\_N.raw

##### Fraction

1

##### Scan Feature

F1:5290

##### De Novo Score

92

##### Confidence score

92

##### Mass Charge Ratio

512.7221

##### Mass

1023.4291

##### Charge

2

##### Retention Time

19.77

##### Predicted Retention Time

-

##### Area

886400

##### Parts Per Million

0.5

##### Fragmentation Mode

ETHCD
