## Supplementary material for "Template-based assembly of proteomic short reads for *de novo* antibody sequencing and repertoire profiling": F1_3639.html

Details F1\_3639

OverviewUndefined

### Read F1:3639

#### Sequence

DLSGSPATLR

#### Sequence Length

10

#### Meta Information from PEAKS

##### Scan Identifier

F1:3639

##### Original Sequence (length=18)

D

+58.01

L

S

G

S

P

A

T

L

R

##### Posttranslational Modifications

Carboxymethyl (KW X@N-term)

##### Source File

20191211\_F1\_Ag5\_peng0013\_SA\_Flag\_Asp\_N.raw

##### Fraction

1

##### Scan Feature

F1:6013

##### De Novo Score

93

##### Confidence score

93

##### Mass Charge Ratio

537.7804

##### Mass

1073.5352

##### Charge

2

##### Retention Time

19.84

##### Predicted Retention Time

-

##### Area

1542400

##### Parts Per Million

10.3

##### Fragmentation Mode

HCD
