## Supplementary material for "Template-based assembly of proteomic short reads for *de novo* antibody sequencing and repertoire profiling": F1_3649.html

Details F1\_3649

OverviewUndefined

### Read F1:3649

#### Sequence

DLTKVNKECCHG

#### Sequence Length

12

#### Meta Information from PEAKS

##### Scan Identifier

F1:3649

##### Original Sequence (length=28)

D

L

T

K

V

N

K

E

C

+58.01

C

+58.01

H

G

##### Posttranslational Modifications

Carboxymethyl

##### Source File

20191211\_F1\_Ag5\_peng0013\_SA\_Flag\_Asp\_N.raw

##### Fraction

1

##### Scan Feature

F1:4667

##### De Novo Score

99

##### Confidence score

99

##### Mass Charge Ratio

488.2149

##### Mass

1461.6228

##### Charge

3

##### Retention Time

19.91

##### Predicted Retention Time

-

##### Area

754640

##### Parts Per Million

0.1

##### Fragmentation Mode

ETHCD

##### Also found in scans

F1:3654
