## Supplementary material for "Template-based assembly of proteomic short reads for *de novo* antibody sequencing and repertoire profiling": F1_3681.html

Details F1\_3681

OverviewUndefined

### Read F1:3681

#### Sequence

DAALKTVM

#### Sequence Length

8

#### Meta Information from PEAKS

##### Scan Identifier

F1:3681

##### Original Sequence (length=16)

D

A

A

L

K

T

V

M

+15.99

##### Posttranslational Modifications

Oxidation (M)

##### Source File

20191211\_F1\_Ag5\_peng0013\_SA\_Flag\_Asp\_N.raw

##### Fraction

1

##### Scan Feature

F1:3022

##### De Novo Score

92

##### Confidence score

92

##### Mass Charge Ratio

432.7287

##### Mass

863.4423

##### Charge

2

##### Retention Time

20.04

##### Predicted Retention Time

-

##### Area

228070

##### Parts Per Million

0.6

##### Fragmentation Mode

HCD
