## Supplementary material for "Template-based assembly of proteomic short reads for *de novo* antibody sequencing and repertoire profiling": F1_3685.html

Details F1\_3685

OverviewUndefined

### Read F1:3685

#### Sequence

DVKLELRRA

#### Sequence Length

9

#### Meta Information from PEAKS

##### Scan Identifier

F1:3685

##### Original Sequence (length=17)

D

+58.01

V

K

L

E

L

R

R

A

##### Posttranslational Modifications

Carboxymethyl (KW X@N-term)

##### Source File

20191211\_F1\_Ag5\_peng0013\_SA\_Flag\_Asp\_N.raw

##### Fraction

1

##### Scan Feature

F1:1197

##### De Novo Score

94

##### Confidence score

94

##### Mass Charge Ratio

386.5629

##### Mass

1156.6562

##### Charge

3

##### Retention Time

20.09

##### Predicted Retention Time

-

##### Area

165140

##### Parts Per Million

9.3

##### Fragmentation Mode

ETHCD
