## Supplementary material for "Template-based assembly of proteomic short reads for *de novo* antibody sequencing and repertoire profiling": F1_3690.html

Details F1\_3690

OverviewUndefined

### Read F1:3690

#### Sequence

DDVEVHTAQTQPRE

#### Sequence Length

14

#### Meta Information from PEAKS

##### Scan Identifier

F1:3690

##### Original Sequence (length=14)

D

D

V

E

V

H

T

A

Q

T

Q

P

R

E

##### Posttranslational Modifications

##### Source File

20191211\_F1\_Ag5\_peng0013\_SA\_Flag\_Asp\_N.raw

##### Fraction

1

##### Scan Feature

F1:15805

##### De Novo Score

96

##### Confidence score

96

##### Mass Charge Ratio

812.8812

##### Mass

1623.7488

##### Charge

2

##### Retention Time

20.09

##### Predicted Retention Time

-

##### Area

997760

##### Fragmentation Mode

HCD

##### Also found in scans

F1:3686
