## Supplementary material for "Template-based assembly of proteomic short reads for *de novo* antibody sequencing and repertoire profiling": F1_3698.html

Details F1\_3698

OverviewUndefined

### Read F1:3698

#### Sequence

DKGVLVPR

#### Sequence Length

8

#### Meta Information from PEAKS

##### Scan Identifier

F1:3698

##### Original Sequence (length=8)

D

K

G

V

L

V

P

R

##### Posttranslational Modifications

##### Source File

20191211\_F1\_Ag5\_peng0013\_SA\_Flag\_Asp\_N.raw

##### Fraction

1

##### Scan Feature

F1:3319

##### De Novo Score

92

##### Confidence score

92

##### Mass Charge Ratio

442.2715

##### Mass

882.5287

##### Charge

2

##### Retention Time

20.13

##### Predicted Retention Time

-

##### Area

54784

##### Fragmentation Mode

ETHCD
