## Supplementary material for "Template-based assembly of proteomic short reads for *de novo* antibody sequencing and repertoire profiling": F1_3720.html

Details F1\_3720

OverviewUndefined

### Read F1:3720

#### Sequence

DDVEVHTAQTQPREE

#### Sequence Length

15

#### Meta Information from PEAKS

##### Scan Identifier

F1:3720

##### Original Sequence (length=15)

D

D

V

E

V

H

T

A

Q

T

Q

P

R

E

E

##### Posttranslational Modifications

##### Source File

20191211\_F1\_Ag5\_peng0013\_SA\_Flag\_Asp\_N.raw

##### Fraction

1

##### Scan Feature

F1:7522

##### De Novo Score

91

##### Confidence score

91

##### Mass Charge Ratio

585.27

##### Mass

1752.7915

##### Charge

3

##### Retention Time

20.27

##### Predicted Retention Time

-

##### Area

17693

##### Fragmentation Mode

ETHCD
