## Supplementary material for "Template-based assembly of proteomic short reads for *de novo* antibody sequencing and repertoire profiling": F1_3743.html

Details F1\_3743

OverviewUndefined

### Read F1:3743

#### Sequence

DNQRVLVNTM

#### Sequence Length

10

#### Meta Information from PEAKS

##### Scan Identifier

F1:3743

##### Original Sequence (length=10)

D

N

Q

R

V

L

V

N

T

M

##### Posttranslational Modifications

##### Source File

20191211\_F1\_Ag5\_peng0013\_SA\_Flag\_Asp\_N.raw

##### Fraction

1

##### Scan Feature

F1:1638

##### De Novo Score

96

##### Confidence score

96

##### Mass Charge Ratio

397.1996

##### Mass

1188.592

##### Charge

3

##### Retention Time

20.37

##### Predicted Retention Time

-

##### Area

26548

##### Fragmentation Mode

ETHCD
