## Supplementary material for "Template-based assembly of proteomic short reads for *de novo* antibody sequencing and repertoire profiling": F1_3751.html

Details F1\_3751

OverviewUndefined

### Read F1:3751

#### Sequence

DNQRVLVNTK

#### Sequence Length

10

#### Meta Information from PEAKS

##### Scan Identifier

F1:3751

##### Original Sequence (length=10)

D

N

Q

R

V

L

V

N

T

K

##### Posttranslational Modifications

##### Source File

20191211\_F1\_Ag5\_peng0013\_SA\_Flag\_Asp\_N.raw

##### Fraction

1

##### Scan Feature

F1:1601

##### De Novo Score

98

##### Confidence score

98

##### Mass Charge Ratio

396.2229

##### Mass

1185.6465

##### Charge

3

##### Retention Time

20.47

##### Predicted Retention Time

-

##### Area

907250

##### Parts Per Million

0.4

##### Fragmentation Mode

HCD
