## Supplementary material for "Template-based assembly of proteomic short reads for *de novo* antibody sequencing and repertoire profiling": F1_3761.html

Details F1\_3761

OverviewUndefined

### Read F1:3761

#### Sequence

DYLPASSTKVDKKLVPR

#### Sequence Length

17

#### Meta Information from PEAKS

##### Scan Identifier

F1:3761

##### Original Sequence (length=25)

D

+58.01

Y

L

P

A

S

S

T

K

V

D

K

K

L

V

P

R

##### Posttranslational Modifications

Carboxymethyl (KW X@N-term)

##### Source File

20191211\_F1\_Ag5\_peng0013\_SA\_Flag\_Asp\_N.raw

##### Fraction

1

##### Scan Feature

F1:10691

##### De Novo Score

96

##### Confidence score

96

##### Mass Charge Ratio

659.0413

##### Mass

1974.0786

##### Charge

3

##### Retention Time

20.52

##### Predicted Retention Time

-

##### Area

177430

##### Parts Per Million

11.9

##### Fragmentation Mode

HCD
