## Supplementary material for "Template-based assembly of proteomic short reads for *de novo* antibody sequencing and repertoire profiling": F1_3783.html

Details F1\_3783

OverviewUndefined

### Read F1:3783

#### Sequence

DSLHTSRNTYTA

#### Sequence Length

12

#### Meta Information from PEAKS

##### Scan Identifier

F1:3783

##### Original Sequence (length=12)

D

S

L

H

T

S

R

N

T

Y

T

A

##### Posttranslational Modifications

##### Source File

20191211\_F1\_Ag5\_peng0013\_SA\_Flag\_Asp\_N.raw

##### Fraction

1

##### Scan Feature

F1:11382

##### De Novo Score

96

##### Confidence score

96

##### Mass Charge Ratio

683.3228

##### Mass

1364.6321

##### Charge

2

##### Retention Time

20.42

##### Predicted Retention Time

-

##### Area

8659500

##### Fragmentation Mode

ETHCD
