## Supplementary material for "Template-based assembly of proteomic short reads for *de novo* antibody sequencing and repertoire profiling": F1_3793.html

Details F1\_3793

OverviewUndefined

### Read F1:3793

#### Sequence

DTAAHPASSTKV

#### Sequence Length

12

#### Meta Information from PEAKS

##### Scan Identifier

F1:3793

##### Original Sequence (length=12)

D

T

A

A

H

P

A

S

S

T

K

V

##### Posttranslational Modifications

##### Source File

20191211\_F1\_Ag5\_peng0013\_SA\_Flag\_Asp\_N.raw

##### Fraction

1

##### Scan Feature

F1:7805

##### De Novo Score

92

##### Confidence score

92

##### Mass Charge Ratio

592.7991

##### Mass

1183.5833

##### Charge

2

##### Retention Time

20.71

##### Predicted Retention Time

-

##### Area

1435000

##### Parts Per Million

0.4

##### Fragmentation Mode

HCD
