## Supplementary material for "Template-based assembly of proteomic short reads for *de novo* antibody sequencing and repertoire profiling": F1_3807.html

Details F1\_3807

OverviewUndefined

### Read F1:3807

#### Sequence

DTFSHEGLHNHHTEKSLSHSPG

#### Sequence Length

22

#### Meta Information from PEAKS

##### Scan Identifier

F1:3807

##### Original Sequence (length=22)

D

T

F

S

H

E

G

L

H

N

H

H

T

E

K

S

L

S

H

S

P

G

##### Posttranslational Modifications

##### Source File

20191211\_F1\_Ag5\_peng0013\_SA\_Flag\_Asp\_N.raw

##### Fraction

1

##### Scan Feature

F1:8609

##### De Novo Score

91

##### Confidence score

91

##### Mass Charge Ratio

614.2861

##### Mass

2453.1108

##### Charge

4

##### Retention Time

20.78

##### Predicted Retention Time

-

##### Area

1826200

##### Parts Per Million

1.9

##### Fragmentation Mode

ETHCD
