## Supplementary material for "Template-based assembly of proteomic short reads for *de novo* antibody sequencing and repertoire profiling": F1_3832.html

Details F1\_3832

OverviewUndefined

### Read F1:3832

#### Sequence

DEYERHNSYTC

#### Sequence Length

11

#### Meta Information from PEAKS

##### Scan Identifier

F1:3832

##### Original Sequence (length=19)

D

E

Y

E

R

H

N

S

Y

T

C

+58.01

##### Posttranslational Modifications

Carboxymethyl

##### Source File

20191211\_F1\_Ag5\_peng0013\_SA\_Flag\_Asp\_N.raw

##### Fraction

1

##### Scan Feature

F1:4771

##### De Novo Score

99

##### Confidence score

99

##### Mass Charge Ratio

492.1901

##### Mass

1473.5466

##### Charge

3

##### Retention Time

20.93

##### Predicted Retention Time

-

##### Area

3246200

##### Parts Per Million

1.2

##### Fragmentation Mode

ETHCD

##### Also found in scans

F1:3843
