## Supplementary material for "Template-based assembly of proteomic short reads for *de novo* antibody sequencing and repertoire profiling": F1_3850.html

Details F1\_3850

OverviewUndefined

### Read F1:3850

#### Sequence

DPPKTSTSPLVKSFNRNQ

#### Sequence Length

18

#### Meta Information from PEAKS

##### Scan Identifier

F1:3850

##### Original Sequence (length=18)

D

P

P

K

T

S

T

S

P

L

V

K

S

F

N

R

N

Q

##### Posttranslational Modifications

##### Source File

20191211\_F1\_Ag5\_peng0013\_SA\_Flag\_Asp\_N.raw

##### Fraction

1

##### Scan Feature

F1:1889

##### De Novo Score

94

##### Confidence score

94

##### Mass Charge Ratio

404.019

##### Mass

2015.0435

##### Charge

5

##### Retention Time

21.04

##### Predicted Retention Time

-

##### Area

190290

##### Parts Per Million

7.6

##### Fragmentation Mode

ETHCD
