## Supplementary material for "Template-based assembly of proteomic short reads for *de novo* antibody sequencing and repertoire profiling": F1_3865.html

Details F1\_3865

OverviewUndefined

### Read F1:3865

#### Sequence

DQEVCKNYAEAK

#### Sequence Length

12

#### Meta Information from PEAKS

##### Scan Identifier

F1:3865

##### Original Sequence (length=20)

D

Q

E

V

C

+58.01

K

N

Y

A

E

A

K

##### Posttranslational Modifications

Carboxymethyl

##### Source File

20191211\_F1\_Ag5\_peng0013\_SA\_Flag\_Asp\_N.raw

##### Fraction

1

##### Scan Feature

F1:4614

##### De Novo Score

99

##### Confidence score

99

##### Mass Charge Ratio

485.8855

##### Mass

1454.6348

##### Charge

3

##### Retention Time

21.15

##### Predicted Retention Time

-

##### Area

595040

##### Fragmentation Mode

ETHCD

##### Also found in scans

F1:3870
