## Supplementary material for "Template-based assembly of proteomic short reads for *de novo* antibody sequencing and repertoire profiling": F1_3869.html

Details F1\_3869

OverviewUndefined

### Read F1:3869

#### Sequence

DAELSQMQTHVS

#### Sequence Length

12

#### Meta Information from PEAKS

##### Scan Identifier

F1:3869

##### Original Sequence (length=20)

D

A

E

L

S

Q

M

+15.99

Q

T

H

V

S

##### Posttranslational Modifications

Oxidation (M)

##### Source File

20191211\_F1\_Ag5\_peng0013\_SA\_Flag\_Asp\_N.raw

##### Fraction

1

##### Scan Feature

F1:11319

##### De Novo Score

97

##### Confidence score

97

##### Mass Charge Ratio

681.3041

##### Mass

1360.5928

##### Charge

2

##### Retention Time

21.15

##### Predicted Retention Time

-

##### Area

111950

##### Parts Per Million

0.6

##### Fragmentation Mode

ETHCD
