## Supplementary material for "Template-based assembly of proteomic short reads for *de novo* antibody sequencing and repertoire profiling": F1_3882.html

Details F1\_3882

OverviewUndefined

### Read F1:3882

#### Sequence

DEYERHDSYTC

#### Sequence Length

11

#### Meta Information from PEAKS

##### Scan Identifier

F1:3882

##### Original Sequence (length=19)

D

E

Y

E

R

H

D

S

Y

T

C

+58.01

##### Posttranslational Modifications

Carboxymethyl

##### Source File

20191211\_F1\_Ag5\_peng0013\_SA\_Flag\_Asp\_N.raw

##### Fraction

1

##### Scan Feature

F1:4787

##### De Novo Score

93

##### Confidence score

93

##### Mass Charge Ratio

492.52

##### Mass

1474.5308

##### Charge

3

##### Retention Time

21.2

##### Predicted Retention Time

-

##### Area

96410

##### Parts Per Million

4.9

##### Fragmentation Mode

ETHCD
