## Supplementary material for "Template-based assembly of proteomic short reads for *de novo* antibody sequencing and repertoire profiling": F1_3889.html

Details F1\_3889

OverviewUndefined

### Read F1:3889

#### Sequence

DASTDQDSK

#### Sequence Length

9

#### Meta Information from PEAKS

##### Scan Identifier

F1:3889

##### Original Sequence (length=9)

D

A

S

T

D

Q

D

S

K

##### Posttranslational Modifications

##### Source File

20191211\_F1\_Ag5\_peng0013\_SA\_Flag\_Asp\_N.raw

##### Fraction

1

##### Scan Feature

F1:4562

##### De Novo Score

91

##### Confidence score

91

##### Mass Charge Ratio

483.7115

##### Mass

965.3937

##### Charge

2

##### Retention Time

21.2

##### Predicted Retention Time

-

##### Area

651630

##### Parts Per Million

15.3

##### Fragmentation Mode

ETHCD
