## Supplementary material for "Template-based assembly of proteomic short reads for *de novo* antibody sequencing and repertoire profiling": F1_3918.html

Details F1\_3918

OverviewUndefined

### Read F1:3918

#### Sequence

DQASLSSRSSQSLVHR

#### Sequence Length

16

#### Meta Information from PEAKS

##### Scan Identifier

F1:3918

##### Original Sequence (length=16)

D

Q

A

S

L

S

S

R

S

S

Q

S

L

V

H

R

##### Posttranslational Modifications

##### Source File

20191211\_F1\_Ag5\_peng0013\_SA\_Flag\_Asp\_N.raw

##### Fraction

1

##### Scan Feature

F1:7566

##### De Novo Score

96

##### Confidence score

96

##### Mass Charge Ratio

586.6345

##### Mass

1756.8816

##### Charge

3

##### Retention Time

21.49

##### Predicted Retention Time

-

##### Area

302710

##### Parts Per Million

0.1

##### Fragmentation Mode

ETHCD
