## Supplementary material for "Template-based assembly of proteomic short reads for *de novo* antibody sequencing and repertoire profiling": F1_3944.html

Details F1\_3944

OverviewUndefined

### Read F1:3944

#### Sequence

DSSVAHPASSTVK

#### Sequence Length

13

#### Meta Information from PEAKS

##### Scan Identifier

F1:3944

##### Original Sequence (length=13)

D

S

S

V

A

H

P

A

S

S

T

V

K

##### Posttranslational Modifications

##### Source File

20191211\_F1\_Ag5\_peng0013\_SA\_Flag\_Asp\_N.raw

##### Fraction

1

##### Scan Feature

F1:10111

##### De Novo Score

90

##### Confidence score

90

##### Mass Charge Ratio

643.3228

##### Mass

1284.6309

##### Charge

2

##### Retention Time

21.68

##### Predicted Retention Time

-

##### Area

1003700

##### Parts Per Million

0.1

##### Fragmentation Mode

HCD
