## Supplementary material for "Template-based assembly of proteomic short reads for *de novo* antibody sequencing and repertoire profiling": F1_3956.html

Details F1\_3956

OverviewUndefined

### Read F1:3956

#### Sequence

DPDSSTAYM

#### Sequence Length

9

#### Meta Information from PEAKS

##### Scan Identifier

F1:3956

##### Original Sequence (length=17)

D

P

D

S

S

T

A

Y

M

+15.99

##### Posttranslational Modifications

Oxidation (M)

##### Source File

20191211\_F1\_Ag5\_peng0013\_SA\_Flag\_Asp\_N.raw

##### Fraction

1

##### Scan Feature

F1:5020

##### De Novo Score

98

##### Confidence score

98

##### Mass Charge Ratio

501.6893

##### Mass

1001.3648

##### Charge

2

##### Retention Time

22.04

##### Predicted Retention Time

-

##### Area

783210

##### Fragmentation Mode

ETHCD
