## Supplementary material for "Template-based assembly of proteomic short reads for *de novo* antibody sequencing and repertoire profiling": F1_4008.html

Details F1\_4008

OverviewUndefined

### Read F1:4008

#### Sequence

DKHKTSTPSLVKSFNRNQ

#### Sequence Length

18

#### Meta Information from PEAKS

##### Scan Identifier

F1:4008

##### Original Sequence (length=26)

D

K

+58.01

H

K

T

S

T

P

S

L

V

K

S

F

N

R

N

Q

##### Posttranslational Modifications

Carboxymethyl (KW X@N-term)

##### Source File

20191211\_F1\_Ag5\_peng0013\_SA\_Flag\_Asp\_N.raw

##### Fraction

1

##### Scan Feature

F1:5988

##### De Novo Score

91

##### Confidence score

91

##### Mass Charge Ratio

537.0325

##### Mass

2144.0974

##### Charge

4

##### Retention Time

21.76

##### Predicted Retention Time

-

##### Area

8914400

##### Parts Per Million

1.6

##### Fragmentation Mode

ETHCD
