## Supplementary material for "Template-based assembly of proteomic short reads for *de novo* antibody sequencing and repertoire profiling": F1_4015.html

Details F1\_4015

OverviewUndefined

### Read F1:4015

#### Sequence

DAVRGSPAV

#### Sequence Length

9

#### Meta Information from PEAKS

##### Scan Identifier

F1:4015

##### Original Sequence (length=9)

D

A

V

R

G

S

P

A

V

##### Posttranslational Modifications

##### Source File

20191211\_F1\_Ag5\_peng0013\_SA\_Flag\_Asp\_N.raw

##### Fraction

1

##### Scan Feature

F1:3142

##### De Novo Score

98

##### Confidence score

98

##### Mass Charge Ratio

436.2355

##### Mass

870.4559

##### Charge

2

##### Retention Time

22.01

##### Predicted Retention Time

-

##### Area

161970

##### Parts Per Million

0.6

##### Fragmentation Mode

ETHCD
