## Supplementary material for "Template-based assembly of proteomic short reads for *de novo* antibody sequencing and repertoire profiling": F1_4035.html

Details F1\_4035

OverviewUndefined

### Read F1:4035

#### Sequence

DNQRVLVNTKA

#### Sequence Length

11

#### Meta Information from PEAKS

##### Scan Identifier

F1:4035

##### Original Sequence (length=11)

D

N

Q

R

V

L

V

N

T

K

A

##### Posttranslational Modifications

##### Source File

20191211\_F1\_Ag5\_peng0013\_SA\_Flag\_Asp\_N.raw

##### Fraction

1

##### Scan Feature

F1:2535

##### De Novo Score

92

##### Confidence score

92

##### Mass Charge Ratio

419.902

##### Mass

1256.6836

##### Charge

3

##### Retention Time

22.16

##### Predicted Retention Time

-

##### Area

600320

##### Parts Per Million

0.4

##### Fragmentation Mode

HCD
