## Supplementary material for "Template-based assembly of proteomic short reads for *de novo* antibody sequencing and repertoire profiling": F1_4039.html

Details F1\_4039

OverviewUndefined

### Read F1:4039

#### Sequence

DSWTDQDSK

#### Sequence Length

9

#### Meta Information from PEAKS

##### Scan Identifier

F1:4039

##### Original Sequence (length=9)

D

S

W

T

D

Q

D

S

K

##### Posttranslational Modifications

##### Source File

20191211\_F1\_Ag5\_peng0013\_SA\_Flag\_Asp\_N.raw

##### Fraction

1

##### Scan Feature

F1:6114

##### De Novo Score

99

##### Confidence score

99

##### Mass Charge Ratio

541.2256

##### Mass

1080.436

##### Charge

2

##### Retention Time

22.18

##### Predicted Retention Time

-

##### Area

2661000

##### Parts Per Million

0.7

##### Fragmentation Mode

ETHCD

##### Also found in scans

F1:4097
