## Supplementary material for "Template-based assembly of proteomic short reads for *de novo* antibody sequencing and repertoire profiling": F1_4094.html

Details F1\_4094

OverviewUndefined

### Read F1:4094

#### Sequence

DQASLSARSSQSLVHR

#### Sequence Length

16

#### Meta Information from PEAKS

##### Scan Identifier

F1:4094

##### Original Sequence (length=16)

D

Q

A

S

L

S

A

R

S

S

Q

S

L

V

H

R

##### Posttranslational Modifications

##### Source File

20191211\_F1\_Ag5\_peng0013\_SA\_Flag\_Asp\_N.raw

##### Fraction

1

##### Scan Feature

F1:7393

##### De Novo Score

97

##### Confidence score

97

##### Mass Charge Ratio

581.3034

##### Mass

1740.8867

##### Charge

3

##### Retention Time

22.46

##### Predicted Retention Time

-

##### Area

771310

##### Parts Per Million

1

##### Fragmentation Mode

HCD

##### Also found in scans

F1:4091
