## Supplementary material for "Template-based assembly of proteomic short reads for *de novo* antibody sequencing and repertoire profiling": F1_4119.html

Details F1\_4119

OverviewUndefined

### Read F1:4119

#### Sequence

DQASLSCRSSQSLVHRNGGG

#### Sequence Length

20

#### Meta Information from PEAKS

##### Scan Identifier

F1:4119

##### Original Sequence (length=28)

D

Q

A

S

L

S

C

+58.01

R

S

S

Q

S

L

V

H

R

N

G

G

G

##### Posttranslational Modifications

Carboxymethyl

##### Source File

20191211\_F1\_Ag5\_peng0013\_SA\_Flag\_Asp\_N.raw

##### Fraction

1

##### Scan Feature

F1:12223

##### De Novo Score

95

##### Confidence score

95

##### Mass Charge Ratio

706.3308

##### Mass

2115.9714

##### Charge

3

##### Retention Time

22.58

##### Predicted Retention Time

-

##### Area

1342100

##### Fragmentation Mode

ETHCD
