## Supplementary material for "Template-based assembly of proteomic short reads for *de novo* antibody sequencing and repertoire profiling": F1_4121.html

Details F1\_4121

OverviewUndefined

### Read F1:4121

#### Sequence

DGKEFKCRV

#### Sequence Length

9

#### Meta Information from PEAKS

##### Scan Identifier

F1:4121

##### Original Sequence (length=17)

D

G

K

E

F

K

C

+58.01

R

V

##### Posttranslational Modifications

Carboxymethyl

##### Source File

20191211\_F1\_Ag5\_peng0013\_SA\_Flag\_Asp\_N.raw

##### Fraction

1

##### Scan Feature

F1:1016

##### De Novo Score

97

##### Confidence score

97

##### Mass Charge Ratio

380.5226

##### Mass

1138.5439

##### Charge

3

##### Retention Time

22.62

##### Predicted Retention Time

-

##### Area

435040

##### Parts Per Million

1.7

##### Fragmentation Mode

ETHCD
