## Supplementary material for "Template-based assembly of proteomic short reads for *de novo* antibody sequencing and repertoire profiling": F1_4137.html

Details F1\_4137

OverviewUndefined

### Read F1:4137

#### Sequence

DPPKTSTSPLVKSFNRNEA

#### Sequence Length

19

#### Meta Information from PEAKS

##### Scan Identifier

F1:4137

##### Original Sequence (length=19)

D

P

P

K

T

S

T

S

P

L

V

K

S

F

N

R

N

E

A

##### Posttranslational Modifications

##### Source File

20191211\_F1\_Ag5\_peng0013\_SA\_Flag\_Asp\_N.raw

##### Fraction

1

##### Scan Feature

F1:5579

##### De Novo Score

95

##### Confidence score

95

##### Mass Charge Ratio

522.7767

##### Mass

2087.0647

##### Charge

4

##### Retention Time

22.64

##### Predicted Retention Time

-

##### Area

7613400

##### Parts Per Million

6.3

##### Fragmentation Mode

ETHCD

##### Also found in scans

F1:4108
