## Supplementary material for "Template-based assembly of proteomic short reads for *de novo* antibody sequencing and repertoire profiling": F1_4141.html

Details F1\_4141

OverviewUndefined

### Read F1:4141

#### Sequence

DEYERHNSYTCEA

#### Sequence Length

13

#### Meta Information from PEAKS

##### Scan Identifier

F1:4141

##### Original Sequence (length=21)

D

E

Y

E

R

H

N

S

Y

T

C

+58.01

E

A

##### Posttranslational Modifications

Carboxymethyl

##### Source File

20191211\_F1\_Ag5\_peng0013\_SA\_Flag\_Asp\_N.raw

##### Fraction

1

##### Scan Feature

F1:6631

##### De Novo Score

98

##### Confidence score

98

##### Mass Charge Ratio

558.8832

##### Mass

1673.6265

##### Charge

3

##### Retention Time

22.7

##### Predicted Retention Time

-

##### Area

183980

##### Parts Per Million

0.7

##### Fragmentation Mode

ETHCD
