## Supplementary material for "Template-based assembly of proteomic short reads for *de novo* antibody sequencing and repertoire profiling": F1_4148.html

Details F1\_4148

OverviewUndefined

### Read F1:4148

#### Sequence

DQASLSCRSSQSLVHRGG

#### Sequence Length

18

#### Meta Information from PEAKS

##### Scan Identifier

F1:4148

##### Original Sequence (length=26)

D

Q

A

S

L

S

C

+58.01

R

S

S

Q

S

L

V

H

R

G

G

##### Posttranslational Modifications

Carboxymethyl

##### Source File

20191211\_F1\_Ag5\_peng0013\_SA\_Flag\_Asp\_N.raw

##### Fraction

1

##### Scan Feature

F1:4641

##### De Novo Score

94

##### Confidence score

94

##### Mass Charge Ratio

487.2346

##### Mass

1944.9072

##### Charge

4

##### Retention Time

22.81

##### Predicted Retention Time

-

##### Area

10578000

##### Parts Per Million

1

##### Fragmentation Mode

ETHCD
