## Supplementary material for "Template-based assembly of proteomic short reads for *de novo* antibody sequencing and repertoire profiling": F1_4150.html

Details F1\_4150

OverviewUndefined

### Read F1:4150

#### Sequence

DNQRVLVGG

#### Sequence Length

9

#### Meta Information from PEAKS

##### Scan Identifier

F1:4150

##### Original Sequence (length=9)

D

N

Q

R

V

L

V

G

G

##### Posttranslational Modifications

##### Source File

20191211\_F1\_Ag5\_peng0013\_SA\_Flag\_Asp\_N.raw

##### Fraction

1

##### Scan Feature

F1:4462

##### De Novo Score

98

##### Confidence score

98

##### Mass Charge Ratio

479.2597

##### Mass

956.5039

##### Charge

2

##### Retention Time

22.79

##### Predicted Retention Time

-

##### Area

275050

##### Parts Per Million

1

##### Fragmentation Mode

ETHCD
