## Supplementary material for "Template-based assembly of proteomic short reads for *de novo* antibody sequencing and repertoire profiling": F1_4160.html

Details F1\_4160

OverviewUndefined

### Read F1:4160

#### Sequence

DQASLSCRSSQSLVHRGN

#### Sequence Length

18

#### Meta Information from PEAKS

##### Scan Identifier

F1:4160

##### Original Sequence (length=26)

D

Q

A

S

L

S

C

+58.01

R

S

S

Q

S

L

V

H

R

G

N

##### Posttranslational Modifications

Carboxymethyl

##### Source File

20191211\_F1\_Ag5\_peng0013\_SA\_Flag\_Asp\_N.raw

##### Fraction

1

##### Scan Feature

F1:10954

##### De Novo Score

97

##### Confidence score

97

##### Mass Charge Ratio

668.3173

##### Mass

2001.9287

##### Charge

3

##### Retention Time

22.85

##### Predicted Retention Time

-

##### Area

4486700

##### Parts Per Million

0.6

##### Fragmentation Mode

HCD
