## Supplementary material for "Template-based assembly of proteomic short reads for *de novo* antibody sequencing and repertoire profiling": F1_4186.html

Details F1\_4186

OverviewUndefined

### Read F1:4186

#### Sequence

DEYERHNSYTCEATHKTSTSPLVK

#### Sequence Length

24

#### Meta Information from PEAKS

##### Scan Identifier

F1:4186

##### Original Sequence (length=32)

D

E

Y

E

R

H

N

S

Y

T

C

+58.01

E

A

T

H

K

T

S

T

S

P

L

V

K

##### Posttranslational Modifications

Carboxymethyl

##### Source File

20191211\_F1\_Ag5\_peng0013\_SA\_Flag\_Asp\_N.raw

##### Fraction

1

##### Scan Feature

F1:12563

##### De Novo Score

94

##### Confidence score

94

##### Mass Charge Ratio

714.3295

##### Mass

2853.2874

##### Charge

4

##### Retention Time

22.92

##### Predicted Retention Time

-

##### Area

796210

##### Parts Per Million

0.5

##### Fragmentation Mode

HCD

##### Also found in scans

F4:4195 F4:4199 F1:4185
