## Supplementary material for "Template-based assembly of proteomic short reads for *de novo* antibody sequencing and repertoire profiling": F1_4192.html

Details F1\_4192

OverviewUndefined

### Read F1:4192

#### Sequence

DGLWGNASTA

#### Sequence Length

10

#### Meta Information from PEAKS

##### Scan Identifier

F1:4192

##### Original Sequence (length=18)

D

G

L

W

+58.01

G

N

A

S

T

A

##### Posttranslational Modifications

Carboxymethyl (KW X@N-term)

##### Source File

20191211\_F1\_Ag5\_peng0013\_SA\_Flag\_Asp\_N.raw

##### Fraction

1

##### Scan Feature

F1:5644

##### De Novo Score

93

##### Confidence score

93

##### Mass Charge Ratio

525.2322

##### Mass

1048.446

##### Charge

2

##### Retention Time

22.92

##### Predicted Retention Time

-

##### Area

189410

##### Parts Per Million

3.6

##### Fragmentation Mode

ETHCD
