## Supplementary material for "Template-based assembly of proteomic short reads for *de novo* antibody sequencing and repertoire profiling": F1_4234.html

Details F1\_4234

OverviewUndefined

### Read F1:4234

#### Sequence

DQASLSCRSSQSLVFLE

#### Sequence Length

17

#### Meta Information from PEAKS

##### Scan Identifier

F1:4234

##### Original Sequence (length=25)

D

Q

A

S

L

S

C

+58.01

R

S

S

Q

S

L

V

F

L

E

##### Posttranslational Modifications

Carboxymethyl

##### Source File

20191211\_F1\_Ag5\_peng0013\_SA\_Flag\_Asp\_N.raw

##### Fraction

1

##### Scan Feature

F1:10106

##### De Novo Score

96

##### Confidence score

96

##### Mass Charge Ratio

643.3063

##### Mass

1926.8992

##### Charge

3

##### Retention Time

23.24

##### Predicted Retention Time

-

##### Area

2444900

##### Fragmentation Mode

ETHCD
