## Supplementary material for "Template-based assembly of proteomic short reads for *de novo* antibody sequencing and repertoire profiling": F1_4237.html

Details F1\_4237

OverviewUndefined

### Read F1:4237

#### Sequence

DQASLSCRSSQSLVHRDH

#### Sequence Length

18

#### Meta Information from PEAKS

##### Scan Identifier

F1:4237

##### Original Sequence (length=34)

D

Q

A

S

L

S

C

+58.01

R

S

S

Q

S

L

V

H

R

D

H

+15.99

##### Posttranslational Modifications

Carboxymethyl; Oxidation (HW)

##### Source File

20191211\_F1\_Ag5\_peng0013\_SA\_Flag\_Asp\_N.raw

##### Fraction

1

##### Scan Feature

F1:11968

##### De Novo Score

96

##### Confidence score

96

##### Mass Charge Ratio

700.6554

##### Mass

2098.9451

##### Charge

3

##### Retention Time

23.28

##### Predicted Retention Time

-

##### Area

775240

##### Fragmentation Mode

HCD
