## Supplementary material for "Template-based assembly of proteomic short reads for *de novo* antibody sequencing and repertoire profiling": F1_4244.html

Details F1\_4244

OverviewUndefined

### Read F1:4244

#### Sequence

DKVRALEEANA

#### Sequence Length

11

#### Meta Information from PEAKS

##### Scan Identifier

F1:4244

##### Original Sequence (length=11)

D

K

V

R

A

L

E

E

A

N

A

##### Posttranslational Modifications

##### Source File

20191211\_F1\_Ag5\_peng0013\_SA\_Flag\_Asp\_N.raw

##### Fraction

1

##### Scan Feature

F1:1964

##### De Novo Score

98

##### Confidence score

98

##### Mass Charge Ratio

405.8828

##### Mass

1214.6255

##### Charge

3

##### Retention Time

23.35

##### Predicted Retention Time

-

##### Area

412650

##### Parts Per Million

0.9

##### Fragmentation Mode

ETHCD
