## Supplementary material for "Template-based assembly of proteomic short reads for *de novo* antibody sequencing and repertoire profiling": F1_4248.html

Details F1\_4248

OverviewUndefined

### Read F1:4248

#### Sequence

DKVRALEEAAN

#### Sequence Length

11

#### Meta Information from PEAKS

##### Scan Identifier

F1:4248

##### Original Sequence (length=11)

D

K

V

R

A

L

E

E

A

A

N

##### Posttranslational Modifications

##### Source File

20191211\_F1\_Ag5\_peng0013\_SA\_Flag\_Asp\_N.raw

##### Fraction

1

##### Scan Feature

F1:8347

##### De Novo Score

94

##### Confidence score

94

##### Mass Charge Ratio

608.3203

##### Mass

1214.6255

##### Charge

2

##### Retention Time

23.35

##### Predicted Retention Time

-

##### Area

492540

##### Parts Per Million

0.4

##### Fragmentation Mode

HCD
