## Supplementary material for "Template-based assembly of proteomic short reads for *de novo* antibody sequencing and repertoire profiling": F1_4266.html

Details F1\_4266

OverviewUndefined

### Read F1:4266

#### Sequence

DKHKTSTSPLVKSFNRNEA

#### Sequence Length

19

#### Meta Information from PEAKS

##### Scan Identifier

F1:4266

##### Original Sequence (length=27)

D

K

+58.01

H

K

T

S

T

S

P

L

V

K

S

F

N

R

N

E

A

##### Posttranslational Modifications

Carboxymethyl (KW X@N-term)

##### Source File

20191211\_F1\_Ag5\_peng0013\_SA\_Flag\_Asp\_N.raw

##### Fraction

1

##### Scan Feature

F1:3403

##### De Novo Score

94

##### Confidence score

94

##### Mass Charge Ratio

444.2311

##### Mass

2216.1184

##### Charge

5

##### Retention Time

23.37

##### Predicted Retention Time

-

##### Area

1741500

##### Parts Per Million

0.3

##### Fragmentation Mode

ETHCD

##### Also found in scans

F1:4284
