## Supplementary material for "Template-based assembly of proteomic short reads for *de novo* antibody sequencing and repertoire profiling": F1_4271.html

Details F1\_4271

OverviewUndefined

### Read F1:4271

#### Sequence

DGPSGSPATLR

#### Sequence Length

11

#### Meta Information from PEAKS

##### Scan Identifier

F1:4271

##### Original Sequence (length=11)

D

G

P

S

G

S

P

A

T

L

R

##### Posttranslational Modifications

##### Source File

20191211\_F1\_Ag5\_peng0013\_SA\_Flag\_Asp\_N.raw

##### Fraction

1

##### Scan Feature

F1:5772

##### De Novo Score

92

##### Confidence score

92

##### Mass Charge Ratio

529.2675

##### Mass

1056.52

##### Charge

2

##### Retention Time

23.42

##### Predicted Retention Time

-

##### Area

554760

##### Parts Per Million

0.3

##### Fragmentation Mode

HCD
