## Supplementary material for "Template-based assembly of proteomic short reads for *de novo* antibody sequencing and repertoire profiling": F1_4272.html

Details F1\_4272

OverviewUndefined

### Read F1:4272

#### Sequence

DKVQALEEANGG

#### Sequence Length

12

#### Meta Information from PEAKS

##### Scan Identifier

F1:4272

##### Original Sequence (length=12)

D

K

V

Q

A

L

E

E

A

N

G

G

##### Posttranslational Modifications

##### Source File

20191211\_F1\_Ag5\_peng0013\_SA\_Flag\_Asp\_N.raw

##### Fraction

1

##### Scan Feature

F1:8686

##### De Novo Score

98

##### Confidence score

98

##### Mass Charge Ratio

615.8015

##### Mass

1229.5889

##### Charge

2

##### Retention Time

23.51

##### Predicted Retention Time

-

##### Area

485810

##### Fragmentation Mode

ETHCD
