## Supplementary material for "Template-based assembly of proteomic short reads for *de novo* antibody sequencing and repertoire profiling": F1_4287.html

Details F1\_4287

OverviewUndefined

### Read F1:4287

#### Sequence

DLENKLAG

#### Sequence Length

8

#### Meta Information from PEAKS

##### Scan Identifier

F1:4287

##### Original Sequence (length=8)

D

L

E

N

K

L

A

G

##### Posttranslational Modifications

##### Source File

20191211\_F1\_Ag5\_peng0013\_SA\_Flag\_Asp\_N.raw

##### Fraction

1

##### Scan Feature

F1:2912

##### De Novo Score

97

##### Confidence score

97

##### Mass Charge Ratio

430.2297

##### Mass

858.4446

##### Charge

2

##### Retention Time

23.51

##### Predicted Retention Time

-

##### Area

215690

##### Parts Per Million

0.3

##### Fragmentation Mode

ETHCD
