## Supplementary material for "Template-based assembly of proteomic short reads for *de novo* antibody sequencing and repertoire profiling": F1_4302.html

Details F1\_4302

OverviewUndefined

### Read F1:4302

#### Sequence

DPPKTSTSPLVKSFNRNEC

#### Sequence Length

19

#### Meta Information from PEAKS

##### Scan Identifier

F1:4302

##### Original Sequence (length=27)

D

P

P

K

T

S

T

S

P

L

V

K

S

F

N

R

N

E

C

+58.01

##### Posttranslational Modifications

Carboxymethyl

##### Source File

20191211\_F1\_Ag5\_peng0013\_SA\_Flag\_Asp\_N.raw

##### Fraction

1

##### Scan Feature

F1:3149

##### De Novo Score

90

##### Confidence score

90

##### Mass Charge Ratio

436.4182

##### Mass

2177.0422

##### Charge

5

##### Retention Time

23.71

##### Predicted Retention Time

-

##### Area

146080

##### Parts Per Million

5.6

##### Fragmentation Mode

ETHCD
