## Supplementary material for "Template-based assembly of proteomic short reads for *de novo* antibody sequencing and repertoire profiling": F1_4325.html

Details F1\_4325

OverviewUndefined

### Read F1:4325

#### Sequence

DAELSQMQTHLS

#### Sequence Length

12

#### Meta Information from PEAKS

##### Scan Identifier

F1:4325

##### Original Sequence (length=20)

D

A

E

L

S

Q

M

+15.99

Q

T

H

L

S

##### Posttranslational Modifications

Oxidation (M)

##### Source File

20191211\_F1\_Ag5\_peng0013\_SA\_Flag\_Asp\_N.raw

##### Fraction

1

##### Scan Feature

F1:11583

##### De Novo Score

99

##### Confidence score

99

##### Mass Charge Ratio

688.3116

##### Mass

1374.6086

##### Charge

2

##### Retention Time

23.76

##### Predicted Retention Time

-

##### Area

679000

##### Parts Per Million

0

##### Fragmentation Mode

ETHCD
