## Supplementary material for "Template-based assembly of proteomic short reads for *de novo* antibody sequencing and repertoire profiling": F1_4339.html

Details F1\_4339

OverviewUndefined

### Read F1:4339

#### Sequence

DQASLSCRSSQSLVHR

#### Sequence Length

16

#### Meta Information from PEAKS

##### Scan Identifier

F1:4339

##### Original Sequence (length=24)

D

Q

A

S

L

S

C

+58.01

R

S

S

Q

S

L

V

H

R

##### Posttranslational Modifications

Carboxymethyl

##### Source File

20191211\_F1\_Ag5\_peng0013\_SA\_Flag\_Asp\_N.raw

##### Fraction

1

##### Scan Feature

F1:8473

##### De Novo Score

98

##### Confidence score

98

##### Mass Charge Ratio

611.2949

##### Mass

1830.8643

##### Charge

3

##### Retention Time

23.56

##### Predicted Retention Time

-

##### Area

102190000

##### Fragmentation Mode

HCD

##### Also found in scans

F1:4319 F1:4373 F1:4267 F1:4404
