## Supplementary material for "Template-based assembly of proteomic short reads for *de novo* antibody sequencing and repertoire profiling": F1_4366.html

Details F1\_4366

OverviewUndefined

### Read F1:4366

#### Sequence

DGQPAENYKDTQPLM

#### Sequence Length

15

#### Meta Information from PEAKS

##### Scan Identifier

F1:4366

##### Original Sequence (length=23)

D

G

Q

P

A

E

N

Y

K

D

T

Q

P

L

M

+15.99

##### Posttranslational Modifications

Oxidation (M)

##### Source File

20191211\_F1\_Ag5\_peng0013\_SA\_Flag\_Asp\_N.raw

##### Fraction

1

##### Scan Feature

F1:17160

##### De Novo Score

95

##### Confidence score

95

##### Mass Charge Ratio

861.887

##### Mass

1721.7566

##### Charge

2

##### Retention Time

23.99

##### Predicted Retention Time

-

##### Area

76262

##### Parts Per Million

1.6

##### Fragmentation Mode

ETHCD
