## Supplementary material for "Template-based assembly of proteomic short reads for *de novo* antibody sequencing and repertoire profiling": F1_4379.html

Details F1\_4379

OverviewUndefined

### Read F1:4379

#### Sequence

DGQPAENYKNTQPLM

#### Sequence Length

15

#### Meta Information from PEAKS

##### Scan Identifier

F1:4379

##### Original Sequence (length=23)

D

G

Q

P

A

E

N

Y

K

N

T

Q

P

L

M

+15.99

##### Posttranslational Modifications

Oxidation (M)

##### Source File

20191211\_F1\_Ag5\_peng0013\_SA\_Flag\_Asp\_N.raw

##### Fraction

1

##### Scan Feature

F1:17144

##### De Novo Score

97

##### Confidence score

97

##### Mass Charge Ratio

861.3932

##### Mass

1720.7727

##### Charge

2

##### Retention Time

24.3

##### Predicted Retention Time

-

##### Area

46399000

##### Fragmentation Mode

HCD
