## Supplementary material for "Template-based assembly of proteomic short reads for *de novo* antibody sequencing and repertoire profiling": F1_4381.html

Details F1\_4381

OverviewUndefined

### Read F1:4381

#### Sequence

DGQPAENYKNTQLW

#### Sequence Length

14

#### Meta Information from PEAKS

##### Scan Identifier

F1:4381

##### Original Sequence (length=22)

D

G

Q

P

A

E

N

Y

K

N

T

Q

L

W

+58.01

##### Posttranslational Modifications

Carboxymethyl (KW X@N-term)

##### Source File

20191211\_F1\_Ag5\_peng0013\_SA\_Flag\_Asp\_N.raw

##### Fraction

1

##### Scan Feature

F1:7172

##### De Novo Score

93

##### Confidence score

93

##### Mass Charge Ratio

574.5983

##### Mass

1720.7693

##### Charge

3

##### Retention Time

24.29

##### Predicted Retention Time

-

##### Area

5597000

##### Parts Per Million

2.2

##### Fragmentation Mode

ETHCD
