## Supplementary material for "Template-based assembly of proteomic short reads for *de novo* antibody sequencing and repertoire profiling": F1_4384.html

Details F1\_4384

OverviewUndefined

### Read F1:4384

#### Sequence

DLKVRAL

#### Sequence Length

7

#### Meta Information from PEAKS

##### Scan Identifier

F1:4384

##### Original Sequence (length=7)

D

L

K

V

R

A

L

##### Posttranslational Modifications

##### Source File

20191211\_F1\_Ag5\_peng0013\_SA\_Flag\_Asp\_N.raw

##### Fraction

1

##### Scan Feature

F1:2033

##### De Novo Score

91

##### Confidence score

91

##### Mass Charge Ratio

407.7613

##### Mass

813.5072

##### Charge

2

##### Retention Time

24.1

##### Predicted Retention Time

-

##### Area

95907

##### Parts Per Million

0.9

##### Fragmentation Mode

HCD
