## Supplementary material for "Template-based assembly of proteomic short reads for *de novo* antibody sequencing and repertoire profiling": F1_4418.html

Details F1\_4418

OverviewUndefined

### Read F1:4418

#### Sequence

DQADLSCRSSQSLVHR

#### Sequence Length

16

#### Meta Information from PEAKS

##### Scan Identifier

F1:4418

##### Original Sequence (length=24)

D

Q

A

D

L

S

C

+58.01

R

S

S

Q

S

L

V

H

R

##### Posttranslational Modifications

Carboxymethyl

##### Source File

20191211\_F1\_Ag5\_peng0013\_SA\_Flag\_Asp\_N.raw

##### Fraction

1

##### Scan Feature

F1:4108

##### De Novo Score

94

##### Confidence score

94

##### Mass Charge Ratio

465.7222

##### Mass

1858.8591

##### Charge

4

##### Retention Time

24.41

##### Predicted Retention Time

-

##### Area

399460

##### Parts Per Million

0.4

##### Fragmentation Mode

HCD
