## Supplementary material for "Template-based assembly of proteomic short reads for *de novo* antibody sequencing and repertoire profiling": F1_4422.html

Details F1\_4422

OverviewUndefined

### Read F1:4422

#### Sequence

DKHKTSTSPLVKSFNRNEC

#### Sequence Length

19

#### Meta Information from PEAKS

##### Scan Identifier

F1:4422

##### Original Sequence (length=35)

D

K

+58.01

H

K

T

S

T

S

P

L

V

K

S

F

N

R

N

E

C

+58.01

##### Posttranslational Modifications

Carboxymethyl (KW X@N-term); Carboxymethyl

##### Source File

20191211\_F1\_Ag5\_peng0013\_SA\_Flag\_Asp\_N.raw

##### Fraction

1

##### Scan Feature

F1:4006

##### De Novo Score

96

##### Confidence score

96

##### Mass Charge Ratio

462.2264

##### Mass

2306.0959

##### Charge

5

##### Retention Time

24.41

##### Predicted Retention Time

-

##### Area

5578800

##### Fragmentation Mode

ETHCD

##### Also found in scans

F1:4454
