## Supplementary material for "Template-based assembly of proteomic short reads for *de novo* antibody sequencing and repertoire profiling": F1_4423.html

Details F1\_4423

OverviewUndefined

### Read F1:4423

#### Sequence

DPPRTSTSPLVKSFNRNQ

#### Sequence Length

18

#### Meta Information from PEAKS

##### Scan Identifier

F1:4423

##### Original Sequence (length=18)

D

P

P

R

T

S

T

S

P

L

V

K

S

F

N

R

N

Q

##### Posttranslational Modifications

##### Source File

20191211\_F1\_Ag5\_peng0013\_SA\_Flag\_Asp\_N.raw

##### Fraction

1

##### Scan Feature

F1:5266

##### De Novo Score

91

##### Confidence score

91

##### Mass Charge Ratio

511.7697

##### Mass

2043.0498

##### Charge

4

##### Retention Time

24.3

##### Predicted Retention Time

-

##### Area

5674500

##### Fragmentation Mode

HCD
