## Supplementary material for "Template-based assembly of proteomic short reads for *de novo* antibody sequencing and repertoire profiling": F1_4447.html

Details F1\_4447

OverviewUndefined

### Read F1:4447

#### Sequence

DMTHKTSTSPLVKSFNNRQ

#### Sequence Length

19

#### Meta Information from PEAKS

##### Scan Identifier

F1:4447

##### Original Sequence (length=35)

D

+58.01

M

+15.99

T

H

K

T

S

T

S

P

L

V

K

S

F

N

N

R

Q

##### Posttranslational Modifications

Carboxymethyl (KW X@N-term); Oxidation (M)

##### Source File

20191211\_F1\_Ag5\_peng0013\_SA\_Flag\_Asp\_N.raw

##### Fraction

1

##### Scan Feature

F1:6942

##### De Novo Score

90

##### Confidence score

90

##### Mass Charge Ratio

567.0328

##### Mass

2264.0854

##### Charge

4

##### Retention Time

24.46

##### Predicted Retention Time

-

##### Area

3835100

##### Parts Per Million

7.4

##### Fragmentation Mode

HCD
