## Supplementary material for "Template-based assembly of proteomic short reads for *de novo* antibody sequencing and repertoire profiling": F1_4696.html

Details F1\_4696

OverviewUndefined

### Read F1:4696

#### Sequence

DSAVYYAAR

#### Sequence Length

9

#### Meta Information from PEAKS

##### Scan Identifier

F1:4696

##### Original Sequence (length=9)

D

S

A

V

Y

Y

A

A

R

##### Posttranslational Modifications

##### Source File

20191211\_F1\_Ag5\_peng0013\_SA\_Flag\_Asp\_N.raw

##### Fraction

1

##### Scan Feature

F1:5187

##### De Novo Score

99

##### Confidence score

99

##### Mass Charge Ratio

508.2462

##### Mass

1014.477

##### Charge

2

##### Retention Time

25.87

##### Predicted Retention Time

-

##### Area

5698100

##### Parts Per Million

0.8

##### Fragmentation Mode

HCD
