## Supplementary material for "Template-based assembly of proteomic short reads for *de novo* antibody sequencing and repertoire profiling": F1_4724.html

Details F1\_4724

OverviewUndefined

### Read F1:4724

#### Sequence

DPTTVTCNVAHPASSTKV

#### Sequence Length

18

#### Meta Information from PEAKS

##### Scan Identifier

F1:4724

##### Original Sequence (length=26)

D

P

T

T

V

T

C

+58.01

N

V

A

H

P

A

S

S

T

K

V

##### Posttranslational Modifications

Carboxymethyl

##### Source File

20191211\_F1\_Ag5\_peng0013\_SA\_Flag\_Asp\_N.raw

##### Fraction

1

##### Scan Feature

F1:9425

##### De Novo Score

98

##### Confidence score

98

##### Mass Charge Ratio

629.304

##### Mass

1884.8887

##### Charge

3

##### Retention Time

26.06

##### Predicted Retention Time

-

##### Area

4598400

##### Parts Per Million

0.7

##### Fragmentation Mode

ETHCD
