## Supplementary material for "Template-based assembly of proteomic short reads for *de novo* antibody sequencing and repertoire profiling": F1_4725.html

Details F1\_4725

OverviewUndefined

### Read F1:4725

#### Sequence

DKPLLKKAHCLSEVEH

#### Sequence Length

16

#### Meta Information from PEAKS

##### Scan Identifier

F1:4725

##### Original Sequence (length=24)

D

K

P

L

L

K

K

A

H

C

+58.01

L

S

E

V

E

H

##### Posttranslational Modifications

Carboxymethyl

##### Source File

20191211\_F1\_Ag5\_peng0013\_SA\_Flag\_Asp\_N.raw

##### Fraction

1

##### Scan Feature

F1:9757

##### De Novo Score

99

##### Confidence score

99

##### Mass Charge Ratio

635.6681

##### Mass

1903.9824

##### Charge

3

##### Retention Time

26.06

##### Predicted Retention Time

-

##### Area

774710

##### Parts Per Million

0

##### Fragmentation Mode

HCD

##### Also found in scans

F1:4745
