## Supplementary material for "Template-based assembly of proteomic short reads for *de novo* antibody sequencing and repertoire profiling": F1_4731.html

Details F1\_4731

OverviewUndefined

### Read F1:4731

#### Sequence

DMEATHKTSTSPLVKSFNRNEC

#### Sequence Length

22

#### Meta Information from PEAKS

##### Scan Identifier

F1:4731

##### Original Sequence (length=38)

D

M

+15.99

E

A

T

H

K

T

S

T

S

P

L

V

K

S

F

N

R

N

E

C

+58.01

##### Posttranslational Modifications

Oxidation (M); Carboxymethyl

##### Source File

20191211\_F1\_Ag5\_peng0013\_SA\_Flag\_Asp\_N.raw

##### Fraction

1

##### Scan Feature

F1:10097

##### De Novo Score

96

##### Confidence score

96

##### Mass Charge Ratio

643.0469

##### Mass

2568.1584

##### Charge

4

##### Retention Time

26.11

##### Predicted Retention Time

-

##### Area

1285700

##### Parts Per Million

0

##### Fragmentation Mode

ETHCD
