## Supplementary material for "Template-based assembly of proteomic short reads for *de novo* antibody sequencing and repertoire profiling": F1_4753.html

Details F1\_4753

OverviewUndefined

### Read F1:4753

#### Sequence

DPPLNSYTCEATHKTSTSPLVKSFDVWNC

#### Sequence Length

29

#### Meta Information from PEAKS

##### Scan Identifier

F1:4753

##### Original Sequence (length=45)

D

P

P

L

N

S

Y

T

C

+58.01

E

A

T

H

K

T

S

T

S

P

L

V

K

S

F

D

V

W

N

C

+58.01

##### Posttranslational Modifications

Carboxymethyl

##### Source File

20191211\_F1\_Ag5\_peng0013\_SA\_Flag\_Asp\_N.raw

##### Fraction

1

##### Scan Feature

-

##### De Novo Score

90

##### Confidence score

90

##### Mass Charge Ratio

672.1052

##### Mass

3355.5012

##### Charge

5

##### Retention Time

26.22

##### Predicted Retention Time

-

##### Area

0

##### Fragmentation Mode

HCD
