## Supplementary material for "Template-based assembly of proteomic short reads for *de novo* antibody sequencing and repertoire profiling": F1_4777.html

Details F1\_4777

OverviewUndefined

### Read F1:4777

#### Sequence

DGYMDSLHTSRNTYTA

#### Sequence Length

16

#### Meta Information from PEAKS

##### Scan Identifier

F1:4777

##### Original Sequence (length=24)

D

G

Y

M

+15.99

D

S

L

H

T

S

R

N

T

Y

T

A

##### Posttranslational Modifications

Oxidation (M)

##### Source File

20191211\_F1\_Ag5\_peng0013\_SA\_Flag\_Asp\_N.raw

##### Fraction

1

##### Scan Feature

F1:8721

##### De Novo Score

99

##### Confidence score

99

##### Mass Charge Ratio

616.6006

##### Mass

1846.7793

##### Charge

3

##### Retention Time

26.37

##### Predicted Retention Time

-

##### Area

3152500

##### Parts Per Million

0.3

##### Fragmentation Mode

ETHCD

##### Also found in scans

F1:4667
