## Supplementary material for "Template-based assembly of proteomic short reads for *de novo* antibody sequencing and repertoire profiling": F1_4799.html

Details F1\_4799

OverviewUndefined

### Read F1:4799

#### Sequence

DHQRSYTCEATHKTSTSPLVKSFNRNEC

#### Sequence Length

28

#### Meta Information from PEAKS

##### Scan Identifier

F1:4799

##### Original Sequence (length=44)

D

H

Q

R

S

Y

T

C

+58.01

E

A

T

H

K

T

S

T

S

P

L

V

K

S

F

N

R

N

E

C

+58.01

##### Posttranslational Modifications

Carboxymethyl

##### Source File

20191211\_F1\_Ag5\_peng0013\_SA\_Flag\_Asp\_N.raw

##### Fraction

1

##### Scan Feature

F1:22569

##### De Novo Score

92

##### Confidence score

92

##### Mass Charge Ratio

1119.1724

##### Mass

3354.4993

##### Charge

3

##### Retention Time

26.62

##### Predicted Retention Time

-

##### Area

66893000

##### Fragmentation Mode

ETHCD
