## Supplementary material for "Template-based assembly of proteomic short reads for *de novo* antibody sequencing and repertoire profiling": F1_4801.html

Details F1\_4801

OverviewUndefined

### Read F1:4801

#### Sequence

DWYKP

#### Sequence Length

5

#### Meta Information from PEAKS

##### Scan Identifier

F1:4801

##### Original Sequence (length=13)

D

W

+58.01

Y

K

P

##### Posttranslational Modifications

Carboxymethyl (KW X@N-term)

##### Source File

20191211\_F1\_Ag5\_peng0013\_SA\_Flag\_Asp\_N.raw

##### Fraction

1

##### Scan Feature

-

##### De Novo Score

98

##### Confidence score

98

##### Mass Charge Ratio

383.6741

##### Mass

765.3333

##### Charge

2

##### Retention Time

26.51

##### Predicted Retention Time

-

##### Area

0

##### Parts Per Million

0.5

##### Fragmentation Mode

ETHCD
