## Supplementary material for "Template-based assembly of proteomic short reads for *de novo* antibody sequencing and repertoire profiling": F1_4802.html

Details F1\_4802

OverviewUndefined

### Read F1:4802

#### Sequence

DNFYPK

#### Sequence Length

6

#### Meta Information from PEAKS

##### Scan Identifier

F1:4802

##### Original Sequence (length=6)

D

N

F

Y

P

K

##### Posttranslational Modifications

##### Source File

20191211\_F1\_Ag5\_peng0013\_SA\_Flag\_Asp\_N.raw

##### Fraction

1

##### Scan Feature

F1:1488

##### De Novo Score

96

##### Confidence score

96

##### Mass Charge Ratio

392.1876

##### Mass

782.3599

##### Charge

2

##### Retention Time

26.32

##### Predicted Retention Time

-

##### Area

31983000

##### Parts Per Million

0.9

##### Fragmentation Mode

ETHCD
