## Supplementary material for "Template-based assembly of proteomic short reads for *de novo* antibody sequencing and repertoire profiling": F1_4926.html

Details F1\_4926

OverviewUndefined

### Read F1:4926

#### Sequence

DLGGVFAQKLDGSERQ

#### Sequence Length

16

#### Meta Information from PEAKS

##### Scan Identifier

F1:4926

##### Original Sequence (length=16)

D

L

G

G

V

F

A

Q

K

L

D

G

S

E

R

Q

##### Posttranslational Modifications

##### Source File

20191211\_F1\_Ag5\_peng0013\_SA\_Flag\_Asp\_N.raw

##### Fraction

1

##### Scan Feature

F1:2937

##### De Novo Score

92

##### Confidence score

92

##### Mass Charge Ratio

430.7222

##### Mass

1718.8586

##### Charge

4

##### Retention Time

27.26

##### Predicted Retention Time

-

##### Area

4128200

##### Parts Per Million

0.7

##### Fragmentation Mode

HCD
