## Supplementary material for "Template-based assembly of proteomic short reads for *de novo* antibody sequencing and repertoire profiling": F1_4937.html

Details F1\_4937

OverviewUndefined

### Read F1:4937

#### Sequence

DLNVKMAKLDGSERQ

#### Sequence Length

15

#### Meta Information from PEAKS

##### Scan Identifier

F1:4937

##### Original Sequence (length=23)

D

L

N

V

K

M

+15.99

A

K

L

D

G

S

E

R

Q

##### Posttranslational Modifications

Oxidation (M)

##### Source File

20191211\_F1\_Ag5\_peng0013\_SA\_Flag\_Asp\_N.raw

##### Fraction

1

##### Scan Feature

F1:7147

##### De Novo Score

94

##### Confidence score

94

##### Mass Charge Ratio

573.9604

##### Mass

1718.8621

##### Charge

3

##### Retention Time

27.26

##### Predicted Retention Time

-

##### Area

10523000

##### Fragmentation Mode

HCD

##### Also found in scans

F1:5227
