## Supplementary material for "Template-based assembly of proteomic short reads for *de novo* antibody sequencing and repertoire profiling": F1_4967.html

Details F1\_4967

OverviewUndefined

### Read F1:4967

#### Sequence

DDVHKTSTSPLVKSFNRNEC

#### Sequence Length

20

#### Meta Information from PEAKS

##### Scan Identifier

F1:4967

##### Original Sequence (length=28)

D

D

V

H

K

T

S

T

S

P

L

V

K

S

F

N

R

N

E

C

+58.01

##### Posttranslational Modifications

Carboxymethyl

##### Source File

20191211\_F1\_Ag5\_peng0013\_SA\_Flag\_Asp\_N.raw

##### Fraction

1

##### Scan Feature

F1:7491

##### De Novo Score

93

##### Confidence score

93

##### Mass Charge Ratio

584.5297

##### Mass

2334.0908

##### Charge

4

##### Retention Time

27.57

##### Predicted Retention Time

-

##### Area

9828700

##### Fragmentation Mode

ETHCD
