## Supplementary material for "Template-based assembly of proteomic short reads for *de novo* antibody sequencing and repertoire profiling": F1_5025.html

Details F1\_5025

OverviewUndefined

### Read F1:5025

#### Sequence

DMRHKTSTSPLVKSFNRNEC

#### Sequence Length

20

#### Meta Information from PEAKS

##### Scan Identifier

F1:5025

##### Original Sequence (length=44)

D

+58.01

M

+15.99

R

H

K

T

S

T

S

P

L

V

K

S

F

N

R

N

E

C

+58.01

##### Posttranslational Modifications

Carboxymethyl (KW X@N-term); Oxidation (M); Carboxymethyl

##### Source File

20191211\_F1\_Ag5\_peng0013\_SA\_Flag\_Asp\_N.raw

##### Fraction

1

##### Scan Feature

F1:8937

##### De Novo Score

91

##### Confidence score

91

##### Mass Charge Ratio

621.293

##### Mass

2481.1375

##### Charge

4

##### Retention Time

27.72

##### Predicted Retention Time

-

##### Area

9472700

##### Parts Per Million

2.1

##### Fragmentation Mode

ETHCD
