## Supplementary material for "Template-based assembly of proteomic short reads for *de novo* antibody sequencing and repertoire profiling": F1_5054.html

Details F1\_5054

OverviewUndefined

### Read F1:5054

#### Sequence

DEYERHNSYTCEATHKTSTSPLVKSFNR

#### Sequence Length

28

#### Meta Information from PEAKS

##### Scan Identifier

F1:5054

##### Original Sequence (length=36)

D

E

Y

E

R

H

N

S

Y

T

C

+58.01

E

A

T

H

K

T

S

T

S

P

L

V

K

S

F

N

R

##### Posttranslational Modifications

Carboxymethyl

##### Source File

20191211\_F1\_Ag5\_peng0013\_SA\_Flag\_Asp\_N.raw

##### Fraction

1

##### Scan Feature

F1:11041

##### De Novo Score

96

##### Confidence score

96

##### Mass Charge Ratio

672.5143

##### Mass

3357.532

##### Charge

5

##### Retention Time

27.93

##### Predicted Retention Time

-

##### Area

11792000

##### Parts Per Million

1

##### Fragmentation Mode

HCD
