## Supplementary material for "Template-based assembly of proteomic short reads for *de novo* antibody sequencing and repertoire profiling": F1_5067.html

Details F1\_5067

OverviewUndefined

### Read F1:5067

#### Sequence

DEYERHNSYTCEATHKTSTKLPKSFHRG

#### Sequence Length

28

#### Meta Information from PEAKS

##### Scan Identifier

F1:5067

##### Original Sequence (length=52)

D

E

Y

E

R

H

N

S

Y

T

C

+58.01

E

A

T

H

K

T

S

T

K

+58.01

L

P

K

S

F

H

+15.99

R

G

##### Posttranslational Modifications

Carboxymethyl; Carboxymethyl (KW X@N-term); Oxidation (HW)

##### Source File

20191211\_F1\_Ag5\_peng0013\_SA\_Flag\_Asp\_N.raw

##### Fraction

1

##### Scan Feature

F1:17224

##### De Novo Score

90

##### Confidence score

90

##### Mass Charge Ratio

864.3981

##### Mass

3453.5642

##### Charge

4

##### Retention Time

27.93

##### Predicted Retention Time

-

##### Area

2346900

##### Fragmentation Mode

ETHCD
