## Supplementary material for "Template-based assembly of proteomic short reads for *de novo* antibody sequencing and repertoire profiling": F1_5112.html

Details F1\_5112

OverviewUndefined

### Read F1:5112

#### Sequence

DEYERHNSTYCEATHKTSTKLPKSFNRNEA

#### Sequence Length

30

#### Meta Information from PEAKS

##### Scan Identifier

F1:5112

##### Original Sequence (length=46)

D

E

Y

E

R

H

N

S

T

Y

C

+58.01

E

A

T

H

K

T

S

T

K

+58.01

L

P

K

S

F

N

R

N

E

A

##### Posttranslational Modifications

Carboxymethyl; Carboxymethyl (KW X@N-term)

##### Source File

20191211\_F1\_Ag5\_peng0013\_SA\_Flag\_Asp\_N.raw

##### Fraction

1

##### Scan Feature

F1:19038

##### De Novo Score

92

##### Confidence score

92

##### Mass Charge Ratio

918.9202

##### Mass

3671.6545

##### Charge

4

##### Retention Time

27.61

##### Predicted Retention Time

-

##### Area

61878000

##### Fragmentation Mode

ETHCD
