## Supplementary material for "Template-based assembly of proteomic short reads for *de novo* antibody sequencing and repertoire profiling": F1_5357.html

Details F1\_5357

OverviewUndefined

### Read F1:5357

#### Sequence

DGVLNSAMTDQDSK

#### Sequence Length

14

#### Meta Information from PEAKS

##### Scan Identifier

F1:5357

##### Original Sequence (length=22)

D

G

V

L

N

S

A

M

+15.99

T

D

Q

D

S

K

##### Posttranslational Modifications

Oxidation (M)

##### Source File

20191211\_F1\_Ag5\_peng0013\_SA\_Flag\_Asp\_N.raw

##### Fraction

1

##### Scan Feature

F1:13660

##### De Novo Score

94

##### Confidence score

94

##### Mass Charge Ratio

748.8287

##### Mass

1495.646

##### Charge

2

##### Retention Time

29.51

##### Predicted Retention Time

-

##### Area

3898800

##### Fragmentation Mode

HCD
