## Supplementary material for "Template-based assembly of proteomic short reads for *de novo* antibody sequencing and repertoire profiling": F1_5367.html

Details F1\_5367

OverviewUndefined

### Read F1:5367

#### Sequence

DSTYSTQELTLTK

#### Sequence Length

13

#### Meta Information from PEAKS

##### Scan Identifier

F1:5367

##### Original Sequence (length=13)

D

S

T

Y

S

T

Q

E

L

T

L

T

K

##### Posttranslational Modifications

##### Source File

20191211\_F1\_Ag5\_peng0013\_SA\_Flag\_Asp\_N.raw

##### Fraction

1

##### Scan Feature

F1:4870

##### De Novo Score

91

##### Confidence score

91

##### Mass Charge Ratio

496.2474

##### Mass

1485.72

##### Charge

3

##### Retention Time

29.66

##### Predicted Retention Time

-

##### Area

137130000

##### Parts Per Million

0.3

##### Fragmentation Mode

HCD
