## Supplementary material for "Template-based assembly of proteomic short reads for *de novo* antibody sequencing and repertoire profiling": F1_5389.html

Details F1\_5389

OverviewUndefined

### Read F1:5389

#### Sequence

DQDSKDSTYSMSSTLTLTK

#### Sequence Length

19

#### Meta Information from PEAKS

##### Scan Identifier

F1:5389

##### Original Sequence (length=27)

D

Q

D

S

K

D

S

T

Y

S

M

+15.99

S

S

T

L

T

L

T

K

##### Posttranslational Modifications

Oxidation (M)

##### Source File

20191211\_F1\_Ag5\_peng0013\_SA\_Flag\_Asp\_N.raw

##### Fraction

1

##### Scan Feature

F1:12313

##### De Novo Score

99

##### Confidence score

99

##### Mass Charge Ratio

708.6602

##### Mass

2122.9575

##### Charge

3

##### Retention Time

29.56

##### Predicted Retention Time

-

##### Area

88205000

##### Parts Per Million

0.5

##### Fragmentation Mode

ETHCD

##### Also found in scans

F1:5341 F1:5584
