## Supplementary material for "Template-based assembly of proteomic short reads for *de novo* antibody sequencing and repertoire profiling": F1_5406.html

Details F1\_5406

OverviewUndefined

### Read F1:5406

#### Sequence

DPSSSTAYM

#### Sequence Length

9

#### Meta Information from PEAKS

##### Scan Identifier

F1:5406

##### Original Sequence (length=9)

D

P

S

S

S

T

A

Y

M

##### Posttranslational Modifications

##### Source File

20191211\_F1\_Ag5\_peng0013\_SA\_Flag\_Asp\_N.raw

##### Fraction

1

##### Scan Feature

F1:4465

##### De Novo Score

98

##### Confidence score

98

##### Mass Charge Ratio

479.6953

##### Mass

957.3749

##### Charge

2

##### Retention Time

29.88

##### Predicted Retention Time

-

##### Area

3571600

##### Parts Per Million

1.1

##### Fragmentation Mode

ETHCD

##### Also found in scans

F1:3699 F1:3840
