## Supplementary material for "Template-based assembly of proteomic short reads for *de novo* antibody sequencing and repertoire profiling": F1_5411.html

Details F1\_5411

OverviewUndefined

### Read F1:5411

#### Sequence

DLARFRGTSDGYM

#### Sequence Length

13

#### Meta Information from PEAKS

##### Scan Identifier

F1:5411

##### Original Sequence (length=21)

D

L

A

R

F

R

G

T

S

D

G

Y

M

+15.99

##### Posttranslational Modifications

Oxidation (M)

##### Source File

20191211\_F1\_Ag5\_peng0013\_SA\_Flag\_Asp\_N.raw

##### Fraction

1

##### Scan Feature

F1:5024

##### De Novo Score

97

##### Confidence score

97

##### Mass Charge Ratio

502.2341

##### Mass

1503.6775

##### Charge

3

##### Retention Time

29.93

##### Predicted Retention Time

-

##### Area

2314700

##### Parts Per Million

2

##### Fragmentation Mode

ETHCD
