## Supplementary material for "Template-based assembly of proteomic short reads for *de novo* antibody sequencing and repertoire profiling": F1_5433.html

Details F1\_5433

OverviewUndefined

### Read F1:5433

#### Sequence

DSTYSQETLTLTK

#### Sequence Length

13

#### Meta Information from PEAKS

##### Scan Identifier

F1:5433

##### Original Sequence (length=13)

D

S

T

Y

S

Q

E

T

L

T

L

T

K

##### Posttranslational Modifications

##### Source File

20191211\_F1\_Ag5\_peng0013\_SA\_Flag\_Asp\_N.raw

##### Fraction

1

##### Scan Feature

F1:4870

##### De Novo Score

97

##### Confidence score

97

##### Mass Charge Ratio

496.2474

##### Mass

1485.72

##### Charge

3

##### Retention Time

29.66

##### Predicted Retention Time

-

##### Area

137130000

##### Parts Per Million

0.3

##### Fragmentation Mode

ETHCD

##### Also found in scans

F4:5399 F4:5458 F1:5505
