## Supplementary material for "Template-based assembly of proteomic short reads for *de novo* antibody sequencing and repertoire profiling": F1_5437.html

Details F1\_5437

OverviewUndefined

### Read F1:5437

#### Sequence

DLNVKWKLDGSERQ

#### Sequence Length

14

#### Meta Information from PEAKS

##### Scan Identifier

F1:5437

##### Original Sequence (length=22)

D

L

N

V

K

W

+15.99

K

L

D

G

S

E

R

Q

##### Posttranslational Modifications

Oxidation (HW)

##### Source File

20191211\_F1\_Ag5\_peng0013\_SA\_Flag\_Asp\_N.raw

##### Fraction

1

##### Scan Feature

F1:2784

##### De Novo Score

99

##### Confidence score

99

##### Mass Charge Ratio

426.724

##### Mass

1702.8638

##### Charge

4

##### Retention Time

30.03

##### Predicted Retention Time

-

##### Area

1515500

##### Parts Per Million

1.9

##### Fragmentation Mode

HCD

##### Also found in scans

F1:6333 F1:4439 F1:6378 F1:4451 F1:6551 F1:5567 F1:6167 F1:6209 F1:6992
