## Supplementary material for "Template-based assembly of proteomic short reads for *de novo* antibody sequencing and repertoire profiling": F1_5492.html

Details F1\_5492

OverviewUndefined

### Read F1:5492

#### Sequence

DTQPLM

#### Sequence Length

6

#### Meta Information from PEAKS

##### Scan Identifier

F1:5492

##### Original Sequence (length=6)

D

T

Q

P

L

M

##### Posttranslational Modifications

##### Source File

20191211\_F1\_Ag5\_peng0013\_SA\_Flag\_Asp\_N.raw

##### Fraction

1

##### Scan Feature

F1:12131

##### De Novo Score

91

##### Confidence score

91

##### Mass Charge Ratio

704.3278

##### Mass

703.3211

##### Charge

1

##### Retention Time

30.24

##### Predicted Retention Time

-

##### Area

150070

##### Fragmentation Mode

ETHCD
