## Supplementary material for "Template-based assembly of proteomic short reads for *de novo* antibody sequencing and repertoire profiling": F1_5589.html

Details F1\_5589

OverviewUndefined

### Read F1:5589

#### Sequence

DSTYEMSSTLTLTK

#### Sequence Length

14

#### Meta Information from PEAKS

##### Scan Identifier

F1:5589

##### Original Sequence (length=22)

D

S

T

Y

E

M

+15.99

S

S

T

L

T

L

T

K

##### Posttranslational Modifications

Oxidation (M)

##### Source File

20191211\_F1\_Ag5\_peng0013\_SA\_Flag\_Asp\_N.raw

##### Fraction

1

##### Scan Feature

-

##### De Novo Score

93

##### Confidence score

93

##### Mass Charge Ratio

531.5836

##### Mass

1591.7288

##### Charge

3

##### Retention Time

30.93

##### Predicted Retention Time

-

##### Area

0

##### Parts Per Million

0.1

##### Fragmentation Mode

ETHCD

##### Also found in scans

F1:5533 F1:5737
