## Supplementary material for "Template-based assembly of proteomic short reads for *de novo* antibody sequencing and repertoire profiling": F1_5630.html

Details F1\_5630

OverviewUndefined

### Read F1:5630

#### Sequence

DEYERHNSYTCEATHKTSTSPLVKSF

#### Sequence Length

26

#### Meta Information from PEAKS

##### Scan Identifier

F1:5630

##### Original Sequence (length=34)

D

E

Y

E

R

H

N

S

Y

T

C

+58.01

E

A

T

H

K

T

S

T

S

P

L

V

K

S

F

##### Posttranslational Modifications

Carboxymethyl

##### Source File

20191211\_F1\_Ag5\_peng0013\_SA\_Flag\_Asp\_N.raw

##### Fraction

1

##### Scan Feature

F1:14504

##### De Novo Score

96

##### Confidence score

96

##### Mass Charge Ratio

772.8541

##### Mass

3087.3879

##### Charge

4

##### Retention Time

31.22

##### Predicted Retention Time

-

##### Area

19144000

##### Fragmentation Mode

ETHCD
