## Supplementary material for "Template-based assembly of proteomic short reads for *de novo* antibody sequencing and repertoire profiling": F1_5769.html

Details F1\_5769

OverviewUndefined

### Read F1:5769

#### Sequence

DSTYSMSSPVTLTK

#### Sequence Length

14

#### Meta Information from PEAKS

##### Scan Identifier

F1:5769

##### Original Sequence (length=22)

D

S

T

Y

S

M

+15.99

S

S

P

V

T

L

T

K

##### Posttranslational Modifications

Oxidation (M)

##### Source File

20191211\_F1\_Ag5\_peng0013\_SA\_Flag\_Asp\_N.raw

##### Fraction

1

##### Scan Feature

-

##### De Novo Score

94

##### Confidence score

94

##### Mass Charge Ratio

511.5755

##### Mass

1531.7075

##### Charge

3

##### Retention Time

31.98

##### Predicted Retention Time

-

##### Area

0

##### Fragmentation Mode

ETHCD
