## Supplementary material for "Template-based assembly of proteomic short reads for *de novo* antibody sequencing and repertoire profiling": F1_5771.html

Details F1\_5771

OverviewUndefined

### Read F1:5771

#### Sequence

DSTVCCSSTLTLTK

#### Sequence Length

14

#### Meta Information from PEAKS

##### Scan Identifier

F1:5771

##### Original Sequence (length=30)

D

S

T

V

C

+58.01

C

+58.01

S

S

T

L

T

L

T

K

##### Posttranslational Modifications

Carboxymethyl

##### Source File

20191211\_F1\_Ag5\_peng0013\_SA\_Flag\_Asp\_N.raw

##### Fraction

1

##### Scan Feature

F1:5670

##### De Novo Score

94

##### Confidence score

94

##### Mass Charge Ratio

525.5661

##### Mass

1573.6851

##### Charge

3

##### Retention Time

32.11

##### Predicted Retention Time

-

##### Area

14132000

##### Fragmentation Mode

ETHCD
