## Supplementary material for "Template-based assembly of proteomic short reads for *de novo* antibody sequencing and repertoire profiling": F1_5776.html

Details F1\_5776

OverviewUndefined

### Read F1:5776

#### Sequence

DSYTCEATHKTSTSPLVKSLMMTAPC

#### Sequence Length

26

#### Meta Information from PEAKS

##### Scan Identifier

F1:5776

##### Original Sequence (length=50)

D

S

Y

T

C

+58.01

E

A

T

H

K

T

S

T

S

P

L

V

K

S

L

M

M

+15.99

T

A

P

C

+58.01

##### Posttranslational Modifications

Carboxymethyl; Oxidation (M)

##### Source File

20191211\_F1\_Ag5\_peng0013\_SA\_Flag\_Asp\_N.raw

##### Fraction

1

##### Scan Feature

F1:7613

##### De Novo Score

90

##### Confidence score

90

##### Mass Charge Ratio

587.6635

##### Mass

2933.2803

##### Charge

5

##### Retention Time

32.14

##### Predicted Retention Time

-

##### Area

33847000

##### Parts Per Million

0.2

##### Fragmentation Mode

HCD
