## Supplementary material for "Template-based assembly of proteomic short reads for *de novo* antibody sequencing and repertoire profiling": F1_5783.html

Details F1\_5783

OverviewUndefined

### Read F1:5783

#### Sequence

DSYTCEATHKTSTSPLVKSFNRNEC

#### Sequence Length

25

#### Meta Information from PEAKS

##### Scan Identifier

F1:5783

##### Original Sequence (length=41)

D

S

Y

T

C

+58.01

E

A

T

H

K

T

S

T

S

P

L

V

K

S

F

N

R

N

E

C

+58.01

##### Posttranslational Modifications

Carboxymethyl

##### Source File

20191211\_F1\_Ag5\_peng0013\_SA\_Flag\_Asp\_N.raw

##### Fraction

1

##### Scan Feature

F1:13182

##### De Novo Score

92

##### Confidence score

92

##### Mass Charge Ratio

734.3271

##### Mass

2933.2808

##### Charge

4

##### Retention Time

32.14

##### Predicted Retention Time

-

##### Area

152110000

##### Fragmentation Mode

ETHCD
