## Supplementary material for "Template-based assembly of proteomic short reads for *de novo* antibody sequencing and repertoire profiling": F1_5790.html

Details F1\_5790

OverviewUndefined

### Read F1:5790

#### Sequence

DSTYSMSSTLTLTK

#### Sequence Length

14

#### Meta Information from PEAKS

##### Scan Identifier

F1:5790

##### Original Sequence (length=22)

D

S

T

Y

S

M

+15.99

S

S

T

L

T

L

T

K

##### Posttranslational Modifications

Oxidation (M)

##### Source File

20191211\_F1\_Ag5\_peng0013\_SA\_Flag\_Asp\_N.raw

##### Fraction

1

##### Scan Feature

F1:14591

##### De Novo Score

99

##### Confidence score

99

##### Mass Charge Ratio

775.8666

##### Mass

1549.7183

##### Charge

2

##### Retention Time

32.06

##### Predicted Retention Time

-

##### Area

4966800000

##### Parts Per Million

0.2

##### Fragmentation Mode

HCD

##### Also found in scans

F1:14204 F1:15174 F1:7213 F1:12533 F1:13269 F1:11411 F1:11940 F1:12146 F1:6352 F1:10540 F4:5873 F4:6152 F1:12204 F1:5831 F4:6296 F1:6176 F1:6359 F1:5770 F1:14322 F1:7494 F4:14185 F4:5807 F1:9356 F4:5804
