## Supplementary material for "Template-based assembly of proteomic short reads for *de novo* antibody sequencing and repertoire profiling": F1_5802.html

Details F1\_5802

OverviewUndefined

### Read F1:5802

#### Sequence

DSYTCEATHKTSTSPLVKSFNNREC

#### Sequence Length

25

#### Meta Information from PEAKS

##### Scan Identifier

F1:5802

##### Original Sequence (length=41)

D

S

Y

T

C

+58.01

E

A

T

H

K

T

S

T

S

P

L

V

K

S

F

N

N

R

E

C

+58.01

##### Posttranslational Modifications

Carboxymethyl

##### Source File

20191211\_F1\_Ag5\_peng0013\_SA\_Flag\_Asp\_N.raw

##### Fraction

1

##### Scan Feature

F1:20957

##### De Novo Score

93

##### Confidence score

93

##### Mass Charge Ratio

978.7661

##### Mass

2933.2808

##### Charge

3

##### Retention Time

32.13

##### Predicted Retention Time

-

##### Area

40050000

##### Fragmentation Mode

HCD
