## Supplementary material for "Template-based assembly of proteomic short reads for *de novo* antibody sequencing and repertoire profiling": F1_5803.html

Details F1\_5803

OverviewUndefined

### Read F1:5803

#### Sequence

DQRLPCVE

#### Sequence Length

8

#### Meta Information from PEAKS

##### Scan Identifier

F1:5803

##### Original Sequence (length=16)

D

Q

R

L

P

C

+58.01

V

E

##### Posttranslational Modifications

Carboxymethyl

##### Source File

20191211\_F1\_Ag5\_peng0013\_SA\_Flag\_Asp\_N.raw

##### Fraction

1

##### Scan Feature

F1:5219

##### De Novo Score

92

##### Confidence score

92

##### Mass Charge Ratio

509.2366

##### Mass

1016.4597

##### Charge

2

##### Retention Time

32.11

##### Predicted Retention Time

-

##### Area

1540900

##### Fragmentation Mode

HCD
