## Supplementary material for "Template-based assembly of proteomic short reads for *de novo* antibody sequencing and repertoire profiling": F1_5804.html

Details F1\_5804

OverviewUndefined

### Read F1:5804

#### Sequence

DKVSLTSMLT

#### Sequence Length

10

#### Meta Information from PEAKS

##### Scan Identifier

F1:5804

##### Original Sequence (length=18)

D

K

V

S

L

T

S

M

+15.99

L

T

##### Posttranslational Modifications

Oxidation (M)

##### Source File

20191211\_F1\_Ag5\_peng0013\_SA\_Flag\_Asp\_N.raw

##### Fraction

1

##### Scan Feature

F1:6541

##### De Novo Score

96

##### Confidence score

96

##### Mass Charge Ratio

555.7893

##### Mass

1109.5637

##### Charge

2

##### Retention Time

32.22

##### Predicted Retention Time

-

##### Area

10156000

##### Parts Per Million

0.3

##### Fragmentation Mode

HCD
