## Supplementary material for "Template-based assembly of proteomic short reads for *de novo* antibody sequencing and repertoire profiling": F1_5874.html

Details F1\_5874

OverviewUndefined

### Read F1:5874

#### Sequence

DSTYMESDTLTLTK

#### Sequence Length

14

#### Meta Information from PEAKS

##### Scan Identifier

F1:5874

##### Original Sequence (length=22)

D

S

T

Y

M

+15.99

E

S

D

T

L

T

L

T

K

##### Posttranslational Modifications

Oxidation (M)

##### Source File

20191211\_F1\_Ag5\_peng0013\_SA\_Flag\_Asp\_N.raw

##### Fraction

1

##### Scan Feature

F1:6106

##### De Novo Score

92

##### Confidence score

92

##### Mass Charge Ratio

540.9153

##### Mass

1619.7236

##### Charge

3

##### Retention Time

32.59

##### Predicted Retention Time

-

##### Area

4108300

##### Parts Per Million

0.2

##### Fragmentation Mode

ETHCD
