## Supplementary material for "Template-based assembly of proteomic short reads for *de novo* antibody sequencing and repertoire profiling": F1_5886.html

Details F1\_5886

OverviewUndefined

### Read F1:5886

#### Sequence

DSMFMMSSTLTLTK

#### Sequence Length

14

#### Meta Information from PEAKS

##### Scan Identifier

F1:5886

##### Original Sequence (length=22)

D

S

M

F

M

M

+15.99

S

S

T

L

T

L

T

K

##### Posttranslational Modifications

Oxidation (M)

##### Source File

20191211\_F1\_Ag5\_peng0013\_SA\_Flag\_Asp\_N.raw

##### Fraction

1

##### Scan Feature

F1:15586

##### De Novo Score

91

##### Confidence score

91

##### Mass Charge Ratio

804.8698

##### Mass

1607.7246

##### Charge

2

##### Retention Time

32.59

##### Predicted Retention Time

-

##### Area

561520

##### Parts Per Million

0.3

##### Fragmentation Mode

ETHCD
