## Supplementary material for "Template-based assembly of proteomic short reads for *de novo* antibody sequencing and repertoire profiling": F1_6008.html

Details F1\_6008

OverviewUndefined

### Read F1:6008

#### Sequence

DLNVKWKLDGSERHS

#### Sequence Length

15

#### Meta Information from PEAKS

##### Scan Identifier

F1:6008

##### Original Sequence (length=15)

D

L

N

V

K

W

K

L

D

G

S

E

R

H

S

##### Posttranslational Modifications

##### Source File

20191211\_F1\_Ag5\_peng0013\_SA\_Flag\_Asp\_N.raw

##### Fraction

1

##### Scan Feature

F1:3514

##### De Novo Score

96

##### Confidence score

96

##### Mass Charge Ratio

446.7331

##### Mass

1782.9014

##### Charge

4

##### Retention Time

33.26

##### Predicted Retention Time

-

##### Area

2597300

##### Parts Per Million

1.1

##### Fragmentation Mode

HCD
