## Supplementary material for "Template-based assembly of proteomic short reads for *de novo* antibody sequencing and repertoire profiling": F1_6045.html

Details F1\_6045

OverviewUndefined

### Read F1:6045

#### Sequence

DLNVKWKLDGSERHA

#### Sequence Length

15

#### Meta Information from PEAKS

##### Scan Identifier

F1:6045

##### Original Sequence (length=23)

D

L

N

V

K

W

K

L

D

G

S

E

R

H

+15.99

A

##### Posttranslational Modifications

Oxidation (HW)

##### Source File

20191211\_F1\_Ag5\_peng0013\_SA\_Flag\_Asp\_N.raw

##### Fraction

1

##### Scan Feature

F1:7913

##### De Novo Score

92

##### Confidence score

92

##### Mass Charge Ratio

595.3076

##### Mass

1782.9014

##### Charge

3

##### Retention Time

33.26

##### Predicted Retention Time

-

##### Area

7796500

##### Fragmentation Mode

ETHCD
