## Supplementary material for "Template-based assembly of proteomic short reads for *de novo* antibody sequencing and repertoire profiling": F1_6076.html

Details F1\_6076

OverviewUndefined

### Read F1:6076

#### Sequence

DSTYSMSSELTLTK

#### Sequence Length

14

#### Meta Information from PEAKS

##### Scan Identifier

F1:6076

##### Original Sequence (length=22)

D

S

T

Y

S

M

+15.99

S

S

E

L

T

L

T

K

##### Posttranslational Modifications

Oxidation (M)

##### Source File

20191211\_F1\_Ag5\_peng0013\_SA\_Flag\_Asp\_N.raw

##### Fraction

1

##### Scan Feature

F1:5708

##### De Novo Score

91

##### Confidence score

91

##### Mass Charge Ratio

526.9114

##### Mass

1577.7131

##### Charge

3

##### Retention Time

33.68

##### Predicted Retention Time

-

##### Area

510280

##### Fragmentation Mode

HCD
