## Supplementary material for "Template-based assembly of proteomic short reads for *de novo* antibody sequencing and repertoire profiling": F1_6082.html

Details F1\_6082

OverviewUndefined

### Read F1:6082

#### Sequence

DLNVKWKLDGSERQN

#### Sequence Length

15

#### Meta Information from PEAKS

##### Scan Identifier

F1:6082

##### Original Sequence (length=15)

D

L

N

V

K

W

K

L

D

G

S

E

R

Q

N

##### Posttranslational Modifications

##### Source File

20191211\_F1\_Ag5\_peng0013\_SA\_Flag\_Asp\_N.raw

##### Fraction

1

##### Scan Feature

F1:8122

##### De Novo Score

98

##### Confidence score

98

##### Mass Charge Ratio

601.3112

##### Mass

1800.9119

##### Charge

3

##### Retention Time

33.62

##### Predicted Retention Time

-

##### Area

9121900

##### Fragmentation Mode

HCD

##### Also found in scans

F1:6073
