## Supplementary material for "Template-based assembly of proteomic short reads for *de novo* antibody sequencing and repertoire profiling": F1_6083.html

Details F1\_6083

OverviewUndefined

### Read F1:6083

#### Sequence

DSKDSTYSMSSTLTLTK

#### Sequence Length

17

#### Meta Information from PEAKS

##### Scan Identifier

F1:6083

##### Original Sequence (length=25)

D

S

K

D

S

T

Y

S

M

+15.99

S

S

T

L

T

L

T

K

##### Posttranslational Modifications

Oxidation (M)

##### Source File

20191211\_F1\_Ag5\_peng0013\_SA\_Flag\_Asp\_N.raw

##### Fraction

1

##### Scan Feature

-

##### De Novo Score

98

##### Confidence score

98

##### Mass Charge Ratio

627.6311

##### Mass

1879.8721

##### Charge

3

##### Retention Time

33.7

##### Predicted Retention Time

-

##### Area

0

##### Fragmentation Mode

ETHCD
