## Supplementary material for "Template-based assembly of proteomic short reads for *de novo* antibody sequencing and repertoire profiling": F1_6145.html

Details F1\_6145

OverviewUndefined

### Read F1:6145

#### Sequence

DLNVKWKLDGSWVQ

#### Sequence Length

14

#### Meta Information from PEAKS

##### Scan Identifier

F1:6145

##### Original Sequence (length=14)

D

L

N

V

K

W

K

L

D

G

S

W

V

Q

##### Posttranslational Modifications

##### Source File

20191211\_F1\_Ag5\_peng0013\_SA\_Flag\_Asp\_N.raw

##### Fraction

1

##### Scan Feature

F1:2619

##### De Novo Score

97

##### Confidence score

97

##### Mass Charge Ratio

422.7244

##### Mass

1686.8728

##### Charge

4

##### Retention Time

34.09

##### Predicted Retention Time

-

##### Area

157500000

##### Fragmentation Mode

HCD
