## Supplementary material for "Template-based assembly of proteomic short reads for *de novo* antibody sequencing and repertoire profiling": F1_6147.html

Details F1\_6147

OverviewUndefined

### Read F1:6147

#### Sequence

DSTYDMSSTLTLTK

#### Sequence Length

14

#### Meta Information from PEAKS

##### Scan Identifier

F1:6147

##### Original Sequence (length=22)

D

S

T

Y

D

M

+15.99

S

S

T

L

T

L

T

K

##### Posttranslational Modifications

Oxidation (M)

##### Source File

20191211\_F1\_Ag5\_peng0013\_SA\_Flag\_Asp\_N.raw

##### Fraction

1

##### Scan Feature

F1:15075

##### De Novo Score

99

##### Confidence score

99

##### Mass Charge Ratio

789.864

##### Mass

1577.7131

##### Charge

2

##### Retention Time

33.62

##### Predicted Retention Time

-

##### Area

49639000

##### Parts Per Million

0.2

##### Fragmentation Mode

HCD
