## Supplementary material for "Template-based assembly of proteomic short reads for *de novo* antibody sequencing and repertoire profiling": F1_6147_002.html

Details F1\_6147\_002

OverviewUndefined

### Read F1:6147\_002

#### Sequence

DSEYSMFELTLTK

#### Sequence Length

13

#### Meta Information from PEAKS

##### Scan Identifier

F1:6147

##### Original Sequence (length=21)

D

S

E

Y

S

M

+15.99

F

E

L

T

L

T

K

##### Posttranslational Modifications

Oxidation (M)

##### Source File

20191211\_F1\_Ag5\_peng0013\_SA\_Flag\_Asp\_N.raw

##### Fraction

1

##### Scan Feature

-

##### De Novo Score

96

##### Confidence score

96

##### Mass Charge Ratio

790.3657

##### Mass

1578.7124

##### Charge

2

##### Retention Time

34.07

##### Predicted Retention Time

-

##### Area

0

##### Parts Per Million

2.8

##### Fragmentation Mode

HCD
