## Supplementary material for "Template-based assembly of proteomic short reads for *de novo* antibody sequencing and repertoire profiling": F1_6154.html

Details F1\_6154

OverviewUndefined

### Read F1:6154

#### Sequence

DKVSLTAMLT

#### Sequence Length

10

#### Meta Information from PEAKS

##### Scan Identifier

F1:6154

##### Original Sequence (length=18)

D

K

V

S

L

T

A

M

+15.99

L

T

##### Posttranslational Modifications

Oxidation (M)

##### Source File

20191211\_F1\_Ag5\_peng0013\_SA\_Flag\_Asp\_N.raw

##### Fraction

1

##### Scan Feature

F1:6275

##### De Novo Score

96

##### Confidence score

96

##### Mass Charge Ratio

547.7914

##### Mass

1093.5688

##### Charge

2

##### Retention Time

34.09

##### Predicted Retention Time

-

##### Area

35623000

##### Fragmentation Mode

HCD
