## Supplementary material for "Template-based assembly of proteomic short reads for *de novo* antibody sequencing and repertoire profiling": F1_6155.html

Details F1\_6155

OverviewUndefined

### Read F1:6155

#### Sequence

DGVLNSSMTDQ

#### Sequence Length

11

#### Meta Information from PEAKS

##### Scan Identifier

F1:6155

##### Original Sequence (length=11)

D

G

V

L

N

S

S

M

T

D

Q

##### Posttranslational Modifications

##### Source File

20191211\_F1\_Ag5\_peng0013\_SA\_Flag\_Asp\_N.raw

##### Fraction

1

##### Scan Feature

F1:7470

##### De Novo Score

94

##### Confidence score

94

##### Mass Charge Ratio

583.751

##### Mass

1165.4922

##### Charge

2

##### Retention Time

34.09

##### Predicted Retention Time

-

##### Area

3562600

##### Fragmentation Mode

ETHCD
