## Supplementary material for "Template-based assembly of proteomic short reads for *de novo* antibody sequencing and repertoire profiling": F1_6166.html

Details F1\_6166

OverviewUndefined

### Read F1:6166

#### Sequence

DSTYSMSSTLTLTVS

#### Sequence Length

15

#### Meta Information from PEAKS

##### Scan Identifier

F1:6166

##### Original Sequence (length=23)

D

S

T

Y

S

M

+15.99

S

S

T

L

T

L

T

V

S

##### Posttranslational Modifications

Oxidation (M)

##### Source File

20191211\_F1\_Ag5\_peng0013\_SA\_Flag\_Asp\_N.raw

##### Fraction

1

##### Scan Feature

F1:15587

##### De Novo Score

96

##### Confidence score

96

##### Mass Charge Ratio

804.8701

##### Mass

1607.7236

##### Charge

2

##### Retention Time

34.45

##### Predicted Retention Time

-

##### Area

28392000

##### Parts Per Million

1.3

##### Fragmentation Mode

ETHCD
