## Supplementary material for "Template-based assembly of proteomic short reads for *de novo* antibody sequencing and repertoire profiling": F1_6202.html

Details F1\_6202

OverviewUndefined

### Read F1:6202

#### Sequence

DKDTCFSTEGPNLVTRCK

#### Sequence Length

18

#### Meta Information from PEAKS

##### Scan Identifier

F1:6202

##### Original Sequence (length=34)

D

K

D

T

C

+58.01

F

S

T

E

G

P

N

L

V

T

R

C

+58.01

K

##### Posttranslational Modifications

Carboxymethyl

##### Source File

20191211\_F1\_Ag5\_peng0013\_SA\_Flag\_Asp\_N.raw

##### Fraction

1

##### Scan Feature

F1:5891

##### De Novo Score

96

##### Confidence score

96

##### Mass Charge Ratio

533.2423

##### Mass

2128.9404

##### Charge

4

##### Retention Time

34.36

##### Predicted Retention Time

-

##### Area

1295400

##### Fragmentation Mode

ETHCD
