## Supplementary material for "Template-based assembly of proteomic short reads for *de novo* antibody sequencing and repertoire profiling": F1_6313.html

Details F1\_6313

OverviewUndefined

### Read F1:6313

#### Sequence

DKVSHDCMLT

#### Sequence Length

10

#### Meta Information from PEAKS

##### Scan Identifier

F1:6313

##### Original Sequence (length=26)

D

K

V

S

H

D

C

+58.01

M

+15.99

L

T

##### Posttranslational Modifications

Carboxymethyl; Oxidation (M)

##### Source File

20191211\_F1\_Ag5\_peng0013\_SA\_Flag\_Asp\_N.raw

##### Fraction

1

##### Scan Feature

F1:2042

##### De Novo Score

93

##### Confidence score

93

##### Mass Charge Ratio

408.1721

##### Mass

1221.5005

##### Charge

3

##### Retention Time

34.97

##### Predicted Retention Time

-

##### Area

2077400

##### Fragmentation Mode

ETHCD
