## Supplementary material for "Template-based assembly of proteomic short reads for *de novo* antibody sequencing and repertoire profiling": F1_6315.html

Details F1\_6315

OverviewUndefined

### Read F1:6315

#### Sequence

DKVSLTCMLT

#### Sequence Length

10

#### Meta Information from PEAKS

##### Scan Identifier

F1:6315

##### Original Sequence (length=26)

D

K

V

S

L

T

C

+58.01

M

+15.99

L

T

##### Posttranslational Modifications

Carboxymethyl; Oxidation (M)

##### Source File

20191211\_F1\_Ag5\_peng0013\_SA\_Flag\_Asp\_N.raw

##### Fraction

1

##### Scan Feature

F1:7803

##### De Novo Score

96

##### Confidence score

96

##### Mass Charge Ratio

592.7817

##### Mass

1183.5464

##### Charge

2

##### Retention Time

35.02

##### Predicted Retention Time

-

##### Area

2387500000

##### Parts Per Million

2.1

##### Fragmentation Mode

HCD
