## Supplementary material for "Template-based assembly of proteomic short reads for *de novo* antibody sequencing and repertoire profiling": F1_6370.html

Details F1\_6370

OverviewUndefined

### Read F1:6370

#### Sequence

DSTYSMSDTLTLTK

#### Sequence Length

14

#### Meta Information from PEAKS

##### Scan Identifier

F1:6370

##### Original Sequence (length=22)

D

S

T

Y

S

M

+15.99

S

D

T

L

T

L

T

K

##### Posttranslational Modifications

Oxidation (M)

##### Source File

20191211\_F1\_Ag5\_peng0013\_SA\_Flag\_Asp\_N.raw

##### Fraction

1

##### Scan Feature

F1:15073

##### De Novo Score

97

##### Confidence score

97

##### Mass Charge Ratio

789.8629

##### Mass

1577.7131

##### Charge

2

##### Retention Time

35.09

##### Predicted Retention Time

-

##### Area

41026000

##### Fragmentation Mode

ETHCD
