## Supplementary material for "Template-based assembly of proteomic short reads for *de novo* antibody sequencing and repertoire profiling": F1_6473.html

Details F1\_6473

OverviewUndefined

### Read F1:6473

#### Sequence

DPSSSTAYMEL

#### Sequence Length

11

#### Meta Information from PEAKS

##### Scan Identifier

F1:6473

##### Original Sequence (length=19)

D

P

S

S

S

T

A

Y

M

+15.99

E

L

##### Posttranslational Modifications

Oxidation (M)

##### Source File

20191211\_F1\_Ag5\_peng0013\_SA\_Flag\_Asp\_N.raw

##### Fraction

1

##### Scan Feature

F1:8358

##### De Novo Score

96

##### Confidence score

96

##### Mass Charge Ratio

608.7559

##### Mass

1215.4966

##### Charge

2

##### Retention Time

35.97

##### Predicted Retention Time

-

##### Area

6159500

##### Parts Per Million

0.5

##### Fragmentation Mode

HCD
