## Supplementary material for "Template-based assembly of proteomic short reads for *de novo* antibody sequencing and repertoire profiling": F1_6689.html

Details F1\_6689

OverviewUndefined

### Read F1:6689

#### Sequence

DLNVKMAKLDGSERQNGVLNSWT

#### Sequence Length

23

#### Meta Information from PEAKS

##### Scan Identifier

F1:6689

##### Original Sequence (length=31)

D

L

N

V

K

M

A

K

L

D

G

S

E

R

Q

N

G

V

L

N

S

W

+15.99

T

##### Posttranslational Modifications

Oxidation (HW)

##### Source File

20191211\_F1\_Ag5\_peng0013\_SA\_Flag\_Asp\_N.raw

##### Fraction

1

##### Scan Feature

F1:10321

##### De Novo Score

91

##### Confidence score

91

##### Mass Charge Ratio

648.5763

##### Mass

2590.2808

##### Charge

4

##### Retention Time

37.06

##### Predicted Retention Time

-

##### Area

1039400

##### Fragmentation Mode

ETHCD
