## Supplementary material for "Template-based assembly of proteomic short reads for *de novo* antibody sequencing and repertoire profiling": F1_10082.html

Details F1\_10082

OverviewUndefined

### Read F1:10082

#### Sequence

DTDGSYFKYKLNVQKSNWEQNTFTCSVLHEGLH

#### Sequence Length

33

#### Meta Information from PEAKS

##### Scan Identifier

F1:10082

##### Original Sequence (length=49)

D

T

D

G

S

Y

F

K

+58.01

Y

K

L

N

V

Q

K

S

N

W

E

Q

N

T

F

T

C

+58.01

S

V

L

H

E

G

L

H

##### Posttranslational Modifications

Carboxymethyl (KW X@N-term); Carboxymethyl

##### Source File

20191211\_F1\_Ag5\_peng0013\_SA\_Flag\_Asp\_N.raw

##### Fraction

1

##### Scan Feature

F1:21278

##### De Novo Score

91

##### Confidence score

91

##### Mass Charge Ratio

1001.9655

##### Mass

4003.8323

##### Charge

4

##### Retention Time

55.66

##### Predicted Retention Time

-

##### Area

14845000

##### Parts Per Million

0.1

##### Fragmentation Mode

ETHCD
