## Supplementary material for "Template-based assembly of proteomic short reads for *de novo* antibody sequencing and repertoire profiling": F1_10123.html

Details F1\_10123

OverviewUndefined

### Read F1:10123

#### Sequence

DETTLTADPSSSTAYMEKQLTSY

#### Sequence Length

23

#### Meta Information from PEAKS

##### Scan Identifier

F1:10123

##### Original Sequence (length=31)

D

E

T

T

L

T

A

D

P

S

S

S

T

A

Y

M

E

K

+58.01

Q

L

T

S

Y

##### Posttranslational Modifications

Carboxymethyl (KW X@N-term)

##### Source File

20191211\_F1\_Ag5\_peng0013\_SA\_Flag\_Asp\_N.raw

##### Fraction

1

##### Scan Feature

F1:17301

##### De Novo Score

94

##### Confidence score

94

##### Mass Charge Ratio

866.3964

##### Mass

2596.1375

##### Charge

3

##### Retention Time

56.09

##### Predicted Retention Time

-

##### Area

2986600

##### Parts Per Million

11.5

##### Fragmentation Mode

ETHCD
