## Supplementary material for "Template-based assembly of proteomic short reads for *de novo* antibody sequencing and repertoire profiling": F1_10203.html

Details F1\_10203

OverviewUndefined

### Read F1:10203

#### Sequence

DPSSSTASHSELNSLTSEDSAVYYCAR

#### Sequence Length

27

#### Meta Information from PEAKS

##### Scan Identifier

F1:10203

##### Original Sequence (length=35)

D

P

S

S

S

T

A

S

H

S

E

L

N

S

L

T

S

E

D

S

A

V

Y

Y

C

+58.01

A

R

##### Posttranslational Modifications

Carboxymethyl

##### Source File

20191211\_F1\_Ag5\_peng0013\_SA\_Flag\_Asp\_N.raw

##### Fraction

1

##### Scan Feature

F1:20964

##### De Novo Score

90

##### Confidence score

90

##### Mass Charge Ratio

979.0913

##### Mass

2934.2461

##### Charge

3

##### Retention Time

56.4

##### Predicted Retention Time

-

##### Area

1392800

##### Parts Per Million

2

##### Fragmentation Mode

ETHCD
