## Supplementary material for "Template-based assembly of proteomic short reads for *de novo* antibody sequencing and repertoire profiling": F1_10255.html

Details F1\_10255

OverviewUndefined

### Read F1:10255

#### Sequence

DALGVYYCFQGSHVPLPWVGLDAG

#### Sequence Length

24

#### Meta Information from PEAKS

##### Scan Identifier

F1:10255

##### Original Sequence (length=48)

D

+58.01

A

L

G

V

Y

Y

C

+58.01

F

Q

G

S

H

V

P

L

P

W

+15.99

V

G

L

D

A

G

##### Posttranslational Modifications

Carboxymethyl (KW X@N-term); Carboxymethyl; Oxidation (HW)

##### Source File

20191211\_F1\_Ag5\_peng0013\_SA\_Flag\_Asp\_N.raw

##### Fraction

1

##### Scan Feature

F1:18281

##### De Novo Score

91

##### Confidence score

91

##### Mass Charge Ratio

899.4167

##### Mass

2695.2263

##### Charge

3

##### Retention Time

56.77

##### Predicted Retention Time

-

##### Area

63076000

##### Parts Per Million

0.8

##### Fragmentation Mode

ETHCD
