## Supplementary material for "Template-based assembly of proteomic short reads for *de novo* antibody sequencing and repertoire profiling": F1_10259.html

Details F1\_10259

OverviewUndefined

### Read F1:10259

#### Sequence

DALGVYYCFQGHAVPYTFGGGTKL

#### Sequence Length

24

#### Meta Information from PEAKS

##### Scan Identifier

F1:10259

##### Original Sequence (length=48)

D

+58.01

A

L

G

V

Y

Y

C

+58.01

F

Q

G

H

+15.99

A

V

P

Y

T

F

G

G

G

T

K

L

##### Posttranslational Modifications

Carboxymethyl (KW X@N-term); Carboxymethyl; Oxidation (HW)

##### Source File

20191211\_F1\_Ag5\_peng0013\_SA\_Flag\_Asp\_N.raw

##### Fraction

1

##### Scan Feature

F1:11102

##### De Novo Score

90

##### Confidence score

90

##### Mass Charge Ratio

674.8152

##### Mass

2695.2263

##### Charge

4

##### Retention Time

56.77

##### Predicted Retention Time

-

##### Area

434290

##### Parts Per Million

2

##### Fragmentation Mode

HCD
