## Supplementary material for "Template-based assembly of proteomic short reads for *de novo* antibody sequencing and repertoire profiling": F1_10260.html

Details F1\_10260

OverviewUndefined

### Read F1:10260

#### Sequence

DAGAKPCLCTVPEVSSVFLFPPKPK

#### Sequence Length

25

#### Meta Information from PEAKS

##### Scan Identifier

F1:10260

##### Original Sequence (length=41)

D

A

G

A

K

P

C

+58.01

L

C

+58.01

T

V

P

E

V

S

S

V

F

L

F

P

P

K

P

K

##### Posttranslational Modifications

Carboxymethyl

##### Source File

20191211\_F1\_Ag5\_peng0013\_SA\_Flag\_Asp\_N.raw

##### Fraction

1

##### Scan Feature

F1:11559

##### De Novo Score

91

##### Confidence score

91

##### Mass Charge Ratio

687.3518

##### Mass

2745.3757

##### Charge

4

##### Retention Time

56.77

##### Predicted Retention Time

-

##### Area

7272400

##### Parts Per Million

0.9

##### Fragmentation Mode

HCD
