## Supplementary material for "Template-based assembly of proteomic short reads for *de novo* antibody sequencing and repertoire profiling": F1_10271.html

Details F1\_10271

OverviewUndefined

### Read F1:10271

#### Sequence

DQQKCCTVPEVSSVFLFPPKPK

#### Sequence Length

22

#### Meta Information from PEAKS

##### Scan Identifier

F1:10271

##### Original Sequence (length=38)

D

Q

Q

K

C

+58.01

C

+58.01

T

V

P

E

V

S

S

V

F

L

F

P

P

K

P

K

##### Posttranslational Modifications

Carboxymethyl

##### Source File

20191211\_F1\_Ag5\_peng0013\_SA\_Flag\_Asp\_N.raw

##### Fraction

1

##### Scan Feature

F1:10352

##### De Novo Score

92

##### Confidence score

92

##### Mass Charge Ratio

649.0747

##### Mass

2592.2603

##### Charge

4

##### Retention Time

56.82

##### Predicted Retention Time

-

##### Area

2701400

##### Parts Per Million

3.7

##### Fragmentation Mode

HCD
