## Supplementary material for "Template-based assembly of proteomic short reads for *de novo* antibody sequencing and repertoire profiling": F1_10308.html

Details F1\_10308

OverviewUndefined

### Read F1:10308

#### Sequence

DLTVEWQWNGQPAENYKNTQPLM

#### Sequence Length

23

#### Meta Information from PEAKS

##### Scan Identifier

F1:10308

##### Original Sequence (length=23)

D

L

T

V

E

W

Q

W

N

G

Q

P

A

E

N

Y

K

N

T

Q

P

L

M

##### Posttranslational Modifications

##### Source File

20191211\_F1\_Ag5\_peng0013\_SA\_Flag\_Asp\_N.raw

##### Fraction

1

##### Scan Feature

F1:19128

##### De Novo Score

98

##### Confidence score

98

##### Mass Charge Ratio

921.4347

##### Mass

2761.2805

##### Charge

3

##### Retention Time

57.04

##### Predicted Retention Time

-

##### Area

8566400

##### Parts Per Million

0.6

##### Fragmentation Mode

ETHCD
