## Supplementary material for "Template-based assembly of proteomic short reads for *de novo* antibody sequencing and repertoire profiling": F1_10331.html

Details F1\_10331

OverviewUndefined

### Read F1:10331

#### Sequence

DELGVYYAFQAGHVPYTFGGGKTL

#### Sequence Length

24

#### Meta Information from PEAKS

##### Scan Identifier

F1:10331

##### Original Sequence (length=32)

D

E

L

G

V

Y

Y

A

F

Q

A

G

H

+15.99

V

P

Y

T

F

G

G

G

K

T

L

##### Posttranslational Modifications

Oxidation (HW)

##### Source File

20191211\_F1\_Ag5\_peng0013\_SA\_Flag\_Asp\_N.raw

##### Fraction

1

##### Scan Feature

F1:17392

##### De Novo Score

91

##### Confidence score

91

##### Mass Charge Ratio

869.4232

##### Mass

2605.2488

##### Charge

3

##### Retention Time

57.2

##### Predicted Retention Time

-

##### Area

861000

##### Fragmentation Mode

ETHCD
