## Supplementary material for "Template-based assembly of proteomic short reads for *de novo* antibody sequencing and repertoire profiling": F1_10354.html

Details F1\_10354

OverviewUndefined

### Read F1:10354

#### Sequence

DLGVYYCYQGSHVPYTFGGGTKL

#### Sequence Length

23

#### Meta Information from PEAKS

##### Scan Identifier

F1:10354

##### Original Sequence (length=31)

D

L

G

V

Y

Y

C

+58.01

Y

Q

G

S

H

V

P

Y

T

F

G

G

G

T

K

L

##### Posttranslational Modifications

Carboxymethyl

##### Source File

20191211\_F1\_Ag5\_peng0013\_SA\_Flag\_Asp\_N.raw

##### Fraction

1

##### Scan Feature

F1:17151

##### De Novo Score

97

##### Confidence score

97

##### Mass Charge Ratio

861.7345

##### Mass

2582.1787

##### Charge

3

##### Retention Time

57.19

##### Predicted Retention Time

-

##### Area

27353000

##### Parts Per Million

1.1

##### Fragmentation Mode

ETHCD
