## Supplementary material for "Template-based assembly of proteomic short reads for *de novo* antibody sequencing and repertoire profiling": F1_10359.html

Details F1\_10359

OverviewUndefined

### Read F1:10359

#### Sequence

DATTLTADPSSSTAYMELNSLTSEDSAVYYCAR

#### Sequence Length

33

#### Meta Information from PEAKS

##### Scan Identifier

F1:10359

##### Original Sequence (length=57)

D

+58.01

A

T

T

L

T

A

D

P

S

S

S

T

A

Y

M

+15.99

E

L

N

S

L

T

S

E

D

S

A

V

Y

Y

C

+58.01

A

R

##### Posttranslational Modifications

Carboxymethyl (KW X@N-term); Oxidation (M); Carboxymethyl

##### Source File

20191211\_F1\_Ag5\_peng0013\_SA\_Flag\_Asp\_N.raw

##### Fraction

1

##### Scan Feature

F1:23949

##### De Novo Score

94

##### Confidence score

94

##### Mass Charge Ratio

1222.5262

##### Mass

3664.5557

##### Charge

3

##### Retention Time

57.47

##### Predicted Retention Time

-

##### Area

15752000

##### Parts Per Million

0.3

##### Fragmentation Mode

ETHCD
