## Supplementary material for "Template-based assembly of proteomic short reads for *de novo* antibody sequencing and repertoire profiling": F1_10385.html

Details F1\_10385

OverviewUndefined

### Read F1:10385

#### Sequence

DLTVEWQW

#### Sequence Length

8

#### Meta Information from PEAKS

##### Scan Identifier

F1:10385

##### Original Sequence (length=16)

D

L

T

V

E

W

Q

W

+15.99

##### Posttranslational Modifications

Oxidation (HW)

##### Source File

20191211\_F1\_Ag5\_peng0013\_SA\_Flag\_Asp\_N.raw

##### Fraction

1

##### Scan Feature

-

##### De Novo Score

97

##### Confidence score

97

##### Mass Charge Ratio

546.7534

##### Mass

1091.4924

##### Charge

2

##### Retention Time

57.55

##### Predicted Retention Time

-

##### Area

0

##### Fragmentation Mode

ETHCD

##### Also found in scans

F1:11822
