## Supplementary material for "Template-based assembly of proteomic short reads for *de novo* antibody sequencing and repertoire profiling": F1_10407.html

Details F1\_10407

OverviewUndefined

### Read F1:10407

#### Sequence

DCQKCPLCTVPEVSSVFLFPPKPK

#### Sequence Length

24

#### Meta Information from PEAKS

##### Scan Identifier

F1:10407

##### Original Sequence (length=48)

D

C

+58.01

Q

K

C

+58.01

P

L

C

+58.01

T

V

P

E

V

S

S

V

F

L

F

P

P

K

P

K

##### Posttranslational Modifications

Carboxymethyl

##### Source File

20191211\_F1\_Ag5\_peng0013\_SA\_Flag\_Asp\_N.raw

##### Fraction

1

##### Scan Feature

F1:12385

##### De Novo Score

90

##### Confidence score

90

##### Mass Charge Ratio

709.8442

##### Mass

2835.3533

##### Charge

4

##### Retention Time

57.57

##### Predicted Retention Time

-

##### Area

38467000

##### Fragmentation Mode

HCD
