## Supplementary material for "Template-based assembly of proteomic short reads for *de novo* antibody sequencing and repertoire profiling": F1_10426.html

Details F1\_10426

OverviewUndefined

### Read F1:10426

#### Sequence

DLPCMNMCTVPEVSSVFLFPPKPK

#### Sequence Length

24

#### Meta Information from PEAKS

##### Scan Identifier

F1:10426

##### Original Sequence (length=48)

D

L

P

C

+58.01

M

+15.99

N

M

C

+58.01

T

V

P

E

V

S

S

V

F

L

F

P

P

K

P

K

##### Posttranslational Modifications

Carboxymethyl; Oxidation (M)

##### Source File

20191211\_F1\_Ag5\_peng0013\_SA\_Flag\_Asp\_N.raw

##### Fraction

1

##### Scan Feature

F1:12115

##### De Novo Score

93

##### Confidence score

93

##### Mass Charge Ratio

703.5836

##### Mass

2810.304

##### Charge

4

##### Retention Time

57.82

##### Predicted Retention Time

-

##### Area

18611000

##### Parts Per Million

0.5

##### Fragmentation Mode

HCD
