## Supplementary material for "Template-based assembly of proteomic short reads for *de novo* antibody sequencing and repertoire profiling": F1_10462.html

Details F1\_10462

OverviewUndefined

### Read F1:10462

#### Sequence

DLGVYYCFQGSHVPYTFGGGTKL

#### Sequence Length

23

#### Meta Information from PEAKS

##### Scan Identifier

F1:10462

##### Original Sequence (length=31)

D

L

G

V

Y

Y

C

+58.01

F

Q

G

S

H

V

P

Y

T

F

G

G

G

T

K

L

##### Posttranslational Modifications

Carboxymethyl

##### Source File

20191211\_F1\_Ag5\_peng0013\_SA\_Flag\_Asp\_N.raw

##### Fraction

1

##### Scan Feature

F1:16972

##### De Novo Score

98

##### Confidence score

98

##### Mass Charge Ratio

856.4032

##### Mass

2566.1838

##### Charge

3

##### Retention Time

57.67

##### Predicted Retention Time

-

##### Area

588630000

##### Parts Per Million

1.5

##### Fragmentation Mode

ETHCD

##### Also found in scans

F1:10521 F1:10587
