## Supplementary material for "Template-based assembly of proteomic short reads for *de novo* antibody sequencing and repertoire profiling": F1_10484.html

Details F1\_10484

OverviewUndefined

### Read F1:10484

#### Sequence

DAGKAPCLCTVPEVSSVFLFPPKPK

#### Sequence Length

25

#### Meta Information from PEAKS

##### Scan Identifier

F1:10484

##### Original Sequence (length=41)

D

A

G

K

A

P

C

+58.01

L

C

+58.01

T

V

P

E

V

S

S

V

F

L

F

P

P

K

P

K

##### Posttranslational Modifications

Carboxymethyl

##### Source File

20191211\_F1\_Ag5\_peng0013\_SA\_Flag\_Asp\_N.raw

##### Fraction

1

##### Scan Feature

-

##### De Novo Score

90

##### Confidence score

90

##### Mass Charge Ratio

687.3503

##### Mass

2745.3757

##### Charge

4

##### Retention Time

58.15

##### Predicted Retention Time

-

##### Area

0

##### Fragmentation Mode

HCD
