## Supplementary material for "Template-based assembly of proteomic short reads for *de novo* antibody sequencing and repertoire profiling": F1_10517.html

Details F1\_10517

OverviewUndefined

### Read F1:10517

#### Sequence

DFAELTKVKDLTKVNKECCHG

#### Sequence Length

21

#### Meta Information from PEAKS

##### Scan Identifier

F1:10517

##### Original Sequence (length=45)

D

F

A

E

L

T

K

V

K

+58.01

D

L

T

K

V

N

K

E

C

+58.01

C

+58.01

H

G

##### Posttranslational Modifications

Carboxymethyl (KW X@N-term); Carboxymethyl

##### Source File

20191211\_F1\_Ag5\_peng0013\_SA\_Flag\_Asp\_N.raw

##### Fraction

1

##### Scan Feature

F1:9887

##### De Novo Score

93

##### Confidence score

93

##### Mass Charge Ratio

638.8051

##### Mass

2551.1934

##### Charge

4

##### Retention Time

58.34

##### Predicted Retention Time

-

##### Area

2737500

##### Fragmentation Mode

ETHCD
