## Supplementary material for "Template-based assembly of proteomic short reads for *de novo* antibody sequencing and repertoire profiling": F1_10529.html

Details F1\_10529

OverviewUndefined

### Read F1:10529

#### Sequence

DCGCKPCLCTVPEVSSVFLFPPKPK

#### Sequence Length

25

#### Meta Information from PEAKS

##### Scan Identifier

F1:10529

##### Original Sequence (length=57)

D

C

+58.01

G

C

+58.01

K

P

C

+58.01

L

C

+58.01

T

V

P

E

V

S

S

V

F

L

F

P

P

K

P

K

##### Posttranslational Modifications

Carboxymethyl

##### Source File

20191211\_F1\_Ag5\_peng0013\_SA\_Flag\_Asp\_N.raw

##### Fraction

1

##### Scan Feature

F1:13126

##### De Novo Score

97

##### Confidence score

97

##### Mass Charge Ratio

732.3407

##### Mass

2925.3308

##### Charge

4

##### Retention Time

58.4

##### Predicted Retention Time

-

##### Area

688030000

##### Parts Per Million

1

##### Fragmentation Mode

HCD

##### Also found in scans

F1:10513 F1:10500
