## Supplementary material for "Template-based assembly of proteomic short reads for *de novo* antibody sequencing and repertoire profiling": F1_10734.html

Details F1\_10734

OverviewUndefined

### Read F1:10734

#### Sequence

DLGVYYCFQGSNHKTTFGGGTKL

#### Sequence Length

23

#### Meta Information from PEAKS

##### Scan Identifier

F1:10734

##### Original Sequence (length=47)

D

L

G

V

Y

Y

C

+58.01

F

Q

G

S

N

H

+15.99

K

+58.01

T

T

F

G

G

G

T

K

L

##### Posttranslational Modifications

Carboxymethyl; Oxidation (HW); Carboxymethyl (KW X@N-term)

##### Source File

20191211\_F1\_Ag5\_peng0013\_SA\_Flag\_Asp\_N.raw

##### Fraction

1

##### Scan Feature

F1:17621

##### De Novo Score

91

##### Confidence score

91

##### Mass Charge Ratio

875.7373

##### Mass

2624.1853

##### Charge

3

##### Retention Time

59.72

##### Predicted Retention Time

-

##### Area

2733000

##### Parts Per Million

1.8

##### Fragmentation Mode

ETHCD
