## Supplementary material for "Template-based assembly of proteomic short reads for *de novo* antibody sequencing and repertoire profiling": F1_10840.html

Details F1\_10840

OverviewUndefined

### Read F1:10840

#### Sequence

DLSKDDPEVQFSSMFV

#### Sequence Length

16

#### Meta Information from PEAKS

##### Scan Identifier

F1:10840

##### Original Sequence (length=16)

D

L

S

K

D

D

P

E

V

Q

F

S

S

M

F

V

##### Posttranslational Modifications

##### Source File

20191211\_F1\_Ag5\_peng0013\_SA\_Flag\_Asp\_N.raw

##### Fraction

1

##### Scan Feature

F1:8647

##### De Novo Score

94

##### Confidence score

94

##### Mass Charge Ratio

615.2845

##### Mass

1842.8345

##### Charge

3

##### Retention Time

60.24

##### Predicted Retention Time

-

##### Area

930770

##### Fragmentation Mode

ETHCD
