## Supplementary material for "Template-based assembly of proteomic short reads for *de novo* antibody sequencing and repertoire profiling": F1_10904.html

Details F1\_10904

OverviewUndefined

### Read F1:10904

#### Sequence

DLPQFSWFVDDVEVHTAQTQPR

#### Sequence Length

22

#### Meta Information from PEAKS

##### Scan Identifier

F1:10904

##### Original Sequence (length=22)

D

L

P

Q

F

S

W

F

V

D

D

V

E

V

H

T

A

Q

T

Q

P

R

##### Posttranslational Modifications

##### Source File

20191211\_F1\_Ag5\_peng0013\_SA\_Flag\_Asp\_N.raw

##### Fraction

1

##### Scan Feature

F1:17500

##### De Novo Score

96

##### Confidence score

96

##### Mass Charge Ratio

872.4216

##### Mass

2614.2451

##### Charge

3

##### Retention Time

60.61

##### Predicted Retention Time

-

##### Area

368720

##### Fragmentation Mode

HCD
