## Supplementary material for "Template-based assembly of proteomic short reads for *de novo* antibody sequencing and repertoire profiling": F1_10956.html

Details F1\_10956

OverviewUndefined

### Read F1:10956

#### Sequence

DVLTLTLTPKDFDVVV

#### Sequence Length

16

#### Meta Information from PEAKS

##### Scan Identifier

F1:10956

##### Original Sequence (length=16)

D

V

L

T

L

T

L

T

P

K

D

F

D

V

V

V

##### Posttranslational Modifications

##### Source File

20191211\_F1\_Ag5\_peng0013\_SA\_Flag\_Asp\_N.raw

##### Fraction

1

##### Scan Feature

F1:7776

##### De Novo Score

92

##### Confidence score

92

##### Mass Charge Ratio

592.3342

##### Mass

1773.9763

##### Charge

3

##### Retention Time

61.23

##### Predicted Retention Time

-

##### Area

19289000

##### Parts Per Million

2.6

##### Fragmentation Mode

ETHCD
