## Supplementary material for "Template-based assembly of proteomic short reads for *de novo* antibody sequencing and repertoire profiling": F1_10973.html

Details F1\_10973

OverviewUndefined

### Read F1:10973

#### Sequence

DLAWLGYLNPSSGYAAYNQNFK

#### Sequence Length

22

#### Meta Information from PEAKS

##### Scan Identifier

F1:10973

##### Original Sequence (length=22)

D

L

A

W

L

G

Y

L

N

P

S

S

G

Y

A

A

Y

N

Q

N

F

K

##### Posttranslational Modifications

##### Source File

20191211\_F1\_Ag5\_peng0013\_SA\_Flag\_Asp\_N.raw

##### Fraction

1

##### Scan Feature

F1:16246

##### De Novo Score

94

##### Confidence score

94

##### Mass Charge Ratio

831.402

##### Mass

2491.1807

##### Charge

3

##### Retention Time

60.97

##### Predicted Retention Time

-

##### Area

812440

##### Parts Per Million

1.4

##### Fragmentation Mode

HCD
