## Supplementary material for "Template-based assembly of proteomic short reads for *de novo* antibody sequencing and repertoire profiling": F1_11057.html

Details F1\_11057

OverviewUndefined

### Read F1:11057

#### Sequence

DVLTLTLTPKVTCVVV

#### Sequence Length

16

#### Meta Information from PEAKS

##### Scan Identifier

F1:11057

##### Original Sequence (length=24)

D

V

L

T

L

T

L

T

P

K

V

T

C

+58.01

V

V

V

##### Posttranslational Modifications

Carboxymethyl

##### Source File

20191211\_F1\_Ag5\_peng0013\_SA\_Flag\_Asp\_N.raw

##### Fraction

1

##### Scan Feature

F1:7578

##### De Novo Score

98

##### Confidence score

98

##### Mass Charge Ratio

587.0016

##### Mass

1757.9849

##### Charge

3

##### Retention Time

61.55

##### Predicted Retention Time

-

##### Area

83099000

##### Fragmentation Mode

ETHCD

##### Also found in scans

F1:11535 F1:11332 F1:11449 F1:11942 F1:11601 F1:12007 F1:11417 F1:11770 F1:11362 F1:11538 F1:13817 F1:12199 F1:11766 F1:11576
