## Supplementary material for "Template-based assembly of proteomic short reads for *de novo* antibody sequencing and repertoire profiling": F1_11237.html

Details F1\_11237

OverviewUndefined

### Read F1:11237

#### Sequence

DVLTLTLTPKVTCVVVDLSK

#### Sequence Length

20

#### Meta Information from PEAKS

##### Scan Identifier

F1:11237

##### Original Sequence (length=28)

D

V

L

T

L

T

L

T

P

K

V

T

C

+58.01

V

V

V

D

L

S

K

##### Posttranslational Modifications

Carboxymethyl

##### Source File

20191211\_F1\_Ag5\_peng0013\_SA\_Flag\_Asp\_N.raw

##### Fraction

1

##### Scan Feature

F1:13195

##### De Novo Score

97

##### Confidence score

97

##### Mass Charge Ratio

734.748

##### Mass

2201.2229

##### Charge

3

##### Retention Time

61.94

##### Predicted Retention Time

-

##### Area

865930

##### Fragmentation Mode

HCD
