## Supplementary material for "Template-based assembly of proteomic short reads for *de novo* antibody sequencing and repertoire profiling": F1_11300.html

Details F1\_11300

OverviewUndefined

### Read F1:11300

#### Sequence

DVLTLTLTPKVTAPVV

#### Sequence Length

16

#### Meta Information from PEAKS

##### Scan Identifier

F1:11300

##### Original Sequence (length=16)

D

V

L

T

L

T

L

T

P

K

V

T

A

P

V

V

##### Posttranslational Modifications

##### Source File

20191211\_F1\_Ag5\_peng0013\_SA\_Flag\_Asp\_N.raw

##### Fraction

1

##### Scan Feature

F1:16383

##### De Novo Score

95

##### Confidence score

95

##### Mass Charge Ratio

834.0021

##### Mass

1665.9917

##### Charge

2

##### Retention Time

62.77

##### Predicted Retention Time

-

##### Area

4716000

##### Fragmentation Mode

ETHCD
