## Supplementary material for "Template-based assembly of proteomic short reads for *de novo* antibody sequencing and repertoire profiling": F1_11361.html

Details F1\_11361

OverviewUndefined

### Read F1:11361

#### Sequence

DALQQAKEDRALLLR

#### Sequence Length

15

#### Meta Information from PEAKS

##### Scan Identifier

F1:11361

##### Original Sequence (length=15)

D

A

L

Q

Q

A

K

E

D

R

A

L

L

L

R

##### Posttranslational Modifications

##### Source File

20191211\_F1\_Ag5\_peng0013\_SA\_Flag\_Asp\_N.raw

##### Fraction

1

##### Scan Feature

F1:7377

##### De Novo Score

92

##### Confidence score

92

##### Mass Charge Ratio

580.6635

##### Mass

1738.969

##### Charge

3

##### Retention Time

63.18

##### Predicted Retention Time

-

##### Area

198080

##### Fragmentation Mode

HCD
