## Supplementary material for "Template-based assembly of proteomic short reads for *de novo* antibody sequencing and repertoire profiling": F1_11449_002.html

Details F1\_11449\_002

OverviewUndefined

### Read F1:11449\_002

#### Sequence

DPLTLTKMTLTLDVVP

#### Sequence Length

16

#### Meta Information from PEAKS

##### Scan Identifier

F1:11449

##### Original Sequence (length=16)

D

P

L

T

L

T

K

M

T

L

T

L

D

V

V

P

##### Posttranslational Modifications

##### Source File

20191211\_F1\_Ag5\_peng0013\_SA\_Flag\_Asp\_N.raw

##### Fraction

1

##### Scan Feature

F1:17701

##### De Novo Score

92

##### Confidence score

92

##### Mass Charge Ratio

879.0089

##### Mass

1755.9692

##### Charge

2

##### Retention Time

63.65

##### Predicted Retention Time

-

##### Area

350650

##### Parts Per Million

19.3

##### Fragmentation Mode

ETHCD
