## Supplementary material for "Template-based assembly of proteomic short reads for *de novo* antibody sequencing and repertoire profiling": F1_11458.html

Details F1\_11458

OverviewUndefined

### Read F1:11458

#### Sequence

DPSSSTAYMELGGSLTSEDSAVYYCAREKFYGY

#### Sequence Length

33

#### Meta Information from PEAKS

##### Scan Identifier

F1:11458

##### Original Sequence (length=49)

D

P

S

S

S

T

A

Y

M

+15.99

E

L

G

G

S

L

T

S

E

D

S

A

V

Y

Y

C

+58.01

A

R

E

K

F

Y

G

Y

##### Posttranslational Modifications

Oxidation (M); Carboxymethyl

##### Source File

20191211\_F1\_Ag5\_peng0013\_SA\_Flag\_Asp\_N.raw

##### Fraction

1

##### Scan Feature

F1:24241

##### De Novo Score

99

##### Confidence score

99

##### Mass Charge Ratio

1241.2002

##### Mass

3720.5759

##### Charge

3

##### Retention Time

63.73

##### Predicted Retention Time

-

##### Area

212970000

##### Parts Per Million

0.8

##### Fragmentation Mode

HCD
