## Supplementary material for "Template-based assembly of proteomic short reads for *de novo* antibody sequencing and repertoire profiling": F1_11514.html

Details F1\_11514

OverviewUndefined

### Read F1:11514

#### Sequence

DLSKDDPEVQFSWFVDDVEVHTAQTQEPR

#### Sequence Length

29

#### Meta Information from PEAKS

##### Scan Identifier

F1:11514

##### Original Sequence (length=29)

D

L

S

K

D

D

P

E

V

Q

F

S

W

F

V

D

D

V

E

V

H

T

A

Q

T

Q

E

P

R

##### Posttranslational Modifications

##### Source File

20191211\_F1\_Ag5\_peng0013\_SA\_Flag\_Asp\_N.raw

##### Fraction

1

##### Scan Feature

F1:22939

##### De Novo Score

91

##### Confidence score

91

##### Mass Charge Ratio

1139.8665

##### Mass

3416.5796

##### Charge

3

##### Retention Time

64.11

##### Predicted Retention Time

-

##### Area

1216800

##### Fragmentation Mode

HCD
