## Supplementary material for "Template-based assembly of proteomic short reads for *de novo* antibody sequencing and repertoire profiling": F1_11533.html

Details F1\_11533

OverviewUndefined

### Read F1:11533

#### Sequence

DLSKDDPEVQFSWFVDDVEVHTAQTQPR

#### Sequence Length

28

#### Meta Information from PEAKS

##### Scan Identifier

F1:11533

##### Original Sequence (length=28)

D

L

S

K

D

D

P

E

V

Q

F

S

W

F

V

D

D

V

E

V

H

T

A

Q

T

Q

P

R

##### Posttranslational Modifications

##### Source File

20191211\_F1\_Ag5\_peng0013\_SA\_Flag\_Asp\_N.raw

##### Fraction

1

##### Scan Feature

F1:22321

##### De Novo Score

99

##### Confidence score

99

##### Mass Charge Ratio

1096.8538

##### Mass

3287.5369

##### Charge

3

##### Retention Time

64.17

##### Predicted Retention Time

-

##### Area

5165700

##### Parts Per Million

0.8

##### Fragmentation Mode

HCD

##### Also found in scans

F1:11510
