## Supplementary material for "Template-based assembly of proteomic short reads for *de novo* antibody sequencing and repertoire profiling": F1_11597.html

Details F1\_11597

OverviewUndefined

### Read F1:11597

#### Sequence

DALNFLRALY

#### Sequence Length

10

#### Meta Information from PEAKS

##### Scan Identifier

F1:11597

##### Original Sequence (length=18)

D

+58.01

A

L

N

F

L

R

A

L

Y

##### Posttranslational Modifications

Carboxymethyl (KW X@N-term)

##### Source File

20191211\_F1\_Ag5\_peng0013\_SA\_Flag\_Asp\_N.raw

##### Fraction

1

##### Scan Feature

F1:9287

##### De Novo Score

96

##### Confidence score

96

##### Mass Charge Ratio

627.3303

##### Mass

1252.645

##### Charge

2

##### Retention Time

64.59

##### Predicted Retention Time

-

##### Area

394120

##### Parts Per Million

0.9

##### Fragmentation Mode

HCD
