## Supplementary material for "Template-based assembly of proteomic short reads for *de novo* antibody sequencing and repertoire profiling": F1_11598.html

Details F1\_11598

OverviewUndefined

### Read F1:11598

#### Sequence

DDPEVQFSSMFV

#### Sequence Length

12

#### Meta Information from PEAKS

##### Scan Identifier

F1:11598

##### Original Sequence (length=12)

D

D

P

E

V

Q

F

S

S

M

F

V

##### Posttranslational Modifications

##### Source File

20191211\_F1\_Ag5\_peng0013\_SA\_Flag\_Asp\_N.raw

##### Fraction

1

##### Scan Feature

F1:11976

##### De Novo Score

96

##### Confidence score

96

##### Mass Charge Ratio

700.8041

##### Mass

1399.5964

##### Charge

2

##### Retention Time

64.59

##### Predicted Retention Time

-

##### Area

3803000

##### Fragmentation Mode

ETHCD
