## Supplementary material for "Template-based assembly of proteomic short reads for *de novo* antibody sequencing and repertoire profiling": F1_11612.html

Details F1\_11612

OverviewUndefined

### Read F1:11612

#### Sequence

DLSKDDPEVQFSWFVD

#### Sequence Length

16

#### Meta Information from PEAKS

##### Scan Identifier

F1:11612

##### Original Sequence (length=16)

D

L

S

K

D

D

P

E

V

Q

F

S

W

F

V

D

##### Posttranslational Modifications

##### Source File

20191211\_F1\_Ag5\_peng0013\_SA\_Flag\_Asp\_N.raw

##### Fraction

1

##### Scan Feature

F1:10096

##### De Novo Score

98

##### Confidence score

98

##### Mass Charge Ratio

642.9633

##### Mass

1925.8682

##### Charge

3

##### Retention Time

64.74

##### Predicted Retention Time

-

##### Area

463980

##### Fragmentation Mode

ETHCD

##### Also found in scans

F1:11617
