## Supplementary material for "Template-based assembly of proteomic short reads for *de novo* antibody sequencing and repertoire profiling": F1_11633.html

Details F1\_11633

OverviewUndefined

### Read F1:11633

#### Sequence

DDPEVQFSWFVDDVEVHTAQTQPR

#### Sequence Length

24

#### Meta Information from PEAKS

##### Scan Identifier

F1:11633

##### Original Sequence (length=24)

D

D

P

E

V

Q

F

S

W

F

V

D

D

V

E

V

H

T

A

Q

T

Q

P

R

##### Posttranslational Modifications

##### Source File

20191211\_F1\_Ag5\_peng0013\_SA\_Flag\_Asp\_N.raw

##### Fraction

1

##### Scan Feature

F1:20152

##### De Novo Score

98

##### Confidence score

98

##### Mass Charge Ratio

949.108

##### Mass

2844.2991

##### Charge

3

##### Retention Time

65.03

##### Predicted Retention Time

-

##### Area

8009100

##### Parts Per Million

1.1

##### Fragmentation Mode

HCD

##### Also found in scans

F9:11353 F1:11651 F9:11476
