## Supplementary material for "Template-based assembly of proteomic short reads for *de novo* antibody sequencing and repertoire profiling": F1_11642.html

Details F1\_11642

OverviewUndefined

### Read F1:11642

#### Sequence

DDPEVQFSWFVDDVEVHTAQTQHKT

#### Sequence Length

25

#### Meta Information from PEAKS

##### Scan Identifier

F1:11642

##### Original Sequence (length=33)

D

D

P

E

V

Q

F

S

W

F

V

D

D

V

E

V

H

T

A

Q

T

Q

H

+15.99

K

T

##### Posttranslational Modifications

Oxidation (HW)

##### Source File

20191211\_F1\_Ag5\_peng0013\_SA\_Flag\_Asp\_N.raw

##### Fraction

1

##### Scan Feature

F1:21171

##### De Novo Score

95

##### Confidence score

95

##### Mass Charge Ratio

992.1215

##### Mass

2973.3416

##### Charge

3

##### Retention Time

64.95

##### Predicted Retention Time

-

##### Area

2588600

##### Parts Per Million

0.3

##### Fragmentation Mode

HCD
