## Supplementary material for "Template-based assembly of proteomic short reads for *de novo* antibody sequencing and repertoire profiling": F1_11691.html

Details F1\_11691

OverviewUndefined

### Read F1:11691

#### Sequence

DLSKDDPEVQFSMAFV

#### Sequence Length

16

#### Meta Information from PEAKS

##### Scan Identifier

F1:11691

##### Original Sequence (length=24)

D

L

S

K

D

D

P

E

V

Q

F

S

M

+15.99

A

F

V

##### Posttranslational Modifications

Oxidation (M)

##### Source File

20191211\_F1\_Ag5\_peng0013\_SA\_Flag\_Asp\_N.raw

##### Fraction

1

##### Scan Feature

F1:19168

##### De Novo Score

91

##### Confidence score

91

##### Mass Charge Ratio

922.4226

##### Mass

1842.8345

##### Charge

2

##### Retention Time

64.95

##### Predicted Retention Time

-

##### Area

7710900

##### Fragmentation Mode

ETHCD
