## Supplementary material for "Template-based assembly of proteomic short reads for *de novo* antibody sequencing and repertoire profiling": F1_11705.html

Details F1\_11705

OverviewUndefined

### Read F1:11705

#### Sequence

DETTLTADPSSSTAYMELNSLTSEDSAVYYYTTDATLTDHG

#### Sequence Length

41

#### Meta Information from PEAKS

##### Scan Identifier

F1:11705

##### Original Sequence (length=49)

D

E

T

T

L

T

A

D

P

S

S

S

T

A

Y

M

+15.99

E

L

N

S

L

T

S

E

D

S

A

V

Y

Y

Y

T

T

D

A

T

L

T

D

H

G

##### Posttranslational Modifications

Oxidation (M)

##### Source File

20191211\_F1\_Ag5\_peng0013\_SA\_Flag\_Asp\_N.raw

##### Fraction

1

##### Scan Feature

F1:25283

##### De Novo Score

98

##### Confidence score

98

##### Mass Charge Ratio

1484.9763

##### Mass

4451.9126

##### Charge

3

##### Retention Time

65.31

##### Predicted Retention Time

-

##### Area

42575000

##### Fragmentation Mode

HCD
