## Supplementary material for "Template-based assembly of proteomic short reads for *de novo* antibody sequencing and repertoire profiling": F1_11719.html

Details F1\_11719

OverviewUndefined

### Read F1:11719

#### Sequence

DLKAFLNNFYPK

#### Sequence Length

12

#### Meta Information from PEAKS

##### Scan Identifier

F1:11719

##### Original Sequence (length=12)

D

L

K

A

F

L

N

N

F

Y

P

K

##### Posttranslational Modifications

##### Source File

20191211\_F1\_Ag5\_peng0013\_SA\_Flag\_Asp\_N.raw

##### Fraction

1

##### Scan Feature

F1:13224

##### De Novo Score

93

##### Confidence score

93

##### Mass Charge Ratio

735.3948

##### Mass

1468.7715

##### Charge

2

##### Retention Time

65.26

##### Predicted Retention Time

-

##### Area

158790

##### Parts Per Million

2.4

##### Fragmentation Mode

HCD
