## Supplementary material for "Template-based assembly of proteomic short reads for *de novo* antibody sequencing and repertoire profiling": F1_11750.html

Details F1\_11750

OverviewUndefined

### Read F1:11750

#### Sequence

DPHTNATYVQFSWFVDDVEVH

#### Sequence Length

21

#### Meta Information from PEAKS

##### Scan Identifier

F1:11750

##### Original Sequence (length=21)

D

P

H

T

N

A

T

Y

V

Q

F

S

W

F

V

D

D

V

E

V

H

##### Posttranslational Modifications

##### Source File

20191211\_F1\_Ag5\_peng0013\_SA\_Flag\_Asp\_N.raw

##### Fraction

1

##### Scan Feature

F1:16438

##### De Novo Score

92

##### Confidence score

92

##### Mass Charge Ratio

836.0504

##### Mass

2505.1235

##### Charge

3

##### Retention Time

65.48

##### Predicted Retention Time

-

##### Area

195350

##### Parts Per Million

2.3

##### Fragmentation Mode

ETHCD
