## Supplementary material for "Template-based assembly of proteomic short reads for *de novo* antibody sequencing and repertoire profiling": F1_11757.html

Details F1\_11757

OverviewUndefined

### Read F1:11757

#### Sequence

DATTLTADPSSSTAYMELNSLTSWASVYYYWDATLTDGH

#### Sequence Length

39

#### Meta Information from PEAKS

##### Scan Identifier

F1:11757

##### Original Sequence (length=71)

D

+58.01

A

T

T

L

T

A

D

P

S

S

S

T

A

Y

M

+15.99

E

L

N

S

L

T

S

W

+58.01

A

S

V

Y

Y

Y

W

+15.99

D

A

T

L

T

D

G

H

##### Posttranslational Modifications

Carboxymethyl (KW X@N-term); Oxidation (M); Oxidation (HW)

##### Source File

20191211\_F1\_Ag5\_peng0013\_SA\_Flag\_Asp\_N.raw

##### Fraction

1

##### Scan Feature

F1:22501

##### De Novo Score

94

##### Confidence score

94

##### Mass Charge Ratio

1113.9844

##### Mass

4451.9067

##### Charge

4

##### Retention Time

65.31

##### Predicted Retention Time

-

##### Area

11989000

##### Parts Per Million

0.4

##### Fragmentation Mode

HCD
