## Supplementary material for "Template-based assembly of proteomic short reads for *de novo* antibody sequencing and repertoire profiling": F1_11803.html

Details F1\_11803

OverviewUndefined

### Read F1:11803

#### Sequence

DKNYDDLTVEWQWNGQPAENYKNTPQLM

#### Sequence Length

28

#### Meta Information from PEAKS

##### Scan Identifier

F1:11803

##### Original Sequence (length=36)

D

K

N

Y

D

D

L

T

V

E

W

Q

W

N

G

Q

P

A

E

N

Y

K

N

T

P

Q

L

M

+15.99

##### Posttranslational Modifications

Oxidation (M)

##### Source File

20191211\_F1\_Ag5\_peng0013\_SA\_Flag\_Asp\_N.raw

##### Fraction

1

##### Scan Feature

F1:22918

##### De Novo Score

92

##### Confidence score

92

##### Mass Charge Ratio

1138.5183

##### Mass

3412.5305

##### Charge

3

##### Retention Time

65.82

##### Predicted Retention Time

-

##### Area

28683000

##### Parts Per Million

0.8

##### Fragmentation Mode

ETHCD
