## Supplementary material for "Template-based assembly of proteomic short reads for *de novo* antibody sequencing and repertoire profiling": F1_11824.html

Details F1\_11824

OverviewUndefined

### Read F1:11824

#### Sequence

DVLTPNLPLSLPVSLG

#### Sequence Length

16

#### Meta Information from PEAKS

##### Scan Identifier

F1:11824

##### Original Sequence (length=16)

D

V

L

T

P

N

L

P

L

S

L

P

V

S

L

G

##### Posttranslational Modifications

##### Source File

20191211\_F1\_Ag5\_peng0013\_SA\_Flag\_Asp\_N.raw

##### Fraction

1

##### Scan Feature

F1:6226

##### De Novo Score

91

##### Confidence score

91

##### Mass Charge Ratio

545.6503

##### Mass

1633.929

##### Charge

3

##### Retention Time

65.94

##### Predicted Retention Time

-

##### Area

11095000

##### Parts Per Million

0

##### Fragmentation Mode

ETHCD
