## Supplementary material for "Template-based assembly of proteomic short reads for *de novo* antibody sequencing and repertoire profiling": F1_11828.html

Details F1\_11828

OverviewUndefined

### Read F1:11828

#### Sequence

DVLTLTLTVPKTCVVV

#### Sequence Length

16

#### Meta Information from PEAKS

##### Scan Identifier

F1:11828

##### Original Sequence (length=24)

D

V

L

T

L

T

L

T

V

P

K

T

C

+58.01

V

V

V

##### Posttranslational Modifications

Carboxymethyl

##### Source File

20191211\_F1\_Ag5\_peng0013\_SA\_Flag\_Asp\_N.raw

##### Fraction

1

##### Scan Feature

-

##### De Novo Score

92

##### Confidence score

92

##### Mass Charge Ratio

880.0005

##### Mass

1757.9849

##### Charge

2

##### Retention Time

65.93

##### Predicted Retention Time

-

##### Area

0

##### Parts Per Million

1

##### Fragmentation Mode

ETHCD
