## Supplementary material for "Template-based assembly of proteomic short reads for *de novo* antibody sequencing and repertoire profiling": F1_11868.html

Details F1\_11868

OverviewUndefined

### Read F1:11868

#### Sequence

DKAVCFLNNFYPK

#### Sequence Length

13

#### Meta Information from PEAKS

##### Scan Identifier

F1:11868

##### Original Sequence (length=21)

D

K

A

V

C

+58.01

F

L

N

N

F

Y

P

K

##### Posttranslational Modifications

Carboxymethyl

##### Source File

20191211\_F1\_Ag5\_peng0013\_SA\_Flag\_Asp\_N.raw

##### Fraction

1

##### Scan Feature

F1:15693

##### De Novo Score

95

##### Confidence score

95

##### Mass Charge Ratio

808.8929

##### Mass

1615.7705

##### Charge

2

##### Retention Time

66.11

##### Predicted Retention Time

-

##### Area

1733700

##### Parts Per Million

0.4

##### Fragmentation Mode

HCD
