## Supplementary material for "Template-based assembly of proteomic short reads for *de novo* antibody sequencing and repertoire profiling": F1_11913.html

Details F1\_11913

OverviewUndefined

### Read F1:11913

#### Sequence

DVLTLTKGTLTLTWVV

#### Sequence Length

16

#### Meta Information from PEAKS

##### Scan Identifier

F1:11913

##### Original Sequence (length=32)

D

+58.01

V

L

T

L

T

K

G

T

L

T

L

T

W

+15.99

V

V

##### Posttranslational Modifications

Carboxymethyl (KW X@N-term); Oxidation (HW)

##### Source File

20191211\_F1\_Ag5\_peng0013\_SA\_Flag\_Asp\_N.raw

##### Fraction

1

##### Scan Feature

F1:18978

##### De Novo Score

92

##### Confidence score

92

##### Mass Charge Ratio

917.5142

##### Mass

1833.0134

##### Charge

2

##### Retention Time

66.43

##### Predicted Retention Time

-

##### Area

1119400

##### Parts Per Million

0.2

##### Fragmentation Mode

ETHCD
