## Supplementary material for "Template-based assembly of proteomic short reads for *de novo* antibody sequencing and repertoire profiling": F1_11933.html

Details F1\_11933

OverviewUndefined

### Read F1:11933

#### Sequence

DKLMTQLPLSLPVSLG

#### Sequence Length

16

#### Meta Information from PEAKS

##### Scan Identifier

F1:11933

##### Original Sequence (length=32)

D

K

+58.01

L

M

+15.99

T

Q

L

P

L

S

L

P

V

S

L

G

##### Posttranslational Modifications

Carboxymethyl (KW X@N-term); Oxidation (M)

##### Source File

20191211\_F1\_Ag5\_peng0013\_SA\_Flag\_Asp\_N.raw

##### Fraction

1

##### Scan Feature

F1:18142

##### De Novo Score

97

##### Confidence score

97

##### Mass Charge Ratio

893.4874

##### Mass

1784.9595

##### Charge

2

##### Retention Time

66.69

##### Predicted Retention Time

-

##### Area

24392000

##### Parts Per Million

0.5

##### Fragmentation Mode

ETHCD
