## Supplementary material for "Template-based assembly of proteomic short reads for *de novo* antibody sequencing and repertoire profiling": F1_11944.html

Details F1\_11944

OverviewUndefined

### Read F1:11944

#### Sequence

DLSKDDPEVQFYNFV

#### Sequence Length

15

#### Meta Information from PEAKS

##### Scan Identifier

F1:11944

##### Original Sequence (length=15)

D

L

S

K

D

D

P

E

V

Q

F

Y

N

F

V

##### Posttranslational Modifications

##### Source File

20191211\_F1\_Ag5\_peng0013\_SA\_Flag\_Asp\_N.raw

##### Fraction

1

##### Scan Feature

F1:8250

##### De Novo Score

98

##### Confidence score

98

##### Mass Charge Ratio

605.9529

##### Mass

1814.8362

##### Charge

3

##### Retention Time

66.72

##### Predicted Retention Time

-

##### Area

9812200

##### Parts Per Million

0.5

##### Fragmentation Mode

ETHCD
