## Supplementary material for "Template-based assembly of proteomic short reads for *de novo* antibody sequencing and repertoire profiling": F1_11959.html

Details F1\_11959

OverviewUndefined

### Read F1:11959

#### Sequence

DVVVCFLNNFYPK

#### Sequence Length

13

#### Meta Information from PEAKS

##### Scan Identifier

F1:11959

##### Original Sequence (length=29)

D

+58.01

V

V

V

C

+58.01

F

L

N

N

F

Y

P

K

##### Posttranslational Modifications

Carboxymethyl (KW X@N-term); Carboxymethyl

##### Source File

20191211\_F1\_Ag5\_peng0013\_SA\_Flag\_Asp\_N.raw

##### Fraction

1

##### Scan Feature

F1:16464

##### De Novo Score

92

##### Confidence score

92

##### Mass Charge Ratio

837.4034

##### Mass

1672.7808

##### Charge

2

##### Retention Time

66.58

##### Predicted Retention Time

-

##### Area

788410

##### Parts Per Million

6.9

##### Fragmentation Mode

HCD
