## Supplementary material for "Template-based assembly of proteomic short reads for *de novo* antibody sequencing and repertoire profiling": F1_12025.html

Details F1\_12025

OverviewUndefined

### Read F1:12025

#### Sequence

DVLMTQPLLSPLVKE

#### Sequence Length

15

#### Meta Information from PEAKS

##### Scan Identifier

F1:12025

##### Original Sequence (length=23)

D

V

L

M

+15.99

T

Q

P

L

L

S

P

L

V

K

E

##### Posttranslational Modifications

Oxidation (M)

##### Source File

20191211\_F1\_Ag5\_peng0013\_SA\_Flag\_Asp\_N.raw

##### Fraction

1

##### Scan Feature

F1:6939

##### De Novo Score

92

##### Confidence score

92

##### Mass Charge Ratio

566.9827

##### Mass

1697.9272

##### Charge

3

##### Retention Time

67.18

##### Predicted Retention Time

-

##### Area

32385000

##### Fragmentation Mode

ETHCD
