## Supplementary material for "Template-based assembly of proteomic short reads for *de novo* antibody sequencing and repertoire profiling": F1_12191.html

Details F1\_12191

OverviewUndefined

### Read F1:12191

#### Sequence

DVLMTQLPLSLPVSLG

#### Sequence Length

16

#### Meta Information from PEAKS

##### Scan Identifier

F1:12191

##### Original Sequence (length=24)

D

V

L

M

+15.99

T

Q

L

P

L

S

L

P

V

S

L

G

##### Posttranslational Modifications

Oxidation (M)

##### Source File

20191211\_F1\_Ag5\_peng0013\_SA\_Flag\_Asp\_N.raw

##### Fraction

1

##### Scan Feature

F1:16752

##### De Novo Score

97

##### Confidence score

97

##### Mass Charge Ratio

849.9706

##### Mass

1697.9272

##### Charge

2

##### Retention Time

67.36

##### Predicted Retention Time

-

##### Area

3578400000

##### Fragmentation Mode

ETHCD

##### Also found in scans

F1:12993 F1:12546
