## Supplementary material for "Template-based assembly of proteomic short reads for *de novo* antibody sequencing and repertoire profiling": F1_12230.html

Details F1\_12230

OverviewUndefined

### Read F1:12230

#### Sequence

DLSKDDPEVQFSWFV

#### Sequence Length

15

#### Meta Information from PEAKS

##### Scan Identifier

F1:12230

##### Original Sequence (length=15)

D

L

S

K

D

D

P

E

V

Q

F

S

W

F

V

##### Posttranslational Modifications

##### Source File

20191211\_F1\_Ag5\_peng0013\_SA\_Flag\_Asp\_N.raw

##### Fraction

1

##### Scan Feature

F1:8207

##### De Novo Score

99

##### Confidence score

99

##### Mass Charge Ratio

604.6203

##### Mass

1810.8413

##### Charge

3

##### Retention Time

68.29

##### Predicted Retention Time

-

##### Area

24833000

##### Fragmentation Mode

ETHCD

##### Also found in scans

F1:11949 F1:12340 F1:11454 F1:12450 F1:12485
