## Supplementary material for "Template-based assembly of proteomic short reads for *de novo* antibody sequencing and repertoire profiling": F1_12247.html

Details F1\_12247

OverviewUndefined

### Read F1:12247

#### Sequence

DLSSEAAPEVQFSWFV

#### Sequence Length

16

#### Meta Information from PEAKS

##### Scan Identifier

F1:12247

##### Original Sequence (length=16)

D

L

S

S

E

A

A

P

E

V

Q

F

S

W

F

V

##### Posttranslational Modifications

##### Source File

20191211\_F1\_Ag5\_peng0013\_SA\_Flag\_Asp\_N.raw

##### Fraction

1

##### Scan Feature

F1:18556

##### De Novo Score

94

##### Confidence score

94

##### Mass Charge Ratio

906.4275

##### Mass

1810.8413

##### Charge

2

##### Retention Time

68.29

##### Predicted Retention Time

-

##### Area

1099500000

##### Fragmentation Mode

ETHCD
