## Supplementary material for "Template-based assembly of proteomic short reads for *de novo* antibody sequencing and repertoire profiling": F1_12262.html

Details F1\_12262

OverviewUndefined

### Read F1:12262

#### Sequence

DLSKDDPEVHSGSWFV

#### Sequence Length

16

#### Meta Information from PEAKS

##### Scan Identifier

F1:12262

##### Original Sequence (length=24)

D

L

S

K

D

D

P

E

V

H

+15.99

S

G

S

W

F

V

##### Posttranslational Modifications

Oxidation (HW)

##### Source File

20191211\_F1\_Ag5\_peng0013\_SA\_Flag\_Asp\_N.raw

##### Fraction

1

##### Scan Feature

-

##### De Novo Score

95

##### Confidence score

95

##### Mass Charge Ratio

917.4141

##### Mass

1832.8218

##### Charge

2

##### Retention Time

68.39

##### Predicted Retention Time

-

##### Area

0

##### Fragmentation Mode

HCD
