## Supplementary material for "Template-based assembly of proteomic short reads for *de novo* antibody sequencing and repertoire profiling": F1_12346.html

Details F1\_12346

OverviewUndefined

### Read F1:12346

#### Sequence

DDPEVQFSWFVD

#### Sequence Length

12

#### Meta Information from PEAKS

##### Scan Identifier

F1:12346

##### Original Sequence (length=12)

D

D

P

E

V

Q

F

S

W

F

V

D

##### Posttranslational Modifications

##### Source File

20191211\_F1\_Ag5\_peng0013\_SA\_Flag\_Asp\_N.raw

##### Fraction

1

##### Scan Feature

F1:13452

##### De Novo Score

96

##### Confidence score

96

##### Mass Charge Ratio

742.3223

##### Mass

1482.6304

##### Charge

2

##### Retention Time

68.84

##### Predicted Retention Time

-

##### Area

3777200

##### Fragmentation Mode

HCD
