## Supplementary material for "Template-based assembly of proteomic short reads for *de novo* antibody sequencing and repertoire profiling": F1_12451.html

Details F1\_12451

OverviewUndefined

### Read F1:12451

#### Sequence

DAALTSGGASVVCFLNNFYPK

#### Sequence Length

21

#### Meta Information from PEAKS

##### Scan Identifier

F1:12451

##### Original Sequence (length=29)

D

A

A

L

T

S

G

G

A

S

V

V

C

+58.01

F

L

N

N

F

Y

P

K

##### Posttranslational Modifications

Carboxymethyl

##### Source File

20191211\_F1\_Ag5\_peng0013\_SA\_Flag\_Asp\_N.raw

##### Fraction

1

##### Scan Feature

F1:22531

##### De Novo Score

99

##### Confidence score

99

##### Mass Charge Ratio

1116.5349

##### Mass

2231.0569

##### Charge

2

##### Retention Time

69.54

##### Predicted Retention Time

-

##### Area

1860000

##### Fragmentation Mode

HCD

##### Also found in scans

F1:12510
