## Supplementary material for "Template-based assembly of proteomic short reads for *de novo* antibody sequencing and repertoire profiling": F1_12463.html

Details F1\_12463

OverviewUndefined

### Read F1:12463

#### Sequence

DTQLTSNASVVCFLNNFYPK

#### Sequence Length

20

#### Meta Information from PEAKS

##### Scan Identifier

F1:12463

##### Original Sequence (length=28)

D

T

Q

L

T

S

N

A

S

V

V

C

+58.01

F

L

N

N

F

Y

P

K

##### Posttranslational Modifications

Carboxymethyl

##### Source File

20191211\_F1\_Ag5\_peng0013\_SA\_Flag\_Asp\_N.raw

##### Fraction

1

##### Scan Feature

F1:23177

##### De Novo Score

94

##### Confidence score

94

##### Mass Charge Ratio

1160.0519

##### Mass

2318.0889

##### Charge

2

##### Retention Time

69.54

##### Predicted Retention Time

-

##### Area

170270

##### Parts Per Million

0.1

##### Fragmentation Mode

HCD
