## Supplementary material for "Template-based assembly of proteomic short reads for *de novo* antibody sequencing and repertoire profiling": F1_12523.html

Details F1\_12523

OverviewUndefined

### Read F1:12523

#### Sequence

DVLMTQLPLDLPVSLG

#### Sequence Length

16

#### Meta Information from PEAKS

##### Scan Identifier

F1:12523

##### Original Sequence (length=24)

D

V

L

M

+15.99

T

Q

L

P

L

D

L

P

V

S

L

G

##### Posttranslational Modifications

Oxidation (M)

##### Source File

20191211\_F1\_Ag5\_peng0013\_SA\_Flag\_Asp\_N.raw

##### Fraction

1

##### Scan Feature

F1:17214

##### De Novo Score

91

##### Confidence score

91

##### Mass Charge Ratio

863.9678

##### Mass

1725.9224

##### Charge

2

##### Retention Time

69.88

##### Predicted Retention Time

-

##### Area

46053000

##### Fragmentation Mode

HCD
