## Supplementary material for "Template-based assembly of proteomic short reads for *de novo* antibody sequencing and repertoire profiling": F1_12626.html

Details F1\_12626

OverviewUndefined

### Read F1:12626

#### Sequence

DLSDKDPEVQFDWFV

#### Sequence Length

15

#### Meta Information from PEAKS

##### Scan Identifier

F1:12626

##### Original Sequence (length=15)

D

L

S

D

K

D

P

E

V

Q

F

D

W

F

V

##### Posttranslational Modifications

##### Source File

20191211\_F1\_Ag5\_peng0013\_SA\_Flag\_Asp\_N.raw

##### Fraction

1

##### Scan Feature

F1:19091

##### De Novo Score

94

##### Confidence score

94

##### Mass Charge Ratio

920.4254

##### Mass

1838.8362

##### Charge

2

##### Retention Time

70.62

##### Predicted Retention Time

-

##### Area

3095400

##### Parts Per Million

0.1

##### Fragmentation Mode

ETHCD
