## Supplementary material for "Template-based assembly of proteomic short reads for *de novo* antibody sequencing and repertoire profiling": F1_12932.html

Details F1\_12932

OverviewUndefined

### Read F1:12932

#### Sequence

DVPFTQLPLSLPVSVA

#### Sequence Length

16

#### Meta Information from PEAKS

##### Scan Identifier

F1:12932

##### Original Sequence (length=16)

D

V

P

F

T

Q

L

P

L

S

L

P

V

S

V

A

##### Posttranslational Modifications

##### Source File

20191211\_F1\_Ag5\_peng0013\_SA\_Flag\_Asp\_N.raw

##### Fraction

1

##### Scan Feature

-

##### De Novo Score

93

##### Confidence score

93

##### Mass Charge Ratio

841.9733

##### Mass

1681.929

##### Charge

2

##### Retention Time

72.21

##### Predicted Retention Time

-

##### Area

0

##### Parts Per Million

1.9

##### Fragmentation Mode

ETHCD
