## Supplementary material for "Template-based assembly of proteomic short reads for *de novo* antibody sequencing and repertoire profiling": F1_13023.html

Details F1\_13023

OverviewUndefined

### Read F1:13023

#### Sequence

DDPEVHSGSWFV

#### Sequence Length

12

#### Meta Information from PEAKS

##### Scan Identifier

F1:13023

##### Original Sequence (length=20)

D

D

P

E

V

H

+15.99

S

G

S

W

F

V

##### Posttranslational Modifications

Oxidation (HW)

##### Source File

20191211\_F1\_Ag5\_peng0013\_SA\_Flag\_Asp\_N.raw

##### Fraction

1

##### Scan Feature

F1:11785

##### De Novo Score

91

##### Confidence score

91

##### Mass Charge Ratio

695.7993

##### Mass

1389.5837

##### Charge

2

##### Retention Time

72.85

##### Predicted Retention Time

-

##### Area

1576700

##### Parts Per Million

0.2

##### Fragmentation Mode

HCD
