## Supplementary material for "Template-based assembly of proteomic short reads for *de novo* antibody sequencing and repertoire profiling": F1_13063.html

Details F1\_13063

OverviewUndefined

### Read F1:13063

#### Sequence

DPEVQFSWFV

#### Sequence Length

10

#### Meta Information from PEAKS

##### Scan Identifier

F1:13063

##### Original Sequence (length=10)

D

P

E

V

Q

F

S

W

F

V

##### Posttranslational Modifications

##### Source File

20191211\_F1\_Ag5\_peng0013\_SA\_Flag\_Asp\_N.raw

##### Fraction

1

##### Scan Feature

F1:9266

##### De Novo Score

98

##### Confidence score

98

##### Mass Charge Ratio

627.2952

##### Mass

1252.5764

##### Charge

2

##### Retention Time

73.03

##### Predicted Retention Time

-

##### Area

2053500

##### Fragmentation Mode

ETHCD
