## Supplementary material for "Template-based assembly of proteomic short reads for *de novo* antibody sequencing and repertoire profiling": F1_13158.html

Details F1\_13158

OverviewUndefined

### Read F1:13158

#### Sequence

DAAPTVSLFPPSSEQLTSGGASVVCFL

#### Sequence Length

27

#### Meta Information from PEAKS

##### Scan Identifier

F1:13158

##### Original Sequence (length=35)

D

A

A

P

T

V

S

L

F

P

P

S

S

E

Q

L

T

S

G

G

A

S

V

V

C

+58.01

F

L

##### Posttranslational Modifications

Carboxymethyl

##### Source File

20191211\_F1\_Ag5\_peng0013\_SA\_Flag\_Asp\_N.raw

##### Fraction

1

##### Scan Feature

F1:18828

##### De Novo Score

98

##### Confidence score

98

##### Mass Charge Ratio

913.4458

##### Mass

2737.3154

##### Charge

3

##### Retention Time

73.32

##### Predicted Retention Time

-

##### Area

32001000

##### Parts Per Million

0.1

##### Fragmentation Mode

ETHCD
