## Supplementary material for "Template-based assembly of proteomic short reads for *de novo* antibody sequencing and repertoire profiling": F1_13183.html

Details F1\_13183

OverviewUndefined

### Read F1:13183

#### Sequence

DVLMTQLPLSLPVSVA

#### Sequence Length

16

#### Meta Information from PEAKS

##### Scan Identifier

F1:13183

##### Original Sequence (length=16)

D

V

L

M

T

Q

L

P

L

S

L

P

V

S

V

A

##### Posttranslational Modifications

##### Source File

20191211\_F1\_Ag5\_peng0013\_SA\_Flag\_Asp\_N.raw

##### Fraction

1

##### Scan Feature

-

##### De Novo Score

91

##### Confidence score

91

##### Mass Charge Ratio

841.9731

##### Mass

1681.9324

##### Charge

2

##### Retention Time

73.58

##### Predicted Retention Time

-

##### Area

0

##### Fragmentation Mode

ETHCD
