## Supplementary material for "Template-based assembly of proteomic short reads for *de novo* antibody sequencing and repertoire profiling": F1_13583.html

Details F1\_13583

OverviewUndefined

### Read F1:13583

#### Sequence

DVLMTQLPLDLPVSAV

#### Sequence Length

16

#### Meta Information from PEAKS

##### Scan Identifier

F1:13583

##### Original Sequence (length=16)

D

V

L

M

T

Q

L

P

L

D

L

P

V

S

A

V

##### Posttranslational Modifications

##### Source File

20191211\_F1\_Ag5\_peng0013\_SA\_Flag\_Asp\_N.raw

##### Fraction

1

##### Scan Feature

F1:16955

##### De Novo Score

97

##### Confidence score

97

##### Mass Charge Ratio

855.9706

##### Mass

1709.9272

##### Charge

2

##### Retention Time

75.75

##### Predicted Retention Time

-

##### Area

4618500

##### Fragmentation Mode

ETHCD
