## Supplementary material for "Template-based assembly of proteomic short reads for *de novo* antibody sequencing and repertoire profiling": F1_13652.html

Details F1\_13652

OverviewUndefined

### Read F1:13652

#### Sequence

DVLMTQLPLSLVPSLG

#### Sequence Length

16

#### Meta Information from PEAKS

##### Scan Identifier

F1:13652

##### Original Sequence (length=24)

D

+58.01

V

L

M

T

Q

L

P

L

S

L

V

P

S

L

G

##### Posttranslational Modifications

Carboxymethyl (KW X@N-term)

##### Source File

20191211\_F1\_Ag5\_peng0013\_SA\_Flag\_Asp\_N.raw

##### Fraction

1

##### Scan Feature

F1:17461

##### De Novo Score

93

##### Confidence score

93

##### Mass Charge Ratio

870.9761

##### Mass

1739.938

##### Charge

2

##### Retention Time

76.6

##### Predicted Retention Time

-

##### Area

3824500

##### Fragmentation Mode

ETHCD
