## Supplementary material for "Template-based assembly of proteomic short reads for *de novo* antibody sequencing and repertoire profiling": F1_14229.html

Details F1\_14229

OverviewUndefined

### Read F1:14229

#### Sequence

DAVVCFLNNFYPK

#### Sequence Length

13

#### Meta Information from PEAKS

##### Scan Identifier

F1:14229

##### Original Sequence (length=21)

D

A

V

V

C

+58.01

F

L

N

N

F

Y

P

K

##### Posttranslational Modifications

Carboxymethyl

##### Source File

20191211\_F1\_Ag5\_peng0013\_SA\_Flag\_Asp\_N.raw

##### Fraction

1

##### Scan Feature

F1:15221

##### De Novo Score

96

##### Confidence score

96

##### Mass Charge Ratio

794.3789

##### Mass

1586.7439

##### Charge

2

##### Retention Time

79.43

##### Predicted Retention Time

-

##### Area

86337

##### Fragmentation Mode

HCD
