## Supplementary material for "Template-based assembly of proteomic short reads for *de novo* antibody sequencing and repertoire profiling": F1_14239.html

Details F1\_14239

OverviewUndefined

### Read F1:14239

#### Sequence

DDPEVQFSWFV

#### Sequence Length

11

#### Meta Information from PEAKS

##### Scan Identifier

F1:14239

##### Original Sequence (length=11)

D

D

P

E

V

Q

F

S

W

F

V

##### Posttranslational Modifications

##### Source File

20191211\_F1\_Ag5\_peng0013\_SA\_Flag\_Asp\_N.raw

##### Fraction

1

##### Scan Feature

F1:11465

##### De Novo Score

99

##### Confidence score

99

##### Mass Charge Ratio

684.8092

##### Mass

1367.6033

##### Charge

2

##### Retention Time

79.54

##### Predicted Retention Time

-

##### Area

761160

##### Parts Per Million

0.4

##### Fragmentation Mode

ETHCD

##### Also found in scans

F1:12643 F1:11889 F1:12992 F1:13241 F1:13305 F1:12590 F1:12135 F1:11827 F1:13360
