## Supplementary material for "Template-based assembly of proteomic short reads for *de novo* antibody sequencing and repertoire profiling": F1_14240.html

Details F1\_14240

OverviewUndefined

### Read F1:14240

#### Sequence

DVVAFLNNFYPK

#### Sequence Length

12

#### Meta Information from PEAKS

##### Scan Identifier

F1:14240

##### Original Sequence (length=12)

D

V

V

A

F

L

N

N

F

Y

P

K

##### Posttranslational Modifications

##### Source File

20191211\_F1\_Ag5\_peng0013\_SA\_Flag\_Asp\_N.raw

##### Fraction

1

##### Scan Feature

F1:12554

##### De Novo Score

98

##### Confidence score

98

##### Mass Charge Ratio

713.8724

##### Mass

1425.7292

##### Charge

2

##### Retention Time

79.53

##### Predicted Retention Time

-

##### Area

816100

##### Parts Per Million

0.8

##### Fragmentation Mode

ETHCD
