## Supplementary figures and images for "Template-based assembly of proteomic short reads for *de novo* antibody sequencing and repertoire profiling"

### export_pdf_example.png

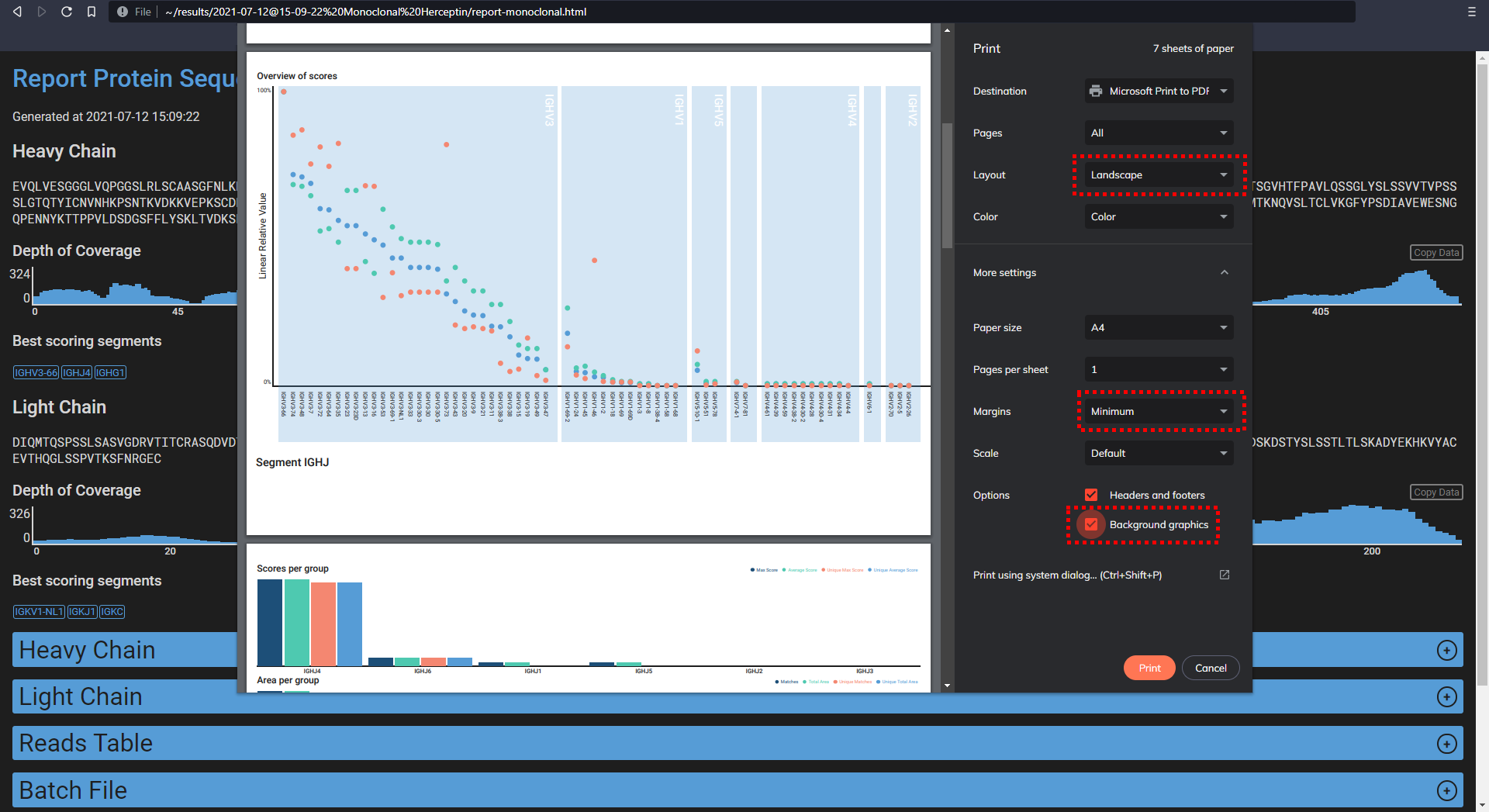
