## Supplementary Information for "Template-based assembly of proteomic short reads for *de novo* antibody sequencing and repertoire profiling"

Supplementary Data legend

Supplementary figures S1-S2

**Supplementary Data.**

All reported STITCH analyses are provided with the complete output reports. The supplementary data can be unzipped to browse the interactive HTML reports. The corresponding analysis parameters are provided under the ‘Batch File’ menu in each report.

```
A) Herceptin

template    ...DTAVYYCAR                  YFDYWGQGTLVTVSS...
de novo        DTAVYYCSRWGG     SRWGGDGFYAMDYWGQGTLVTVSS

overlap        DTAVYYCSRWGG
                      SRWGGDGFYAMDYWGQGTLVTVSS

ref         ...DTAVYYCSRWGGDGFYAMDYWGQGTLVTVSS...


B) anti-FLAG-M2

template    ...DSAVYYCAR                      YFDYWGQGTTLTVSS...
de novo        DSAVYYCAREKFYGY     VYYCAREKFYGYDYWGQGATLTVSS

overlap        DSAVYYCAREKFYGY
                  VYYCAREKFYGYDYWGQGATLTVSS

ref         ...DSAVYYCAREKFYGYDYWGQGATLTVSS...


C) F59

template    ...DTALYYCAKD                     NWFDSWGQGTLVTVSS...
de novo            YYCAKDVRPYYD                 FDSWGRGTPVTVSS

overlap        DTAFYYCAKDVRPYYDF
                        DVRPYYDFFN
                             YDFWAFDSWGR
                                  FDSWGRGTPVTVSS

ref         ...DTAFYYCAKDVRPYYDFWAFDSWGRGTPVTVSS...
```

**Supplementary Figure S1.** Detailed view of CDRH3 reconstruction by STITCH of the antibodies shown in main text figure 2. Shown are the selected template sequences, aligned reads, found overlap and known reference sequence.

```
>Patient 1 (I: 89.1% S: 92.7% G: 0.0%)
de novo   | QTVLGQMPSSSGSPGQSVTISCTGTSSDVGGYNYVSWYQQHPGKAPKLMIYDVTKRPSGVPDRFSGSKSGNTASLTLSGLQAEDEADYYCCSYAGLDLFVLFGGGTKLTV
reference | GPDLTQPRSVSGSPGQSVTLSCTGTSSDVGGYNYVSWYQQHPGKAPKLMIYDVTKRPSGVPDRFSGSKSGTTASLTISGLQAEDEADYYCCSYAGIDIFVLFGGGTKLTV
            *** * ** *         *                                                  *     *                  * *

>Patient 5 (I: 95.3% S: 96.3% G: 0.0%) & (I: 91.6% S: 96.3% G: 0.0%)
de novo     | EIVLTQSPDTLSLSPGERATLSCRASQSVSSSYLAWYQQKPGQAPRLLIYDASTRATGIPDRFSGSGSGTDFTLTISRLEPEDFAVYYCQQYGRSPYTFGPGTKVDI
reference A | EIVLTQSPGTLSLSPGERATLSCRASQSVSSSYLAWYQQKPGQAPRLLIYDASTRATGIPDRFSGSGSGADFLLTISSLEPEDFAMYYCQQYGRSPYTFGPGTKVDI (5A)
reference B | EIVLSQSPDTLSLSPGERATLSCRADKSVSSNYVAWYQQKPGQAPRLLIYDAFTRATGIPDRFSGSGSETDYTLTISTLEPEDFAVYYCQQYGRSPYTFGPGTKVDI (5B)
                  *   *                **    * *                  *               ** **    *       *

>Patient 6 (I: 99.1% S: 99.1% G: 0.0%)
de novo   | DIQMTQSPSSLSASVGDRVSITCRASESISSYVNWYQQKPGKAPKLLIYTASSLQSGVPPRFSGSGSGTDFTLTISSLQPEDFATYYCQQSYSTPITFGQGTRLEI
reference | DIQMTQSPSSLSASVGDRVSITCRASESISSYVNWYQQKPGKAPKLLIYTASSLQSGVPPRFSGSASGTDFTLTISSLQPEDFATYYCQQSYSTPITFGQGTRLEI
                                                                             *

>Patient 7 (I: 93.4% S: 97.2% G: 0.9%)
de novo   | DIQMTQSPSSLSASVGDRVTITCQASQDIAKYLNWYQQKPGKPPKLLIYDTSNLETGVPSRFSGSGSGTDFTLTISSLQPEDIATYYCQQYDDFPLTFGPGTKVDI
reference | DIQMTQSPSSLSASVGDRVTITCQASQDLAKYLNWYQQKPGKPPKLLIYDTSNLETGVPSRFSNG-GGTDFTFTINSLQPEDLATYYCQQYDDFPLTFGPGTKVDI
                                        *                                  ****     *  *      *

>Patient 8 (I: 99.1% S: 100.0% G: 0.0%)
de novo   | DIQMTQSPSTLSASVGDRVTITCRASQSISSSLAWYQQKPGKAPKLLIYDASSLETGVPSRFSGSGSGTEFTLSISSLQPDDFATYYCQHYNSYSLTFGQGTKVEI
reference | DIQMTQSPSTLSASVGDRVTITCRASQSLSSSLAWYQQKPGKAPKLLIYDASSLETGVPSRFSGSGSGTEFTLSISSLQPDDFATYYCQHYNSYSLTFGQGTKVEI
                                        *

>Patient 13 (I: 89.2% S: 95.5% G: 0.0%)
de novo   | QSVLTQPPSVSGAPGQRVTISCTGSSSNIGAGWDVHWYQQLPGTVPKLLIYADRNRPSGVPERFSGSGSGTDFTLTISGLQAEDEADYYCQSYDSALSGFYVFGTGTKVLV
reference | EAPLTQPPSVSGAPGQRVTLSCTGSSSNLGAGWDVHWYQQLPGTVPKLLIYADRNRPSGVPERFSGSKSGTSATVAIAGLQAEDEADYYCQSYDSALSGFYVFGTGTKVIV
            ***                *        *                                      *   ** ** *                               *

>Patient 15 (I: 97.2% S: 99.1% G: 0.0%)
de novo   | DIQMTQSPSTLSASVGDAVTITCRASQSINVWLAWYQQKPGKPPKLLIYEASNLESGVPSRFSGSGSGTEFTLTISRLEPDDFATYYCQQYNSYPYTFGQGAKLEI
reference | DIQMTQSPSTLSASVGDAVTITCRASQSLNVWLAWYQQKPGKPPKLLIYEASNLESGVPSRFSGSGSGTEFTLTISSLQPDDFATYYCQQYNSYPYTFGQGAKLEI
                                        *                                        *

>Patient 18 (I: 98.1% S: 99.1% G: 0.0%)
de novo   | DIQMTQSPSSLSASVGDRVTITCQASRDISNYLNWYQQKPGKAPMLLIYAASNLQTGVPSRFSGSGSGTDFTLTISSLQPEDIATYYCQQYGNLPLTFGGGTKVEI
reference | DIQMTQSPSSLSASVGDRVTITCQASRDISNYLNWYQQKPGKAPMLLIYAASNLQTGVPSRFSGSGSGTDFTFTISSLQPEDLATYYCQQYGNLPLTFGGGTKVEI
                                                                                    *         *

>Patient 19 (I: 99.1% S: 100.0% G: 0.0%)
de novo   | DIQMTQSPSSLSASVGDRVTITCQASQDIGNYLNWYQQKPGKAPRLLIYDASDLEEGVPSRFSGSGSGTDFTFTISSLQPEDFATYYCQQYHTLPPLTFGGGTKVDV
reference | DIQMTQSPSSLSASVGDRVTITCQASQDLGNYLNWYQQKPGKAPRLLIYDASDLEEGVPSRFSGSGSGTDFTFTISSLQPEDFATYYCQQYHTLPPLTFGGGTKVDV
                                        *

>Patient 20 (I: 98.1% S: 99.1% G: 0.0%)
de novo   | DIQMTQSPSTLSTSVGDRVTITCRASQSIRTWLAWYQQKPGKAPKLLIYDASTLETGVPSRFSGSGSGTDFTLTISSLQPEDFATYYCQQYNDYSGTFGQGTKLEI
reference | DIQMTQSPSTLSTSVGDRVTITCRASQSIRTWLAWYQQKPGKAPKLLIYKASTLETGVPSRFSGSGSGTEFTLTISSLQPEDFATYYCQQYNDYSGTFGQGTKLEI
                                                             *                   *
```

**Supplementary Figure S2.** STITCH reconstruction of light chains from multiple myeloma patient urine (Chamot-Rooke and colleagues, ref 43). CDRs are annotated, sequence conflicts highlighted by an asterisk (*) and sequence identity, similarity and gaps indicated in parentheses.
